# Testing genomic offset with common gardens in genetically structured black spruce (*Picea mariana)*

**DOI:** 10.1101/2025.10.30.685617

**Authors:** Anna Fijarczyk, Etienne Robert, Benjamin Marquis, Manuel Lamothe, Patrick Lenz, Martin P. Girardin, Nathalie Isabel

**Affiliations:** Natural Resources Canada, Canadian Forest Service, Laurentian Forestry Centre, Québec, QC G1V 4C7, Canada; Institut des sciences de la forêt tempérée, Université du Québec en Outaouais, Ripon, QC J0V 1V0; Institut des sciences de l’environnement, Université du Québec à Montréal, Montréal, QC H3C 3P8, Canada; Natural Resources Canada, Canadian Forest Service, Great Lakes Forestry Centre, Sault Ste. Marie, ON P6A 2E5, Canada; Centre for Forest Research, Université Laval, Quebec, QC G1V 0A6, Canada

**Keywords:** Black spruce, *Picea mariana*, genomic offset, Gradient Forest, genomic vulnerability, climate change

## Abstract

Boreal forests play a crucial role in regulating climate via storage and release of carbon. Anticipated changes in climate are expected to increase mortality and reduce biomass in many boreal tree species, putting at risk the functioning of this ecosystem and hence its role in carbon sequestration. Genomic offset methods leverage spatial distribution of genomic diversity and its association with environmental variables to predict population vulnerability to projected changes in climate. Here, we analyse over 60 populations and more than 1400 individuals of black spruce (*Picea mariana* (Mill.) B.S.P), a dominant boreal forest species, to compare population-level genomic offsets calculated using Gradient Forest and redundancy analysis (RDA) against multiple fitness traits measured in four long-term (>40 yr) common gardens. Within common gardens, we found that genomic offset projections were largely unaffected by the model choice, the number or type of markers used for model training, with the strongest discrepancies observed for LFMM climate-associated markers. Model performance remained relatively stable when the number or size of populations in the training set was reduced, suggesting that these models can reliably project genomic offsets for new populations. However, model performances varied among common gardens, with highly accurate fitness projections in some gardens but contradictory results in others. Model performance was influenced by the choice of climate predictors, their relationships with fitness traits, and the genetic cluster in which the models were evaluated. Overall, our results highlight the challenges of projecting genomic offsets across large spatial scales in genetically structured species, due to spatial variation in environmental drivers of adaptation and complex interactions among them. By capitalizing on our comprehensive validation, we identified the most robust models for projecting fitness declines in black spruce under future climate scenarios.

## Introduction

Boreal forests comprise 27% of the total forested area on Earth, covering more than 1 billion hectares across the northern hemisphere (GlobalFRA, n.d.). They are dominated by conifer species, including spruces (*Picea* spp.) and pines (*Pinus* spp.), that are adapted to short-growing seasons and cold temperatures. As one of the world’s largest terrestrial carbon reservoirs, these boreal forests play an important role in the regulation of global climate by storing and releasing carbon. Boreal forests have been under constant influence of natural disturbances such as extreme climate events, wildfires, insect and disease outbreaks since the retreat of the glaciers about 7000 ya (Brandt et al. 2013). With projected increase in global temperatures (IPCC 2023), many boreal regions are facing intensification of some of those challenges and increase of drought-related episodes in some parts of its distribution (Price et al. 2013), putting at risk long-term functioning of this ecosystem (Gauthier et al. 2015). It is becoming increasingly urgent to understand how boreal tree species will respond to these challenges, and test what are the potential benefits of assisted migration to species and tree populations at risk of maladaptation.

Ecological methods used to predict forest species dynamics in changing environments either model species distributions by examining shifts in the suitability of their habitats across the landscape (Prasad et al. 2020) or model biomass losses by examining species-specific functional traits such as tree mortality caused by drought, or migration failure (Aubin et al. 2018; Liu et al. 2023). These methods do not consider the evolutionary processes shaping intraspecific genetic variation across the landscape, which can lead to divergent populations’ responses to the same climate variables. Standing genetic diversity within populations has been shaped by past demographic events, such as local extinctions or expansions, by local selective pressures as well as by interactions with other species. These factors ultimately determine the ability of populations to respond to changing environments and can enhance the predictive accuracy of species distribution models (Urban et al. 2016; Aguirre-Liguori et al. 2021). Conifer tree species often display local adaptation to environmental conditions, as demonstrated with common garden experiments, where population-level functional traits often shift when growing conditions differ from those found at the seed source (Oleksyn et al. 1998; Lortie and Hierro 2022; Guo et al. 2022; Housset et al. 2018; Depardieu et al. 2020). Genomic studies further support this, revealing allele frequency changes in genes associated with climate-related traits (Evans et al. 2014; Depardieu et al. 2021; Bergmann 1978). An increasing number of studies considers population-level variation in phenotypes or genotypes to model species productivity predictions (Isaac-Renton et al. 2014; Patsiou et al. 2020; Bradley St Clair and Howe 2007; Robert et al. 2024).

Another group of predictive methods aims to estimate genotype–environment mismatch (genomic offset) under climate change using species-wide genomic variation. Approaches such as Gradient Forest (Fitzpatrick and Keller 2015; Ellis et al. 2012), Risk of Non-Adaptedness (Rellstab et al. 2016), Latent Factor Mixed Models (Gain and François 2021), or Redundancy Analysis (Capblancq and Forester 2021) model linear or non-linear relationships between genetic variants and environmental variables across the landscape. These models are then used to project genomic offset under future climate scenarios. Genomic offset is typically calculated as the Euclidean distance between the genomic composition of a genotype under current conditions and its projected composition under new conditions. Theoretically, this metric has been linked to changes in fitness when a polygenic trait is under stabilizing selection toward an optimal phenotype (Gain et al. 2023; Capblancq et al. 2025). Such methods come with their own limitations; for instance, the underlying assumption is that genotypes are most fit in their local climate, and that maladaptation to new conditions is instantaneous. Therefore, even though these methods hold promise for straightforward and phenotype-free risk assessment of species to changing climate (Rellstab et al. 2021), validation studies are needed to assess their utility for conservation or management purposes in different contexts.

To validate genomic offset methods, several studies have conducted either simulations (Lind and Lotterhos 2025b; Láruson et al. 2022; Lind and Lotterhos 2025a; Gain et al. 2023) or empirical validation in common gardens in which anticipated climate change in some future period is substituted by different climate conditions experienced by the genotype transplanted to a common garden experiment (Fitzpatrick et al. 2021; Capblancq and Forester 2021; Lind et al. 2024; Rhoné et al. 2020; Lachmuth et al. 2024; Archambeau et al. 2025; Verrico et al. 2025; Gain et al. 2023). The consensus across these studies is that genomic offset is often correlated with fitness and often predicts fitness better than climate or geographical distance alone. However, different models can yield inconsistent patterns of maladaptation (Lind et al. 2024) and the strength of these correlations varies depending on the studied species, as well as data-and common garden-specific factors (Fitzpatrick et al. 2025). Simulation studies demonstrated that the accuracy of genomic offset improves when sampling is denser along environmental gradients (Láruson et al. 2022), and when selection is strong (Lind and Lotterhos 2025b). On the other hand, differences in effective population sizes (Láruson et al. 2022), the presence of novel environments, and the inclusion of non-causal environmental variables can all reduce the accuracy of genomic offset projections (Lind and Lotterhos 2025b). Finally, the choice of markers, whether random, neutral, or adaptive, has been shown to have either a limited impact on genomic offset estimates (Lind and Lotterhos 2025b; Lind et al. 2024; Láruson et al. 2022; Fitzpatrick et al. 2021; Lachmuth et al. 2023) or in some cases, to improve the accuracy (Gain et al. 2023). These findings highlight the importance of investigating the limitations of genomic offset across different species. Key unresolved questions include the role of genetic structure in shaping the variability of genomic offset patterns, the identification of fitness traits that best capture local adaptation, and the influence of environmental variable selection, and sampling scheme, on projection accuracy.

Our study focuses on black spruce, a widespread boreal species extending from Alaska to Newfoundland and Labrador (Burns and Honkala 1990). This dominant component of Canadian boreal forests displays adaptation to variable soil and climate conditions, ranging from poorly drained bogs to the northern treeline in northwestern Canada, where it can be found in cold, dry climates and on permafrost-underlain soils. Historically, black spruce has reached its current range from at least two to three distinct glacial refugia situated in the south after the retreat of the Laurentide ice sheet (Gérardi et al. 2010; Jaramillo-Correa et al. 2004). Studies using mitochondrial, cytoplasmic and nuclear markers have identified three main genetic clusters, roughly corresponding to the western, central and eastern regions of Canada (Gérardi et al. 2010; Prunier et al. 2012; Girardin et al. 2021). These findings suggest that current distribution resulted from multiple expansion events from southern refugia and indicate that the species range is composed of partially independent genetic pools within which climate adaptation occurs (Prunier et al. 2012).

The impact of climate on the growth of black spruce has been extensively studied, particularly using dendroecological data. The interplay between temperature and precipitation determines water availability, making drought stress one of the most important factors limiting growth and productivity across the species range (Girardin et al. 2024). Compared to other species such as jack pine, black spruce generally exhibits poor recovery following drought stress (Marchand et al. 2021, 2025). The western and central regions of the species distribution are characterised by climates where the ratio of precipitation to evaporative demand is relatively low, constraining tree growth (Price et al. 2013; Hogg et al. 2013). Although higher temperatures can enhance growth in some northwestern populations, they more often exacerbate the drought conditions and have negative effects on growth (Walker et al. 2015; Sniderhan et al. 2021). In the more humid eastern portion of the range, growth is mostly limited by cold temperatures in the north, whereas in the south, warmer conditions can increase water availability constraints and respiration costs (Girardin et al. 2016).

Taken together, forecasts very broadly predict that increasing temperatures and droughts will lead to higher vulnerability especially of western and central North American black spruce populations where precipitation is the limiting factor (Lesven et al. 2024). While northeastern populations can potentially experience growth increase in the near future, this effect is expected to be transient (D’Orangeville et al. 2018). So far, only a few studies have explicitly considered genetic variation of black spruce, focusing on modelling productivity and resistance to extreme drought events in the context of assisted migration along multiple climate gradients (Girardin et al. 2021; Robert et al. 2024, 2025). These studies showed that genetic divergence between black spruce genetic clusters influences phenotypic response to climate that can accumulate over time (Girardin et al. 2021; Robert et al. 2024). Models suggest that northernmost populations may experience a transient increase in growth due to reduced cold limitation. However, over time, all populations are projected to exhibit declines in biomass, with the western cluster showing lower growth relative to the central and eastern clusters (Robert et al. 2024). It is important to note though that other climate mediated factors can exacerbate or counteract these projections. This includes outbreaks of insects such as eastern spruce budworm (Bellemin-Noël et al. 2021), reduced post-fire recovery (Baltzer et al. 2021), loss of protective snow cover (Marquis et al. 2022), shifts in tree species composition, which can alter interspecific competition dynamics and reshape soil symbiotic communities. In addition, changes in bud phenology, where higher temperatures can lead to earlier bud break in the spring, may increase the risk of frost damage (Mura et al. 2022; Rossi 2015; Guo et al. 2022). While this study focuses specifically on direct effects of climate, common garden experiments in black spruce provide an ideal framework to assess how genome-wide variation can improve predictions of tree response to changing climatic conditions.

Here we examined >1,400 black spruce trees for genotypes and 17 phenotypic traits originating from 66 populations and grown across four common gardens. We leveraged nearly 30,000 DArTseq single nucleotide variants, in combination with climate and phenotypic information to validate genomic offset projections across the species distribution. Our main objectives were: i) to determine fine-scale genetic structure and distribution of genetic diversity across the distribution of black spruce, ii) to validate genomic offsets using a space-for-time substitution based on the phenotypes produced in the common garden experiment and iii) to project genomic offset of black spruce to future climate. Our extensive validation of genomic offset in a species with a pronounced genetic structure provides valuable insights for informing future forestry management strategies in the face of changing climate.

## Materials and Methods

### Sample Collection and DNA Extraction

Samples were collected between 2014 and 2019, from 70 provenances of black spruce, to which we refer to as populations throughout the text. A total of 2,237 specimens were collected, encompassing the species’ entire natural range in Canada and the USA (Fig. 1A, Suppl. Table 1, Suppl. Dataset 1). These samples were gathered from common gardens located in Chibougamau (CH, Quebec), Acadia (AC, New Brunswick), Mont-Laurier (ML, Quebec), Peace River (PR, Alberta), and Valcartier (VL, Quebec), all of which were initially established in 1974 or 1975 as part of the Range-Wide Provenance Study, with seeds sown in 1970 ((Morgenstern 1978); Suppl. Table 2). Only a small fraction of samples (n=37, nine of which remained after filtering) was collected from VL and used exclusively for genotyping. Additionally, we collected samples from 37 red spruce (*P. rubens* Sarg.) trees, from various populations within the allopatric zone of the species (Suppl. Dataset 2). Red spruce is known to hybridize with black spruce, we therefore included it in our analysis to account for genetic admixture between the two species.

**Figure 1.**
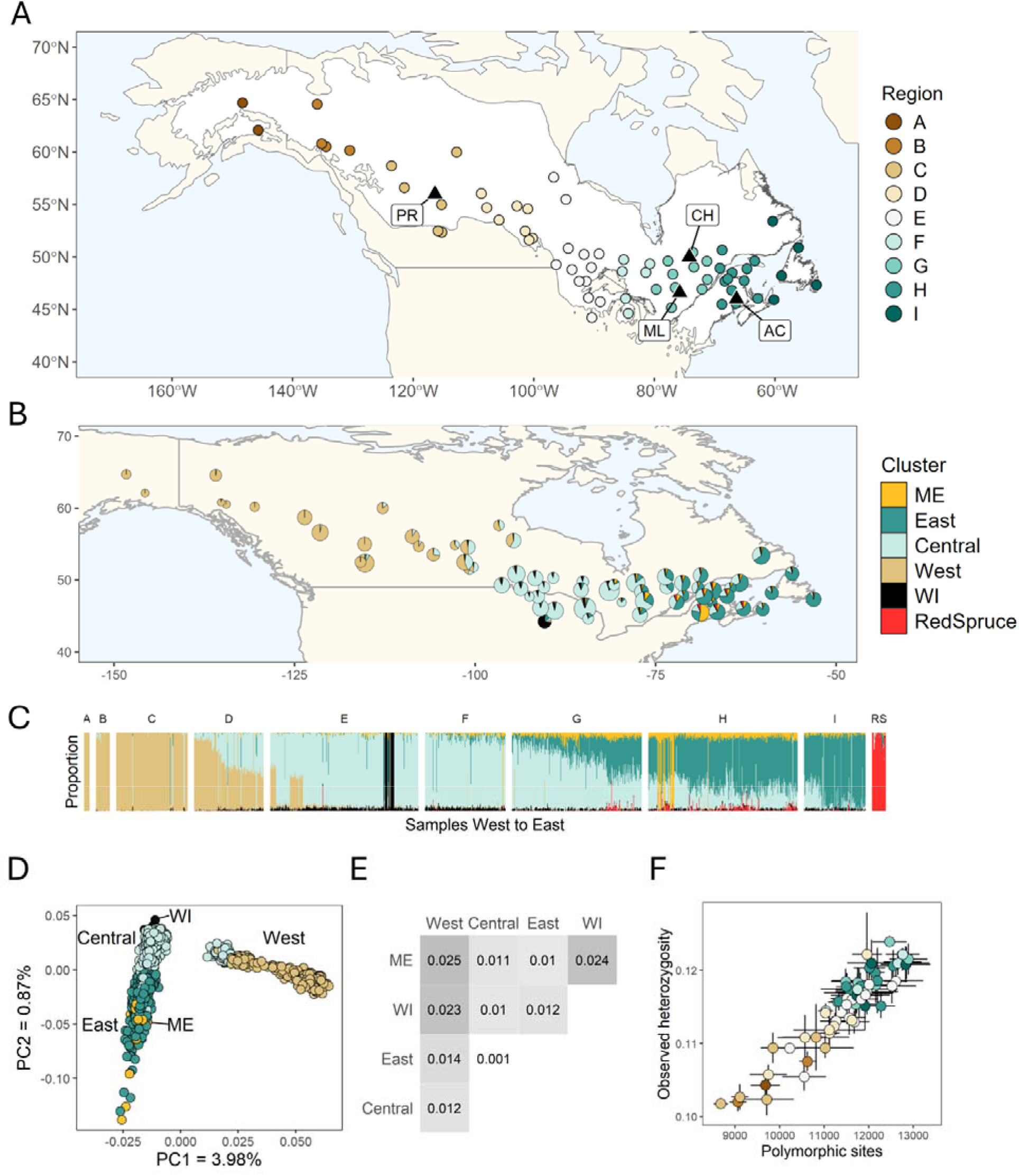
Sampling and population structure of black spruce. A. Sampling and location of common gardens. Dots indicate population locations, colored by geographic regions delineated at equal intervals along the longitudinal axis. Black triangles indicate the location of common gardens (PR - Peace River, ML, Mont-Laurier, CH - Chibougamau, AC - Acadia). White area shows the distribution of black spruce. B. Pie charts show mean genetic ancestry proportions per population, with pie size proportional to population sample size. C. Admixture analysis showing ancestry proportions of samples at K=6, including red spruce (RS). D. Principal component analysis computed on 29,972 biallelic SNPs. Colors correspond to genetic clusters depicted in panel B. E. Pairwise *F*_ST_ matrix among five genetic clusters of black spruce. F. Observed heterozygosity and number of polymorphic sites across populations. Populations are colored by geographical regions depicted in panel A.

The DNA required for sequencing was extracted from 30-50 mg of either frozen needle tissue (for the CH, ML and VL sites) or cambial tissue (for AC and PR sites) with a Nucleospin 96 Plant II kit (Macherey-Nagel, Bethlehem, PA) using the centrifugation processing protocol with a cell lysis step with PL2 buffer for 1h at 65°C. Cambium (with phelloderm) tissue was obtained from bark samples collected with a 1 cm diameter punch sterilized with 70% alcohol between each sample. Prior to extraction, all tissues were ground to powder with a Mixer Mill MM300 (Retsch GmbH, Haan, Germany) after being plunged into liquid nitrogen for two minutes (this process was repeated twice).

### Sequencing and Genotyping

For single nucleotide polymorphism (SNPs) discovery, we employed the DArTseq™ method, which shares similarities with genotyping-by-sequencing (GBS) but includes a complexity reduction step targeting low-copy sequences within the genome. Our procedure involved digesting genomic DNA with PstI and MseI restriction enzymes, ligating barcoded adapters, amplifying the resulting products via PCR, and sequencing them using a HiSeq 2500 system. This entire process was conducted by Diversity Arrays Technology, headquartered in Bruce, Australia, as described by Kilian et al. (2012). Our SNP analysis focused exclusively on biallelic SNPs for subsequent investigation. The DNA plates containing extracted samples were sent and processed in three consecutive years: 2018, 2019, and 2020, referred to herein as batches 1, 2, and 3 (Suppl. Dataset 1). Genotypes were generated separately for each of these three batches, and at the conclusion of the project, all sequences produced were merged and clustered together to create the final dataset for analysis.

## Data processing

### Genetic data

SNPs were generated from ∼70 nt sequence clusters by setting different sequence identity thresholds producing four independent sets of variants. After checking SNP genotyping rates and ensuring no overlap among sequence tags from the four files, SNP sets resulting from the three lowest clustering thresholds were combined. SNPs were filtered separately for the black spruce and red spruce samples. We filtered out variants with more than 50% of missing genotypes, heterozygosity above 50% and the minor allele frequency (MAF) below 1%. We ran admixture analysis to determine population structure and at the same time identify potential mislabelled samples. The number of genetic clusters was determined with ADMIXTURE v1.3.0 (Alexander et al. 2009) using all black and red spruce individuals. First genotypes were saved to plink format using dartR package v2.9.7 (Mijangos et al. 2022). Genetic ancestries were assigned for the K number of clusters, where K ranged between 2 and 15. 5-fold cross-validation error was estimated with each run. The cross-validation error curve flattened starting with K=4, pointing to three main black spruce genetic clusters (West, Central, East) and one red spruce cluster. Several samples showed assignment to groups from distant clusters, implying potential mislabelling of samples. For the final dataset we removed most dubious samples, including 5 samples in the region A and B (Alaska) assigned to the Central or East genetic cluster, and 1 sample from the region I (Newfoundland) assigned to the West cluster. In addition, we removed samples with a call rate below 85% retaining 1467 black spruce samples from 66 populations (mean of 22 samples per population, range 1-49). A dataset of unique black spruce variants consisted of 29,972 biallelic SNPs.

### Climate data

We used the BioSIM v11 (Fortin et al. 2022) to simulate daily weather at the populations’ origins and that of the common gardens for the period 1961-1990. We then calculated 43 climate variables representing temperature, precipitation and the interplay between them–such as climate moisture index (CMI)–at the annual or seasonal scale (see Suppl. Table 3, Suppl. Dataset 3 for a complete list of variables and the climate aggregate over the full period for all populations and 5 common gardens). We obtained simulated daily climate data between 2021 and 2100 for the three emission scenarios (shared socio-economic pathways): SSP2-4.5, SSP3-7.0 and SSP5-8.5, using an ensemble of 13 climate models from the Ouranos ESPO-G6-R2v1.0.0 dataset (Lavoie et al. 2024). Annual and seasonal climate variables corresponding to a subset of those obtained for the past period (a total of 23) were calculated from daily climate data. Mean annual CMI for the future period was obtained by importing daily variables to BioSIM. To ensure that variables obtained with BioSIM and those estimated from PAVICS datasets were comparable, we visually inspected the temporal distributions of annual climate means per population to confirm that means calculated for past and future periods followed a continuous trend and retained 14 of them (CMI, mayMinT, TP, MAT, MMinT, MMaxT, sumTP, WSTP, winMT, sprMT, sumMT, fallMT, MCMT, MWMT). For clarification, we refer to these 14 climate variables, collectively with photoperiod (dPP), as the “small set of climate variables”.

### Phenotypic data

Population means and standard errors of all traits used as a proxy for fitness are given in Suppl. Dataset 4. Accumulated biomass and wood density (PR: n=397, ML: n=612, CH: n=507, AC: n=481) were obtained from wood cores as described in Robert et al. (2024). Tree total aboveground accumulated biomass (kg) was calculated as the sum of annual biomass between years 1980 and 2015 (ABtot), in addition to eight non-overlapping five-year intervals between 1980 and 2020 (AB5). Wood density was reported as the average wood density for the ring from the year 2015. Height in cm and diameter at breast height (DBH) in cm, were measured in 2019 (PR: n=400), 2015 (ML: n=667), 2016 (CH: n=451), and 2017 (AC: n=492). Survival was reported as the % of surviving trees per population in 2011 (PR: n=26), 2007 (ML: n=42), 2006 (CH: n=38), and 2003 (AC: n=40). Four drought-related indices to extreme drought events (*sensu* Lloret et al. (2011)) were calculated from the wood cores for three drought years at AC, two at CH, five at ML and four at PR, and averaged for each site (Robert *et al*. in prep). These indices included resistance (Rt)–growth during the year of the drought over the average growth of the two years prior to the drought (higher ratio means higher resistance); recovery (Rc)–the average growth of the two years after the drought over the growth during the year of the drought (higher ratio means higher recovery); resilience (Rs)–the ratio of growth after the drought event over the growth before the event (higher ratio means higher resilience); and relative resilience (Rr)–resilience minus resistance (higher ratio means higher relative resilience).

To obtain population-level trait estimates while accounting for the experimental design of the common gardens, we fitted the mixed model to each trait except for survival:

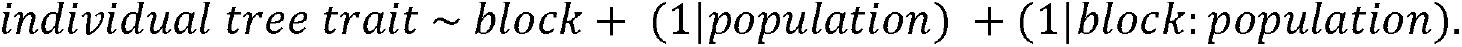

We removed the estimated block effect and the BLUP (best linear unbiased prediction) associated with the population ✕ block interaction from each individual observation. This procedure removes variation due to differences among blocks and to population-specific within-block deviations, while retaining variation among populations. The corrected values were then averaged for each population within each common garden. ABtot and AB5 were first log transformed. Blocks with less than 10 individuals were excluded.

### Population structure analyses

Population structure was determined using an ADMIXTURE program, as described above, and principal component analysis was used to visualize ordination of genetic clusters. Genotypes of the black spruce individuals were saved to gds format using dartR package v2.9.9.5 (Mijangos et al. 2022). PCA was run using package SNPRelate v1.45.0 (Zheng et al. 2012).

To investigate spatial structure, we retained 63 black spruce populations with at least five individuals. Weir and Cockerham estimate of *F*_ST_ between pairs of populations was calculated with assigner v0.7.0 (Gosselin et al. 2019) on a random subset of 1000 SNPs. In this analysis, genotypes were transformed to a tidy format using radiator v1.4.0 (Gosselin 2020). Geographic distances (km) between populations, were derived from spatial coordinates converted to a spatial sf object using a WGS 84 coordinate reference system from sf v1.1-0 (Pebesma 2018). To test for isolation by distance, a Mantel test was performed by correlating matrices of genetic distances (*F*_ST_ / (1 - *F*_ST_)) and geographic distances with package vegan v2.7-3 (Oksanen et al. 2025), with 9999 permutations, and the Spearman’s rank correlation.

In parallel, spatial autocorrelation and the relationship between genetic and geographic structure were assessed using Moran’s eigenvector maps (MEMs). First, a spatial weighting matrix was created for populations with parameters “type = 6, k = 8” using adegenet v2.1.11 (Jombart 2008) and spdep v1.4-2 (Bivand and Wong 2018). MEMs were then computed with package adespatial v0.3-29 (Guénard and Legendre 2022).

For spatial PCA (sPCA), we estimated allele frequencies within populations with dartR package v2.9.9.5 (Mijangos et al. 2022) and removed SNPs with missing values in any of the populations. sPCA was run with package adegenet v2.1.11 (Jombart 2008), using a spatial weighting matrix estimated above. Finally, Moran’s coefficient was calculated with package adespatial v0.3-29 (Guénard and Legendre 2022) to assess whether genetic variation captured by the principal components is spatially autocorrelated.

Pairwise *F*_ST_s between clusters were estimated in the same way as between the populations. Individuals were assigned to one of the five clusters identified with admixture with the number of clusters K=6 excluding red spruce (West, Central, East, WI and ME). Each individual was assigned to a cluster based on the dominant ancestry assignment. Analysis of molecular variance (AMOVA) was conducted at three levels: cluster, population and individual. Only populations with at least 5 individuals were retained. AMOVA was estimated with poppr v2.9.8 (Kamvar et al. 2014). Heterozygosity was calculated individually for each population, using the original dataset only filtered for variants not sequenced in more than 50% of samples, but with rare alleles retained. We focused on 60 populations with seven or more trees and randomly drew seven genotypes five times. For each subset, we removed any missing and monomorphic sites and calculated observed heterozygosity as the number of heterozygous SNPs for each individual divided by all sites and averaged across samples in each population. Expected heterozygosity was calculated for each population as the sum of squared frequency of allele 1 and 2 in each site, averaged across sites and subtracted from 1.

### Spatial associations between genotypes and climate

To visualize clustering of samples by climate, we ran PCA on 43 scaled and centered climate variables using the prcomp function from the base R package stats. To estimate how much distribution of genetic diversity is correlated with climate we conducted partial redundancy analysis (pRDA) by fitting three main climate principal components (PCs), three main Moran eigenvectors (MEMs), and two main PCs from genetic diversity ordination on a matrix of population allele frequencies. pRDA models were fitted to determine the impact of genetic, spatial and climate explanatory variables, one class (spatial, genetic, climate) at a time while controlling for the remaining variables. Models’ adjusted *R*^2^ were estimated and the significance of correlation of each class of variables with genetic variation was tested with a permutation test. In addition, pRDA was conducted at the genetic cluster level, after prior recalculation of two main PCs from genetic diversity ordination and three main MEMs. All functions are found in the vegan package v2.7-3 (Oksanen et al. 2025).

To determine which climate variables are best associated with genetic variation we ran Gradient Forest (Ellis et al. 2012; Fitzpatrick and Keller 2015) using 43 climate variables and 3 MEMs depicting spatial patterns of distribution. First, we used the allele frequency matrix generated in previous steps for 63 populations, removed any non-variant markers and selected a random subset of 1000 markers. We fitted the model using the gradientForest package v0.1-37 (Ellis et al. 2012), with 500 bootstrapped trees and correlation threshold of 0.5. Variables were ranked based on the mean raw accuracy importance.

### Associations between phenotypes and climate

To investigate local adaptation to climate, a Spearman’s rank correlation was calculated between phenotypic traits and climate transfer distance while accounting for spatial autocorrelation using modified t test from the R package SpatialPack v0.4-1 (Vallejos et al. 2020). *P*-values were corrected for multiple testing with the Benjamini-Hochberg procedure at a false discovery rate of 0.05. Climate transfer distance was calculated as the Euclidean distance of each population to its origin in climate space defined by four major PC axes of a Principal Component Analysis based on 43 climate variables. To calculate correlations, populations with less than four trait values were removed.

### Testing genomic offset with common gardens

We used common garden experiments to test the accuracy of genomic offset projections. In each common garden, models were trained using population-specific climate variables and allele frequencies, and genomic offsets were then projected to climatic conditions of the common gardens (Fig. 3). As a measure of model performance (or genomic offset projection accuracy), we calculated Spearman’s rank correlation between genomic offsets, that serve as proxies of population maladaptation to common garden conditions, and population-level phenotypic trait means, that serve as proxies for fitness. More negative correlation coefficients were interpreted as indicating better model performance.

**Figure 2.**
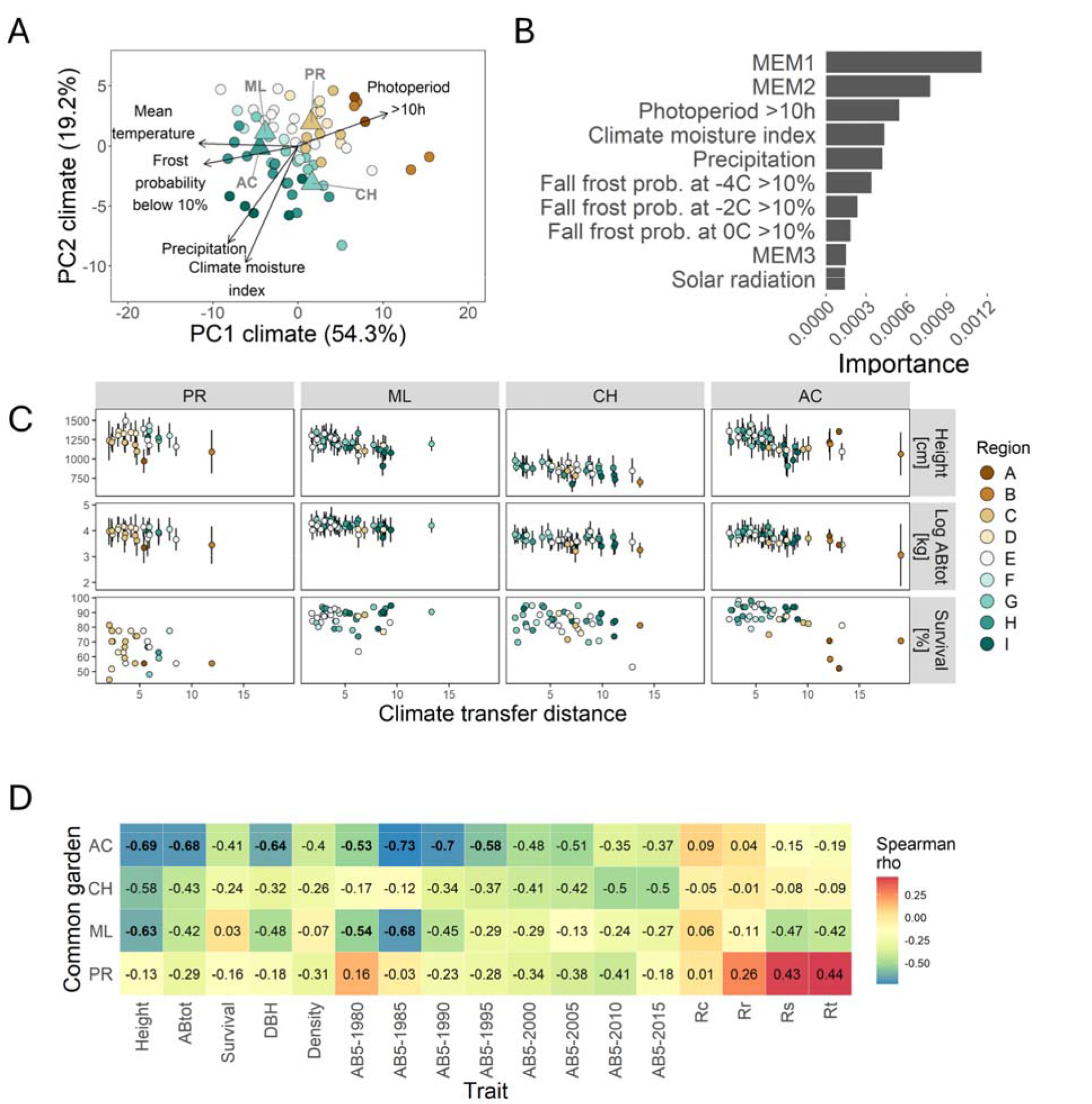
Local adaptation of black spruce to climate. A. Ordination of populations based on 43 climate variables. Vectors representing five climate variables identified using Gradient Forest are shown as arrows, with lengths scaled by a factor of 12. Locations of four common gardens (PR, ML, CH, AC) are shown by triangles. B. Importance ranking of the top climate and spatial variables associated with genetic variation across species distribution. C. Population means with standard deviation of height, accumulated biomass and survival rate plotted against climate transfer distance, defined as the Euclidean distance between the climate of origin of each population and that of the common garden site. D. Rank correlation coefficients between population means of traits and climate transfer distance. Bold values indicate coefficients that are statistically significant (*P* <0.05) after multiple testing correction. ABtot - total accumulated biomass, AB5 - five-year accumulated biomass, Rc - recovery, Rr - relative resilience, Rs - resilience, Rt - resistance. The year following AB5-labels indicates the beginning of the five-year period over which accumulated biomass was estimated. Colors in panels A and C correspond to geographic regions shown in Fig. 1A.

**Figure 3.**
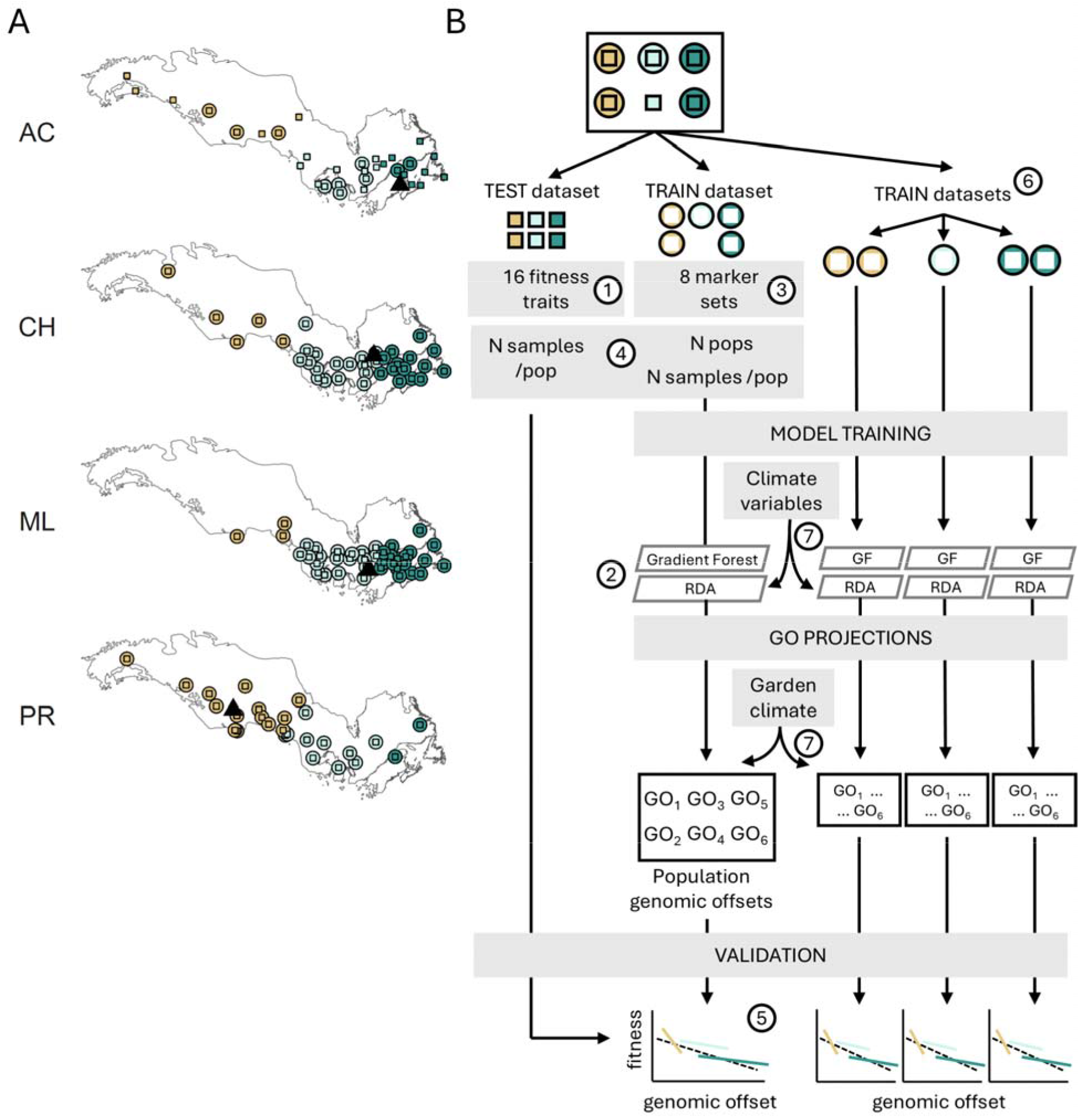
Validation framework for genomic offset projections. A. Sampling scheme showing populations used for model training (circles) and testing (squares) within each common garden. Populations were included in both datasets, when possible, except when genotype data were unavailable (e.g. AC). Triangles indicate common garden locations. B. Workflow used to validate genomic offset projections. For each common garden, samples were partitioned into training and test datasets. Models (Gradient Forest and RDA) were trained using population allele frequencies and climate variables, and genomic offsets (GO) were projected under common garden climate conditions. Model performance was evaluated by correlating genomic offset with population means fitness traits (Spearman rho), where more negative values indicate better predictive performance. Numbers indicate the seven factors evaluated: 1) fitness traits, 2) eight marker types, 3) two modelling approaches, 4) subsampling sensitivity, 5) population vs. cluster scale, 6) training at species vs. cluster level, 7) climate variable selection.

### Defining Training and Test Sets

To compare population genomic offsets with phenotypic traits, distinct sets of samples were used for model training (training dataset) and for phenotypic evaluation (test dataset), thereby avoiding bias associated with testing model accuracy on the same genotypes used for training (Fig. 3). Sample assignment to each dataset followed a structured procedure. Samples with only phenotypic data available, due to exclusion of their genotypes during quality filtering, were assigned to the test dataset, while all remaining samples were initially allocated to the training dataset. We then ensured that each population was represented by at least three individuals in the test dataset for reliable estimation of mean phenotypic traits, and at least five individuals in the training dataset for model fitting. When necessary, samples were transferred from the training dataset to the test dataset for a given population, provided that the training dataset retained at least five individuals for that population. In addition, two populations containing individuals assigned to the ME or WI genetic clusters were excluded from model validation to minimize potential confounding effects associated with genetically distinct regions. In the case of AC, many genotypes were missing due to low quality, resulting in test datasets that had more populations and individuals than the corresponding training datasets.

The selection of training and test datasets, as well as subsequent analyses, were performed separately for each trait, as the number of samples and populations varied among them. The numbers of populations and individuals per common garden and trait included in the training and test datasets are provided in the Suppl. Tables 4-5.

To investigate the behaviour of the genomic offset projection models, we explored seven aspects related to the input data and model training and their effects on model performance (Fig. 3).

#### 1) Common gardens and phenotypic traits

All models were trained separately for each common garden using its corresponding dataset (Fig. 3A). As proxies for fitness-related phenotypic traits in the test datasets, we used population means of 16 traits adjusted as described above, but averaged over test dataset samples only. The traits included: height, total accumulated tree biomass (ABtot), five-year accumulated biomass (AB5) over eight intervals, diameter at breast height (DBH), wood density, resistance (Rt), recovery (Rc), resilience (Rs) and relative resilience (Rr).

#### 2) Marker types

We tested the impact of eight different marker types on genomic offset projections, including genome-wide and climate-associated markers (single nucleotide polymorphisms). Genome-wide markers included all detectable markers (∼29K), a random subset of 100 (0.1K) or 1000 (1K) markers, as well as random subsets of 100 or 1000 markers not filtered for minor allele frequency (01.K-lf, 1K-lf). Climate-associated markers included outliers detected with LFMM, RDA and RDA with control for population structure (RDA-struct).

### Genome-wide markers

Genotype datasets were first prepared from the samples present in the training datasets, by calculating allele frequency per population separately for each common garden, while removing any invariant markers, markers with missing data or low frequency (minor allele frequency < 0.05). Low-frequency marker datasets (0.1K-lf and 1K-lf) were derived from the raw data and subjected to the same filtering pipeline as the standard marker sets, with the sole exception of the minor allele frequency thresholds.

### Climate-Associated Markers

We used a dataset with all detectable markers (∼29K) combined across common gardens to detect markers showing associations with distinct climate axes (precisely with the three main PC axes on 43 climate variables). Three tests were performed: LFMM, RDA, and RDA with correction for population structure. LFMM v2 was run with LEA v3.24 (Gain and François 2021), using K=3 latent factors (corresponding to three black spruce genetic clusters). *P*-values were calculated to test the significance of associations between each marker and individual effects of three climate PCs. *P*-values were then adjusted with Bonferroni multiple test correction. RDA was conducted with vegan v2.7-3 (Oksanen et al. 2025), first with the three climate PCs as only explanatory variables, and second by additionally conditioning for population structure using the two main axes from genetic variation ordination. Marker *P*-values were obtained by calculating Mahalanobis distances of each variant from the centroid of RDA marker loadings and comparing them with chi-squared distribution using the method recipe provided in Capblancq et al. (2018), and finally adjusting with Bonferroni multiple test correction. To estimate the amount of genetic variation represented by the selected list of outliers, observed heterozygosity was calculated. Heterozygosity per population was estimated on a combined set of all available genotypes from the four gardens excluding variants with minor allele frequency < 0.05.

### 3) Genomic offset models

#### Gradient Forest

Gradient Forest was run using the R package gradientForest v0-1.37 (Ellis et al. 2012), with 500 trees and a correlation threshold of 0.5. Genomic offset was defined as the Euclidean distance between vectors of allelic turnover functions projected for each population under its native climate and under the climate conditions of the common garden. Climate variables were summarized using the first three main PC axes derived from 43 climate variables.

#### RDA (redundancy analysis)

In parallel, RDA was performed to train the model using vegan package v2.7-3, using the same three climate PC axes as described above. Genomic space under native and common garden climate conditions was predicted using the predict() function and the argument type=”lc”. Genomic offset was calculated as the Euclidean distance between vectors of the three RDA axes for each population under native and garden climate, weighted by relative importance of each climate axis.

### 4) Training sensitivity and validation stability

First, we tested model performance when trained on the training vs test datasets. Then, we explored the impact of downsampling of the training or test datasets on test model performance in three ways. We systematically removed: i) individual populations from the training dataset, leaving between 6 to 20 populations (with a step of 2) randomly selected in five draws, ii) individual trees from populations from the training dataset, leaving between a maximum of 2 to 10 trees per population (with a step of 2) randomly selected in ten draws, and iii) individual trees from populations from the test dataset, leaving between a maximum of 2 to 10 trees per population (with a step of 2) randomly selected in ten draws. All models were trained using 1000 random markers.

### 5) Stratified validation

We validated model performance by calculating offset-trait correlations for all tested populations, or only for populations from individual genetic clusters (West, Central or East). Each population was classified into one of the three clusters based on individual-ancestry frequency information from admixture analysis. First, trees were assigned to a genetic cluster, and populations were then assigned using the majority rule.

### 6) Model transferability

We investigated whether: i) regional models (models trained independently on each genetic cluster) improve the predictions by capturing local adaptive gradients within a genetic cluster, and ii) models trained in one genetic cluster could predict fitness in another. Because each genetic cluster occupies a different climate niche, this approach effectively tests model transferability to novel environments. Genetic clusters were defined as described above. Models were trained as described above, using training datasets of 1000 random markers and three main climate PC axes as predictors.

### 7) Climate predictor sensitivity

To test the impact of the choice of climate variables on model performance, we trained models using two sets of climate variables: i) the three main principal component (PC) axes derived from 43 climate variables, and ii) three uncorrelated climate variables (dPP, CMI, and fallMT) selected based on their Gradient Forest importance rankings. These three variables were selected out of a candidate pool consisting of the 15 variables from the small set of climate variables to ensure alignment with available future climate predictors.

#### Model validation calculation and consistency

To validate genomic offsets, Spearman’s rank correlation was calculated if at least five populations had estimates of both genomic offset and mean trait value. We also ensured that at least five trees per population were present in the training dataset and three trees per population were present in the test dataset. 95% confidence intervals were computed with 1000 bootstraps.

Spearman rho coefficients between fitness and climate transfer distance were very similar whether or not spatial autocorrelation was accounted for. For consistency, we therefore report all estimates in the validation section without spatial autocorrelation correction. To investigate consistency in genomic offset projections between models, we calculated Pearson’s correlation coefficients.

#### Genomic vulnerability of black spruce to future climate

Future simulated annual means of a small set of climate variables (CMI, mayMinT, TP, MAT, MMinT, MMaxT, sumTP, WSTP, winMT, sprMT, sumMT, fallMT, MCMT, MWMT, dPP) were averaged across four 20-year periods: 2021-2040, 2041-2060, 2061-2080, and 2081-2100. To illustrate the direction and magnitude of projected climate change across emission scenarios, regions and time periods we classified past and multiple future population climates into a limited number of climate types. Climate variables were scaled and clustered using K-means (KMeans() scikit-learn v1.7.0) testing 1 to 25 clusters. For each value of K, initial cluster centres were selected using a single run of the greedy k-means++ algorithm (n_init =’auto’), which spreads initial centres across the data. The optimal number of clusters was selected using the elbow method (yellowbrick v1.5), which identifies the point beyond which adding clusters no longer substantially decreases the distortion score (the sum of squared distances of data points to their cluster centre). The method split all data points into ten clusters, which were visualised on the two main principal components. The resulting climate types were used solely as a visualization tool to summarize similarities among historical and projected climates.

Gradient Forest was run as described above on a subset of 1000 randomly selected SNPs and allele frequencies calculated for each population including those from genetic clusters ME and WI to span the full range of black spruce genetic diversity. SNPs with any missing frequency estimates or no variation were excluded. Genomic offsets were calculated for the four 20-year periods and SSP2-4.5 emission scenario. Two models were trained: i) model trained on all populations and the three main PCs derived from a small set of climate variables to project genomic offsets in the Central/East cluster, and ii) model trained on populations from the West cluster and a subset of three selected variables (dPP, CMI, and fallMT), to project genomic offsets in the West cluster. Although we used two different climate datasets (BioSIM and Ouranos) in order to maintain consistent climate variables across past and future periods, future projections would ideally be based on a single climate dataset. Otherwise, systematic differences between datasets may be confounded with the projected climate change, potentially biasing genomic offset estimates, particularly in regions or periods where projected climate changes are relatively small.

Primary filtering analyses were performed using R v4.4.2 (R Core Team 2024), while all other analyses were conducted using R v4.6.2 (R Core Team 2026) and Python v3.12.

## Results

### Black spruce population structure

We analyzed 1467 black spruce individuals from 66 populations (Fig. 1A, Suppl. Table 1, Suppl. Dataset 1). We identified 3 main genetic clusters of black spruce, West, Central and East, with most samples showing mixed ancestry between West and Central or Central and East clusters (Fig. 1B-C). PCA showed similar results, with the 1st PC separating samples from cluster West, and the 2nd PC spreading samples from clusters East and Central, but with no clear separation (Fig. 1D). At higher values of K, the admixture analysis distinguished two additional subclusters: WI (samples from a population in Wisconsin) within the Central cluster, and ME (samples from a population in Maine) within the East cluster (Fig. 1B-C). *F*_ST_ was 0.012-0.014 between the cluster West and the clusters East or Central, about ten times higher than between clusters Central and East (*F*_ST_=0.001, Fig. 1E). Clusters WI and ME were most differentiated from cluster West (*F*_ST_=0.023 and 0.025, respectively) and from each other (*F*_ST_=0.024). AMOVA showed low but significant variance between clusters (% variance between clusters=4.6%, *P*<0.01, Suppl. Table 6).

Black spruce populations showed a significant pattern of isolation by distance (Mantel’s R=0.78, *P*<1×10^-4^, Suppl. Fig. 1). The inferred genetic clusters were geographically separated in space (Fig. 1B), and the two main axes of spatial autocorrelation (Moran eigenvector maps, MEMs) overlapped with the distribution of these clusters (Suppl. Fig. 2), indicating that geographic distance is an important contributor to genetic differentiation among black spruce populations. The two main axes of spatial PCA, confirmed separation of cluster West, and a split between clusters East and Central (Suppl. Fig. 3), lending support to the presence of three genetic clusters. The gradual transition in ancestry coefficients between neighboring genetic clusters is therefore consistent with the strong spatial genetic structure observed across the species range.

Observed heterozygosity ranged between 0.10 and 0.12 per population (Fig. 1F). Most populations from the West cluster showed lower heterozygosity and fewer polymorphic sites. When heterozygosity was measured excluding monomorphic sites within each population, West cluster populations showed higher observed heterozygosity despite lower number of polymorphic sites (Suppl. Fig. 4), showing that in the West cluster heterozygosity is driven by many monomorphic sites rather than the higher fraction of homozygous sites.

### Local adaptation of black spruce to climate

The three genetic clusters of black spruce occupied different climate niches. Population locations were characterized in terms of climate by compiling 43 climate variables averaged across 30 years (from 1961 to 1990, Suppl. Table 3, Suppl. Dataset 3). The first two principal components (PCs) summarising those variables distinguished populations from the West, Central, and East, with eastern locations being warmest and most humid and western ones being driest and coldest (Fig. 2A). Partial RDA analysis conditioned on genetic structure, spatial structure and climate variables showed that each class of variables explained only a small portion of genetic variation on its own (1.8-3.6% of explained variance, *P*=0.001, Suppl. Table 7). Climate alone explained 1.1% of genetic variation (adj. *R*^2^=0.011), corresponding to 2.6% of the explained variance, while most of the explained variation was shared with spatial and genetic structure. This pattern indicates that climate, geography, and demographic history are strongly confounded across the species range, making it difficult to disentangle their individual contributions to genomic variation. To identify which environmental variables were best associated with distribution of genetic diversity, we ran Gradient Forest on population frequencies estimated for 1000 randomly selected variants. Photoperiod, climate moisture index, precipitation, and fall frost probability were among the most important, besides the main axes of spatial autocorrelation (Fig. 2A-B).

Measurements of multiple phenotypic traits in common gardens revealed that black spruce populations originating from climates distant from the garden’s conditions tended to show reduced fitness compared to those from similar climates, implying local adaptation to climate. Height, total accumulated biomass (ABtot), and diameter at breast height (DBH) showed the strongest negative correlations with climate distance, consistent with local adaptation, although these correlations were consistently weaker in PR (rho>-0.39, Fig. 2C-D). Survival rate and wood density were generally weakly correlated. Five-year accumulated biomass (AB5) traits showed variable patterns depending on tree age and common garden: with stronger negative correlations observed at later growth stages in CH and PR, and at younger stages in ML and AC (Fig. 2D). Also, metrics of drought resilience varied, with weakly negative correlations observed only for resilience (Rs) and resistance (Rt) in AC and ML (Fig. 2D).

### Testing genomic offset with common gardens

#### Common gardens and phenotypic traits (Factor 1)

Using the genomic offset validation framework described in Fig. 3, model performance varied across common gardens and traits (Factor 1; Fig. 4A, Suppl. Fig. 5). In ML, CH and AC, traits that were more correlated with climate transfer distance also tended to show stronger correlations with genomic offset, although genomic offsets rarely outperformed climate-based correlations. The highest model performance was observed in AC, despite the training dataset including the fewest populations (n=10), and most test populations being absent from the training set. In contrast, in PR, genomic offsets were either uncorrelated or positively correlated with fitness traits.

**Figure 4.**
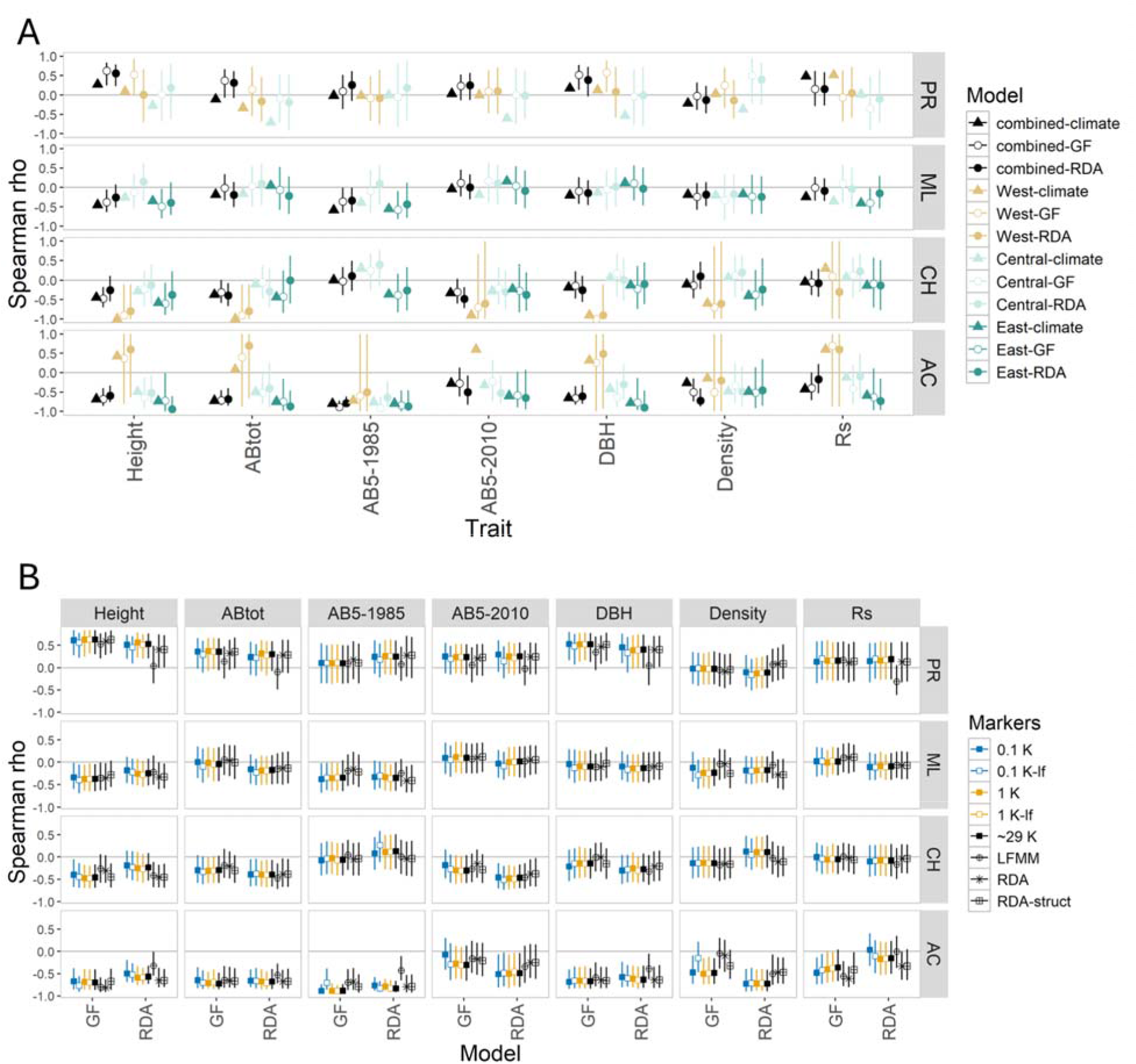
Accuracy of genomic offsets projections across models and marker sets. A. Comparison of rank correlations between fitness traits and climate transfer distance or genomic offsets from two models (Gradient Forest - GF and redundancy analysis - RDA), when evaluated for all populations (combined) or individual genetic clusters. B. Comparison of rank correlations between models trained on eight different marker sets, including all populations (combined). The marker types included subsets of 100 (0.1K), 1000 (1K) or all available markers (∼29K), subsets including low-frequency variants (0.1K-lf, 1K-lf), and markers found to be in association with climate (LFMM, RDA, RDA-struct).

Across AC, CH and ML, height and ABtot showed the strongest negative correlations with both climate distance and genomic offsets. Early interval AB5 were more negatively correlated with genomic offsets in AC and ML, whereas later AB5 showed stronger correlation in CH. Overall, genomic offsets were weakly associated with drought resistance traits, although negative correlations were observed for Rs and Rt in AC.

#### Marker types (Factor 2)

No consistent differences in model performance were observed when using random markers, all available markers, or subsets of markers associated with three climate PCs, either corrected or uncorrected for genetic structure (Factor 2; Fig. 4B, Suppl. Fig. 6). LFMM outliers represented an exception to the general pattern: they improved model performance in PR across most traits, but reduced performance in AC for RDA models and in CH for Gradient Forest models.

Genomic offsets derived from genome-wide marker sets were highly correlated with each other (Suppl. Figs. 7-10), but less so with two climate-associated markers (LFMM and RDA) in both Gradient Forest and RDA models. LFMM outliers also showed the lowest heterozygosity among marker types (Suppl. Fig. 11), suggesting an enrichment in markers with locally fixed alleles.

#### Genomic offset models (Factor 3)

The two modeling approaches, Gradient Forest and RDA, produced broadly similar results, although mean performance varied slightly depending on the garden and trait (Factor 3; Fig. 4A, Suppl. Fig. 5). Gradient Forest performed better on average for traits such as height and Rt in AC, whereas RDA showed stronger correlations with wood density in AC or AB5-1990 in ML.

#### Training sensitivity and validation stability (Factor 4)

Sensitivity to the training dataset was moderate (Factor 4; Suppl. Figs. 12-14). Model performance was usually slightly higher when models were trained on the test datasets compared to the training datasets, suggesting reduced predictive accuracy for unseen genotypes. However, these differences were very small. Among the most significant results in AC, model performance was higher when models were trained on the training dataset than on the test dataset for height and AB5-1985 (Suppl. Fig. 12), most likely because of the smaller number of populations in the test dataset.

The effect of increasing the number of populations in the training dataset varied depending on the garden, model type and trait (Suppl. Fig. 13). For example, for height, adding populations improved performance in Gradient Forest models (AC, ML, CH), but reduced performance in RDA models (AC, CH). In PR, using fewer populations in the training dataset brought offset-trait correlations closer to zero.

The effect of increasing the number of individuals per training population also varied, depending on the garden, model type, and trait (Suppl. Fig. 14). For height and ABtot, increasing sample size had little or no effect, or reduced model performance, in particular for RDA.

The influence of population size in test datasets was more evident (Suppl. Fig. 15). Model performance in AC decreased with smaller test population sizes. In ML and CH, performance was more stable and declined primarily for the smallest populations. In PR, where offset-trait correlations were positive, correlations generally tended towards zero as test population size decreased.

#### Stratified validation (Factor 5)

When evaluated at the level of genetic clusters, model performance was not consistent across clusters (Factor 5; Fig. 4A, Suppl. Fig. 5), a pattern also observed for climate-trait relationships. Offset-trait correlations for the East cluster were negative across all common gardens. In contrast, the Central cluster showed negative correlations only in AC, and the West cluster only in CH. Although discrepancies between climate distance and genomic offset persisted in PR, they were less pronounced for RDA models.

#### Model transferability (Factor 6)

For models trained on populations belonging to individual genetic clusters, model performance was similar to that of models trained on the combined datasets (Factor 6; Fig. 5, Suppl. Fig. 16-17). However, genomic offsets derived from models trained on different clusters were not always strongly correlated. In PR, offsets obtained from models trained on the Central cluster were often poorly correlated with those trained on the West cluster or on the combined dataset (Suppl. Figs. 18-19). In CH, strong agreement was observed only between RDA models trained on combined and East, or Central and West datasets. In addition, models performing well in one cluster, did not have equal performance in other clusters or across the species range (Fig. 5, Suppl. Fig. 16-17). Climate explained a significant part of genetic variation at the cluster level, even though most of it was confounded with genetic and spatial structure (Suppl. Table 8), indicating that the lack of improvement from cluster-specific models was not due to an absence of genotype–climate association within clusters.

**Figure 5.**
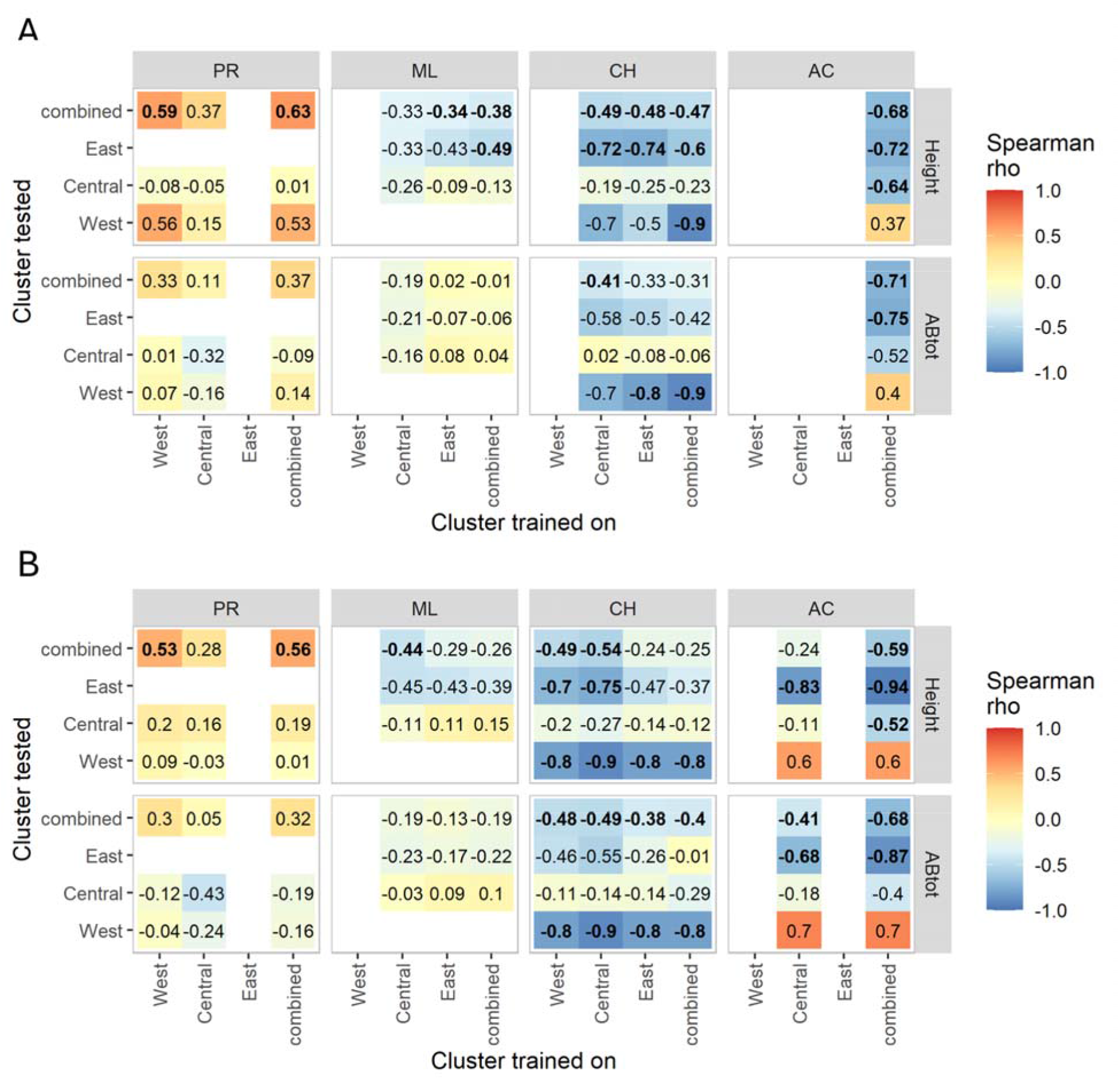
Comparison of models trained and evaluated on combined datasets or individual genetic clusters using three major climate PCs. A. Genomic offset-trait rank correlations for Gradient Forest models. B. Genomic offset-trait rank correlations for RDA models. All models were trained using 1000 random markers.

#### Climate variable selection (Factor 7)

Model sensitivity to climate predictors varied across gardens (Factor 7; Fig. 6, Suppl. Figs. 20-23). In ML, CH, and AC, model performances generally decreased when using three climate variables instead of three climate PCs, regardless of whether the models were trained on combined datasets or individual clusters only, except for those trained on the West cluster in CH. In contrast, in PR, offset-trait correlations obtained with Gradient Forest in West clusters improved and were nearly always negative.

**Figure 6.**
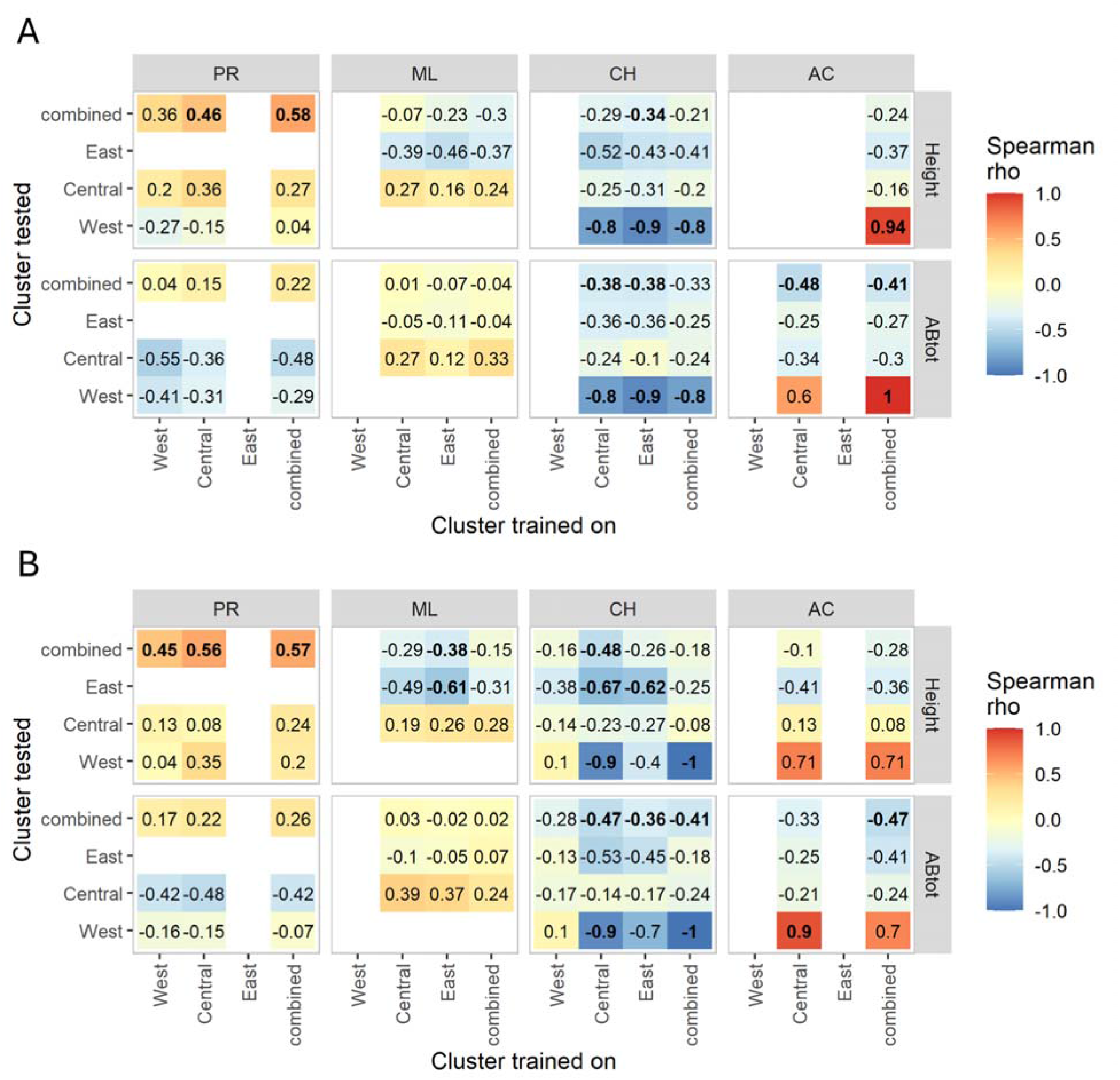
Comparison of models trained and evaluated on combined datasets or individual genetic clusters using three climate variables (dPP, CMI and fallMT). A. Genomic offset-trait rank correlations for Gradient Forest models. B. Genomic offset-trait rank correlations for RDA models. All models were trained using 1000 random markers.

Genomic offsets of models trained on three PCs vs three climate variables were weakly correlated, with mean of 0.55 for Gradient Forest models (range from-0.07 and 0.97) and 0.72 for RDA (range from-0.65 to 0.97; Suppl. Figs. 24-25).

### Vulnerability of black spruce populations to future climate

Common garden experiments are well suited for testing hypotheses regarding the consequences of assisted migration; however, our broader objective is to predict how natural populations will respond to future climate conditions. To visualize the projected climate shifts across the range of black spruce, we classified the climate state of each population in the past and across four future periods (2021-40, 2041-60, 2061-80, and 2081-2100) under three carbon emission scenarios into ten clusters (Suppl. Fig. 26). The West, Central, and East genetic clusters were projected to experience progressively warmer climates over time and along the south-north gradient (Suppl. Fig. 26).

Given the differences in model performance observed during validation, particularly between western and central/eastern common gardens, when depending on climate predictors, we adopted a region-specific modeling approach. In the western part of the species range, genomic offsets were projected to increase across most of the region, reaching their highest values in the northwest (Fig. 7A). In contrast, in the central and eastern regions, projected genomic offsets were generally higher in central populations than in eastern ones (Fig. 7B). Given the variable model performance across gardens and genetic clusters, these projections should be regarded as exploratory.

**Figure 7.**
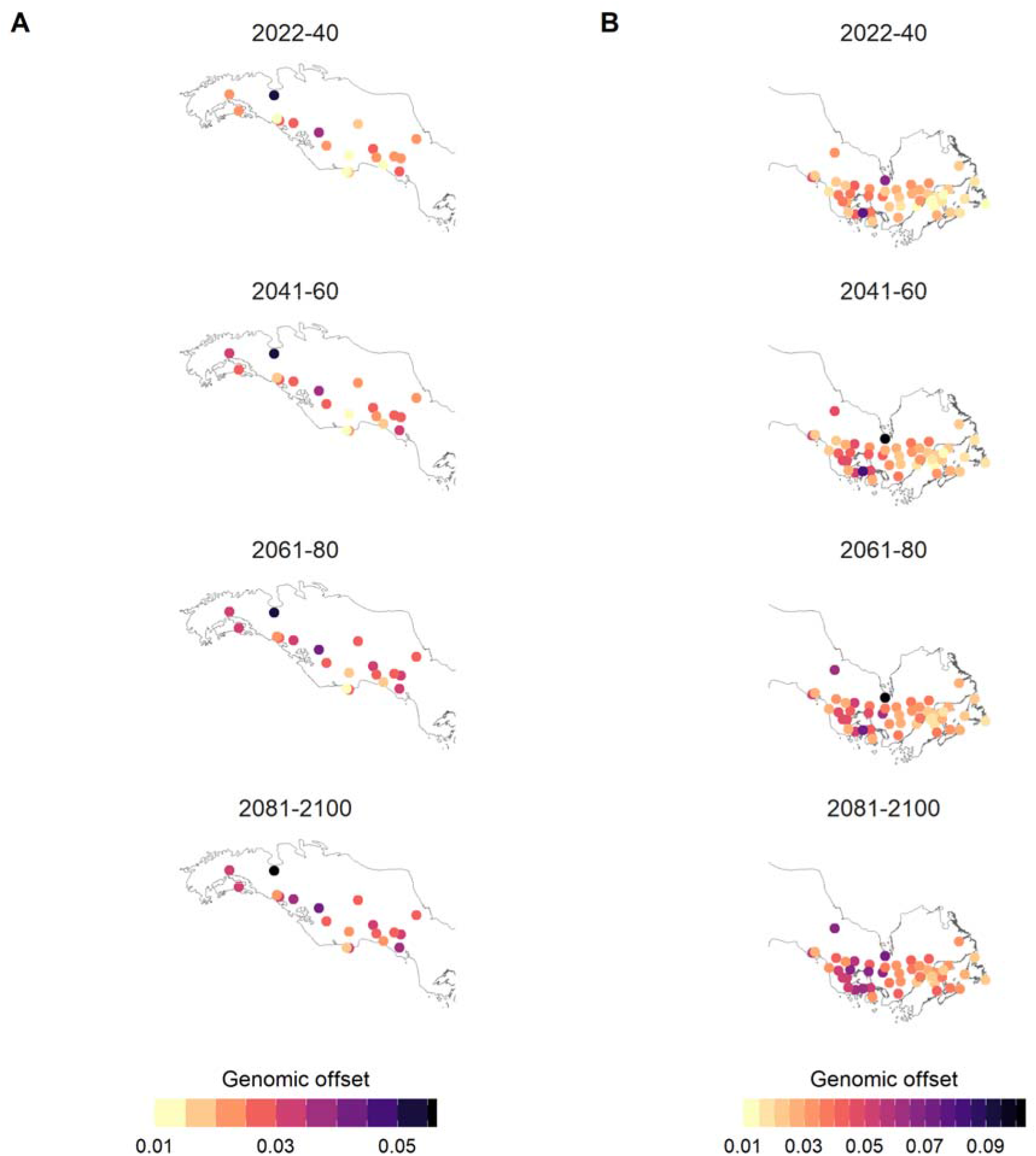
Projected genomic offsets under future climatic scenarios. Genomic offsets were projected for four 20-year periods (2021–2040, 2041–2060, 2061–2080 and 2081–2100) under the medium emission scenario (SSP2-4.5). They were projected separately for each region using the Gradient Forest model and climate predictors that performed best in validation for that region. A. Genomic offset projected across four time periods using a model trained on the West cluster dataset and three climate variables (dPP, CMI and fallMT). B. Genomic offsets projected across four time periods using a model trained on the combined dataset and three climate PCs. Genomic offsets indicate relative differences among populations within each region and should not be compared between panels A and B.

## Discussion

### Distribution of genetic diversity in black spruce

Black spruce carries high levels of genetic variation and has large estimated population sizes across its distribution due to its outcrossing mating system and wind pollination (Isabel et al. 1995; Perry and Bousquet 2001; Bouillé and Bousquet 2005). Our results confirm that variation is sorted among three widespread genetic clusters: West, Central, and East, with the West cluster being most distinct (Gérardi et al. 2010; Prunier et al. 2012; Jaramillo-Correa et al. 2004). This structure is interpreted as the legacy of the last glacial maximum (Jaramillo-Correa et al. 2004), although our data does not exclude the possibility that Central and East clusters were derived from a single glacial refugium, and later diverged after migrating and adapting to distinct environmental conditions. We also identified two isolated subclusters, one within Central (WI) and one within East cluster (ME). While WIs range is most likely limited to a small area, in case of ME it’s difficult to judge how widespread this subcluster is as it is situated at the southern limits of the sampling range, but not at the limit of the species range. Pairwise *F*_ST_s show these two subclusters are genetically differentiated from other clusters, from each other, as well as from the red spruce, which suggests that they could be remnants of glacial microrefugia that failed to expand across the landscape. Such cryptic refugia were indeed identified in other North American species (Shafer et al. 2010; Fernandez et al. 2021).

Heterozygosity estimates based on variant sites only are driven in structured populations by the amount of divergence between those populations which determines the amount of population-specific non-variant sites left in the dataset (Schmidt et al. 2021). This proves to be the case in black spruce, as observed heterozygosity is the lowest in West populations when all sites in a species are considered, but it’s highest in West populations when only polymorphic sites per population are considered. This would suggest that West populations do not necessarily have smaller effective population sizes than East or Central populations.

### Limitations to genomic offset in black spruce

Overall, our validation framework revealed that within common gardens, genomic offset projections were largely invariant to model choice, marker number, or size of the dataset used for model training. In line with accumulating literature, genomic offsets performance was context dependent (Fitzpatrick et al. 2025), varying with the common garden considered, the fitness traits selected for validation, and climate predictors used in the models. In addition, genetic structure influenced model outcomes, as genomic offsets estimated at the species level were not consistent with those calculated within individual genetic clusters.

### Genomic offset and climate-distance projections are garden-dependent

A key assumption underlying genomic offset metrics is that local populations are under stabilizing selection for optimal phenotypes (Gain et al. 2023). Using common garden experiments, we demonstrated that population response to climatic conditions varied depending on the garden environment, the traits measured, and the genetic background (genetic cluster to which population belonged). Model performances varied mostly between gardens, with the strongest results observed in AC, and the weakest in PR, where genomic offsets were either not correlated or positively correlated with fitness traits. Two reasons may account for discrepancy in PR. First, the presence of non-crossover genotype-by-environment interactions, where populations from the Central cluster grow on average better than those from the West cluster (Robert et al. 2024); this may result in lower relative fitness for West cluster trees despite potential local adaptation. Second, populations in the West cluster may respond to a different set of climatic drivers than those included in the species-wide models. Our results support both scenarios, i.e. decoupling of genomic offsets and fitness traits at the species level (Suppl. Fig. 27), and model performance improvement for different sets of climate variables (Figs. 5-6).

Apart from lack of model consistency across gardens, we also observed that model performance varied depending on the genetic cluster on which genomic offset was tested. This suggests that species-wide models may not adequately capture processes operating at local scales. Such regional-dependency resonates with recent findings conducted in genetically structured maritime pine (*Pinus pinaster* Ait.), which found mixed results of offset-fitness correlations across multiple common gardens (Archambeau et al. 2025).

In contrast to previous validation studies (Fitzpatrick et al. 2021; Lachmuth et al. 2024), genomic offset models rarely outperform climate transfer distance, both at species and cluster-level. Genomic offset is expected to model fitness loss better than climate, because climate predictors are weighted by their effects on association with genetic variation (Gain et al. 2023). However, simulations of markers adapted to known causal environmental gradients in common gardens showed that climate-based distance can be an equally good predictor of fitness as Gradient Forest-based genomic offsets, in particular when causal environments are orthogonal and non-linear (Láruson et al. 2022). Moreover, in Douglas-fir, models trained on climate variables identified as influential for fitness in common garden experiments proved more accurate than those trained on other variables (Lind et al. 2024). Together, these findings suggest that climate PCs used in our models capture key environmental gradients driving fitness variation in most common gardens.

### Projections are trait-dependent

The selection of fitness traits clearly influenced model performance, as not all traits are good predictors of environmental change. Height, total accumulated biomass, and DBH in general showed the strongest negative correlations with climate distance and genomic offsets. These traits are commonly used as proxies for fitness in tree species, because they respond to climatic variation in common garden experiments and show moderate heritability (Fitzpatrick et al. 2021; Housset et al. 2018; Girardin et al. 2021; Robert et al. 2024). Interestingly, short-term accumulated biomass showed contrasting patterns across the gardens depending on tree age. In the two warmest gardens (ML and AC), early growth intervals of biomass were better associated with climate distance and genomic offsets, whereas in the two coldest gardens (PR and CH), later growth intervals showed the strongest associations, both at the species and cluster levels. Low correlations between juvenile and mature traits have been previously observed in conifers (Depardieu et al. 2021). In terms of genomic offset, these results only partially align with recent findings in red spruce, where stronger associations were observed for juvenile-stage traits, when compared to later developmental stages (Verrico et al. 2025). As red spruce gardens were located at lower latitudes than those of black spruce, this discrepancy may reflect differences in temperature or photoperiod effects on early vs late-stage response of trees. Although survival can be a useful metric for fitness (Archambeau et al. 2025; Lind et al. 2024), we found it responded weakly to climate, mostly in PR and AC. This may be explained by the absence of early-stage mortality in our measurements, possibly leading to this weak signal.

Mean drought-related metrics also varied across gardens and genetic clusters but were only weakly associated with climatic distance or genomic offsets. An independent dendroecological study conducted in the same trials found that trade-offs between drought resistance (Rt) and post-drought recovery (Rc) varied only weakly among provenances, while their expression depended largely on drought intensity at each garden (Robert et al., in prep.). This pattern is consistent with the weak associations observed between drought-related metrics and genomic offset.

### Factors influencing model performance

We did not find consistent effects of model choice or marker number on model performance. This is generally consistent with previous studies validating genomic offsets in common gardens or via simulations (Lind et al. 2024; Láruson et al. 2022; Fitzpatrick et al. 2021; Lind and Lotterhos 2025a; Verrico et al. 2025), and likely reflect that a large part of segregating variation in black spruce is linked to the environment as a consequence of spatial distribution of genetic clusters. Despite broadly similar model performance across marker sets, genome-wide and neutral markers did not always give the same genomic offset rankings, in particular for LFMM markers and in RDA models. LFMM outliers displayed lower heterozygosity compared to other markers, suggesting a higher fraction of locally fixed alleles, and rendering it unequally informative across the species distribution.

Increases in model performance with larger training or test population sizes are expected when populations are genetically and phenotypically diverse, due to reduced sampling error caused by measurement or other non-genetic effects. In black spruce, a large proportion of genetic and phenotypic variance is distributed within populations (Rossi and Bousquet 2014). Consistent with this expectation, smaller test population sizes indeed led to weaker offset-trait correlations in the best performing models. This pattern aligns with simulation studies showing that genomic offsets based on population-level fitness means outperform those based on individual-level measurements (Lind and Lotterhos 2025a).

On the other hand, reducing the size or number of populations in the training dataset did not have a consistent impact on model training. One possible explanation is that, given the high genetic diversity of black spruce, even a limited number of randomly sampled populations may capture enough genetic variation and its association with climate to train effective models. In some cases, we even observed model improvement when a smaller size or a lower number of populations were used for training. This behavior could potentially arise if traits were correlated with genomic offset at the species level but not at the genetic cluster level–specifically, if correlation strength varied among clusters, meaning that the subsampling or removal of specific trees could inadvertently enrich the dataset for groups with stronger correlations.

Model performance in some gardens was strongly influenced by the selection of climate variables used for training. In particular, the two sets of climate predictors yielded contrasting results in AC and PR, indicating that predictor selection can have a decisive impact on model performance. This likely reflects the regional differences in the climatic factors limiting black spruce growth (D’Orangeville et al. 2018). In eastern regions, where precipitation is high, low temperatures limit growth in the north, whereas in the south, higher temperatures can constrain growth through higher evapotranspiration (Girardin et al. 2016; D’Orangeville et al. 2018; Lesven et al. 2024). In the western and central regions, lower precipitation makes moisture availability a major determinant of growth (Lesven et al. 2024; Price et al. 2013; Girardin et al. 2016, 2024). Because genomic offset measures the magnitude of climatic change regardless of its direction and assumes that populations are adapted to their local climate, changes expected to increase growth (e.g. warming in cold-limited regions) and those expected to decrease it (e.g. warming in moisture-limited regions) are both interpreted as maladaptation. Consequently, no single model was able to accurately predict population responses to climate across the four common gardens. This finding further highlights the importance of identifying the key environmental drivers underlying genomic offset estimation and selecting predictors that reflect the ecological processes shaping fitness variation.

Previous studies pointed to the importance of the causal environmental predictors in certain conditions in genomic offset performance (Láruson et al. 2022; Lind et al. 2024). However, these drivers are rarely known a priori, and Gradient Forest does not always reliably identify them. Our results support this view, indicating that uncertainty in predictor choice is a key limitation of genomic offset approaches. A pragmatic strategy is to combine ecological knowledge (e.g. known growth-limiting factors), sensitivity analyses using alternative subsets of predictors, and dimension-reduction methods such as PCA, which summarize large sets of correlated climate variables into a few most evident ‘’synthetic’’ gradients within a region. While latter approaches can improve model robustness, they may also complicate biological interpretation.

Region-specific models are often expected to improve predictions by better capturing local drivers of adaptation. However, in our study they did not outperform species-wide models, despite the presence of a pronounced genetic structure across the range. This result is noteworthy because these genetic clusters correspond to distinct glacial lineages that likely persisted in separate refugia during the last glaciation and followed different evolutionary trajectories after postglacial recolonization (Prunier et al. 2012). Previous studies using the same populations have also shown that climate responses differ among regions, with western populations often exhibiting distinct growth responses and sensitivities to climatic stressors compared with central and eastern populations (Robert et al. 2024, 2025). Together these findings suggest that regional models may capture some cluster-specific adaptive signals, whereas species-wide models capture broader adaptive gradients shared across the range. Consistent with this interpretation, genomic offsets derived from models trained on different clusters were not always strongly correlated, suggesting that different training datasets can capture different components of adaptive variation. Nevertheless, these regional signals were not sufficient to consistently improve predictive performance relative to species-wide models. Moreover, because future climates within a region increasingly resemble present-day climates elsewhere in the species range, regional models are more likely to require extrapolation beyond their training range. The choice between regional and species-wide models should therefore be guided by empirical validation performance and accompanied by an explicit assessment of climatic extrapolation.

## Conclusions

By combining genome-wide sequencing and phenotype data from distribution-wide populations of black spruce grown in four long-term common garden experiments, we evaluated the ability of genomic offset models to predict fitness in this genetically structured species. Although genomic offset was informative in some contexts, its predictive power was limited across the full species range. This limitation appears to arise, first, because similar ranks in genetic cluster relative fitness are retained between gardens, which prevents reliable projection of fitness across the species, and second, because species responses to climate vary across the landscape not only as a function of individual climatic variables but also by their interactions. Despite these limitations, models trained on carefully selected climate predictors proved to be robust to the choice of markers, but also to the size, amount and range of populations used in model training. Future directions should prioritize the establishment of new common gardens, especially in western regions, where population responses to climate are less well understood. Improving genotype quality and increasing sampling density could also prove useful for refining genomic offset projections by enabling more accurate modeling of fine-scale genotype-environment associations. Finally, incorporating soil properties and associated microbiota in the future, may help improve trait response curves under changing climate conditions.

## Data Archiving Statement

Raw and filtered variant datasets are publicly available on Zenodo at https://doi.org/10.5281/zenodo.19961099. Supplementary Datasets 1–4 are archived at https://doi.org/10.5281/zenodo.20563726, and the associated analysis code is available at https://doi.org/10.5281/zenodo.20563774.

## Supplementary materials

Supplementary Figure 1. Patterns of isolation by distance across species distribution.

Supplementary Figure 2. Four major axes of spatial autocorrelation.

Supplementary Figure 3. Spatial visualization of sPCA four main axes.

Supplementary Figure 4. Observed heterozygosity versus number of polymorphic sites across populations of black spruce.

Supplementary Figure 5. Performance of Gradient Forest (GF) and redundancy analysis (RDA) models validated against all fitness traits and estimated for all populations (“combined”) or individual genetic clusters.

Supplementary Figure 6. Comparison of predictive performance of Gradient Forest (GF) and redundancy analysis (RDA) models trained on eight different marker types and estimated for all populations.

Supplementary Figure 7. Matrices of pairwise correlations between genomic offsets, estimated via Gradient Forest models trained on different sets of markers validated for the first eight fitness traits.

Supplementary Figure 8. Matrices of pairwise correlations between genomic offsets, estimated via Gradient Forest models trained on different sets of markers validated for the last eight fitness traits.

Supplementary Figure 9. Matrices of pairwise correlations between genomic offsets, estimated via RDA models trained on different sets of markers validated for the first eight fitness traits.

Supplementary Figure 10. Matrices of pairwise correlations between genomic offsets, estimated via RDA models trained on different sets of markers validated for the last eight fitness traits.

Supplementary Figure 11. Observed heterozygosity calculated per population using selected marker sets: all variants (All), LFMM outliers (LFMM), RDA outliers (RDA) and RDA outliers corrected for genetic structure (RDA-struct).

Supplementary Figure 12. Comparison of predictive performance between models trained on training (axis x) and test datasets (axis y).

Supplementary Figure 13. Testing the impact of the number of populations in the training dataset on model performance.

Supplementary Figure 14. Testing the impact of the population size in the training dataset on model performance.

Supplementary Figure 15. Testing the impact of the population size in the test dataset on model performance.

Supplementary Figure 16. Gradient Forest model performances when trained on individuals genetic clusters or combined datasets and three climate PCs in predicting fitness traits of the same or other genetic clusters.

Supplementary Figure 17. RDA model performances when trained on individuals genetic clusters or combined datasets and three climate PCs in predicting fitness traits of the same or other genetic clusters.

Supplementary Figure 18. Matrices of pairwise correlations between genomic offsets estimated using Gradient Forest models trained on single genetic clusters or combined datasets and three climate PCs.

Supplementary Figure 19. Matrices of pairwise correlations between genomic offsets estimated using RDA models trained on single genetic clusters or combined datasets and three climate PCs.

Supplementary Figure 20. Gradient Forest model performances when trained on individuals genetic clusters or combined datasets and three selected climate variables in predicting fitness traits of the same or other genetic clusters.

Supplementary Figure 21. RDA model performances when trained on individuals genetic clusters or combined datasets and three selected climate variables in predicting fitness traits of the same or other genetic clusters.

Supplementary Figure 22. Matrices of pairwise correlations between genomic offsets estimated using Gradient Forest models trained on single genetic clusters or combined datasets and three selected climate variables.

Supplementary Figure 23. Matrices of pairwise correlations between genomic offsets estimated using RDA models trained on single genetic clusters or combined datasets and three selected climate variables.

Supplementary Figure 24. Matrices of pairwise correlations between genomic offsets estimated using Gradient Forest models trained on two different climate datasets: a selected subset of three climate variables (“var3”, axis x) and three climate PCs (“PC3”, axis y).

Supplementary Figure 25. Matrices of pairwise correlations between genomic offsets estimated using RDA models trained on two different climate datasets: a selected subset of three climate variables (“var3”, axis x) and three climate PCs (“PC3”, axis y).

Supplementary Figure 26. Change of climate across Canada in the current century. A. Ordination of all population climate states (current and future projections according to three emission scenarios) along the first two principal components derived from 15 climate variables (arrows).

Supplementary Figure 27. Example of a confounding effect of differences in mean biomass increment between clusters on genomic offset estimates.

Supplementary Table 1. Geographical location of all initial black spruce populations (provenances) and the number of retained tree genotypes in each common garden after filtering.

Supplementary Table 2. Geographical location and mean climate conditions of common gardens.

Supplementary Table 3. List of climate variables.

Supplementary Table 4. Number of populations assigned to test and training datasets.

Supplementary Table 5. Number of genotypes assigned to test and training datasets.

Supplementary Table 6. Results of AMOVA.

Supplementary Table 7. Results of partial RDA analysis

Supplementary Table 8. Partition of variance explained by climate in partial RDA models fitted to species-wide or within-cluster data

Supplementary Dataset 1. Tree sample information.

Supplementary Dataset 2. Sampling coordinates of red spruce populations. Supplementary Dataset 3. Past climate normals for 43 climate variables.

Supplementary Dataset 4. Population means and standard errors of measured phenotypic traits.

## Literature cited

Aguirre-Liguori, Jonás A., Santiago Ramírez-Barahona, and Brandon S. Gaut. 2021. “The Evolutionary Genomics of Species’ Responses to Climate Change.” Nature Ecology & Evolution 5 (10): 1350–1360.

Alexander, David H., John Novembre, and Kenneth Lange. 2009. “Fast Model-Based Estimation of Ancestry in Unrelated Individuals.” Genome Research 19 (9): 1655–1664.

Archambeau, Juliette, Marta Benito Garzón, Marina de-Miguel, et al. 2025. “Evaluating Genomic Offset Predictions in a Forest Tree with High Population Genetic Structure.” The American Naturalist, no. 739045 (October). 10.1086/739045.

Aubin, I., L. Boisvert-Marsh, H. Kebli, et al. 2018. “Tree Vulnerability to Climate Change: Improving Exposure-Based Assessments Using Traits as Indicators of Sensitivity.” *Ecosphere (Washington*, D.C) 9 (2): e02108.

Baltzer, Jennifer L., Nicola J. Day, Xanthe J. Walker, et al. 2021. “Increasing Fire and the Decline of Fire Adapted Black Spruce in the Boreal Forest.” Proceedings of the National Academy of Sciences of the United States of America 118 (45): e2024872118.

Bellemin-Noël, Bastien, Stéphane Bourassa, Emma Despland, Louis De Grandpré, and Deepa S. Pureswaran. 2021. “Improved Performance of the Eastern Spruce Budworm on Black Spruce as Warming Temperatures Disrupt Phenological Defences.” Global Change Biology 27 (14): 3358–3366.

Bergmann, F. 1978. “The Allelic Distribution at an Acid Phosphatase Locus in Norway Spruce (Picea Abies) along Similar Climatic Gradients.” Theoretical and Applied Genetics 52 (2): 57–64.

Bivand, Roger S., and David W. S. Wong. 2018. “Comparing Implementations of Global and Local Indicators of Spatial Association.” *Test (Madrid*, Spain) 27 (3): 716–748.

Bouillé, Marie, and Jean Bousquet. 2005. “Trans-Species Shared Polymorphisms at Orthologous Nuclear Gene Loci among Distant Species in the Conifer Picea (Pinaceae): Implications for the Long-Term Maintenance of Genetic Diversity in Trees.” American Journal of Botany 92 (1): 63–73.

Bradley St Clair, J., and Glenn T. Howe. 2007. “Genetic Maladaptation of Coastal Douglas fir Seedlings to Future Climates.” Global Change Biology 13 (7): 1441–1454.

Brandt, J. P., M. D. Flannigan, D. G. Maynard, I. D. Thompson, and W. J. A. Volney. 2013. “An Introduction to Canada’s Boreal Zone: Ecosystem Processes, Health, Sustainability, and Environmental Issues.” Environmental Reviews 21 (4): 207–226.

Burns, Russel M., and Barbara H. Honkala. 1990. Silvics of North America: Conifers. Forest Service, United States Department of Agriculture. Oxford.

Capblancq, Thibaut, and Brenna R. Forester. 2021. “Redundancy Analysis: A Swiss Army Knife for Landscape Genomics.” Methods in Ecology and Evolution 12 (12): 2298–2309.

Capblancq, Thibaut, Keurcien Luu, Michael G. B. Blum, and Eric Bazin. 2018. “Evaluation of Redundancy Analysis to Identify Signatures of Local Adaptation.” Molecular Ecology Resources 18 (6): 1223–1233.

Capblancq, Thibaut, Aurélien Tauzin, Yves Vigouroux, Philippe Cubry, and Olivier François. 2025. “Linking Genomic Offset Statistics to the Shape of Selection Gradients.” The American Naturalist, no. 739079 (October). 10.1086/739079.

Depardieu, Claire, Sébastien Gérardi, Simon Nadeau, et al. 2021. “Connecting Tree-Ring Phenotypes, Genetic Associations and Transcriptomics to Decipher the Genomic Architecture of Drought Adaptation in a Widespread Conifer.” Molecular Ecology 30 (16): 3898–3917.

Depardieu, Claire, Martin P. Girardin, Simon Nadeau, Patrick Lenz, Jean Bousquet, and Nathalie Isabel. 2020. “Adaptive Genetic Variation to Drought in a Widely Distributed Conifer Suggests a Potential for Increasing Forest Resilience in a Drying Climate.” The New Phytologist 227 (2): 427–439.

D’Orangeville, Loïc, Daniel Houle, Louis Duchesne, Richard P. Phillips, Yves Bergeron, and Daniel Kneeshaw. 2018. “Beneficial Effects of Climate Warming on Boreal Tree Growth May Be Transitory.” Nature Communications 9 (1): 3213.

Ellis, Nick, Stephen J. Smith, and C. Roland Pitcher. 2012. “Gradient Forests: Calculating Importance Gradients on Physical Predictors.” Ecology 93 (1): 156–168.

Evans, Luke M., Gancho T. Slavov, Eli Rodgers-Melnick, et al. 2014. “Population Genomics of Populus Trichocarpa Identifies Signatures of Selection and Adaptive Trait Associations.” Nature Genetics 46 (10): 1089–1096.

Fernandez, Matias C., Feng Sheng Hu, Daniel G. Gavin, Guillaume Lafontaine, Katy D. Heath, and Alain Vanderpoorten. 2021. “A Tale of Two Conifers: Migration across a Dispersal Barrier Outpaced Regional Expansion from Refugia.” Journal of Biogeography 48 (9): 2133–2143.

Fitzpatrick, Matthew C., Vikram E. Chhatre, Raju Y. Soolanayakanahally, and Stephen R. Keller. 2021. “Experimental Support for Genomic Prediction of Climate Maladaptation Using the Machine Learning Approach Gradient Forests.” Molecular Ecology Resources 21 (8): 2749–2765.

Fitzpatrick, Matthew C., and Stephen R. Keller. 2015. “Ecological Genomics Meets Community-Level Modelling of Biodiversity: Mapping the Genomic Landscape of Current and Future Environmental Adaptation.” Ecology Letters 18 (1): 1–16.

Fitzpatrick, Matthew C., Stephen R. Keller, and Katie E. Lotterhos. 2025. “The Challenge of Genomic Forecasting in an Era of Global Change.” The American Naturalist, no. 738891 (September). 10.1086/738891.

Fortin, Mathieu, Jean-François Lavoie, Jacques Régnière, and Rémi Saint-Amant. 2022. “A Web API for Weather Generation and Pest Development Simulation in North America.” Environmental Modelling & Software: With Environment Data News 157 (105476): 105476.

Gain, Clément, and Olivier François. 2021. “LEA 3: Factor Models in Population Genetics and Ecological Genomics with R.” Molecular Ecology Resources 21 (8): 2738–2748.

Gain, Clément, Bénédicte Rhoné, Philippe Cubry, et al. 2023. “A Quantitative Theory for Genomic Offset Statistics.” Molecular Biology and Evolution 40 (6): msad140.

Gauthier, S., P. Bernier, T. Kuuluvainen, A. Z. Shvidenko, and D. G. Schepaschenko. 2015. “Boreal Forest Health and Global Change.” *Science (New York*, N.Y.) 349 (6250): 819–822.

Gérardi, Sébastien, Juan P. Jaramillo-Correa, Jean Beaulieu, and Jean Bousquet. 2010. “From Glacial Refugia to Modern Populations: New Assemblages of Organelle Genomes Generated by Differential Cytoplasmic Gene Flow in Transcontinental Black Spruce: ASSEMBLAGES OF ORGANELLE GENOMES.” Molecular Ecology 19 (23): 5265–5280.

Girardin, Martin P., Xiao Jing Guo, William Marchand, and Claire Depardieu. 2024. “Unravelling the Biogeographic Determinants of Tree Growth Sensitivity to Freeze and Drought in Canada’s Forests.” The Journal of Ecology 112 (4): 848–869.

Girardin, Martin P., Edward H. Hogg, Pierre Y. Bernier, Werner A. Kurz, Xiao Jing Guo, and Guillaume Cyr. 2016. “Negative Impacts of High Temperatures on Growth of Black Spruce Forests Intensify with the Anticipated Climate Warming.” Global Change Biology 22 (2): 627–643.

Girardin, Martin P., Nathalie Isabel, Xiao Jing Guo, Manuel Lamothe, Isabelle Duchesne, and Patrick Lenz. 2021. “Annual Aboveground Carbon Uptake Enhancements from Assisted Gene Flow in Boreal Black Spruce Forests Are Not Long-Lasting.” Nature Communications 12 (1): 1169.

GlobalFRA. n.d. “Global Forest Resources Assessment 2020.” Accessed September 21, 2025. https://www.fao.org/forest-resources-assessment/past-assessments/fra-2020/en.

Gosselin, Thierry. 2020. Thierrygosselin/radiator: Update. Zenodo. 10.5281/ZENODO.3687060.

Gosselin, Thierry, Eric C. Anderson, Anne-Laure Ferchaud, and Ido Bar. 2019. Thierrygosselin/assigner: Assigner 0.5.6 2019-05-01. Zenodo. 10.5281/ZENODO.2656728.

Guénard, Guillaume, and Pierre Legendre. 2022. “Hierarchical Clustering with Contiguity Constraint in *R*.” Journal of Statistical Software 103 (7): 1–26.

Guo, Xiali, Marcin Klisz, Radosław Puchałka, et al. 2022. “Common garden Experiment Reveals Clinal Trends of Bud Phenology in Black Spruce Populations from a Latitudinal Gradient in the Boreal Forest.” The Journal of Ecology 110 (5): 1043–1053.

Hogg, E. H., A. G. Barr, and T. A. Black. 2013. “A Simple Soil Moisture Index for Representing Multi-Year Drought Impacts on Aspen Productivity in the Western Canadian Interior.” Agricultural and Forest Meteorology 178–179 (September): 173–182.

Housset, Johann M., Simon Nadeau, Nathalie Isabel, et al. 2018. “Tree Rings Provide a New Class of Phenotypes for Genetic Associations That Foster Insights into Adaptation of Conifers to Climate Change.” The New Phytologist 218 (2): 630–645.

IPCC. 2023. “Climate Change 2023: Synthesis Report. Contribution of Working Groups I, II and III to the Sixth Assessment Report of the Intergovernmental Panel on Climate Change.” Core Writing Team, H. Lee and J. Romero. Preprint, IPCC.

Isaac-Renton, Miriam G., David R. Roberts, Andreas Hamann, and Heinrich Spiecker. 2014. “Douglas-Fir Plantations in Europe: A Retrospective Test of Assisted Migration to Address Climate Change.” Global Change Biology 20 (8): 2607–2617.

Isabel, N., J. Beaulieu, and J. Bousquet. 1995. “Complete Congruence between Gene Diversity Estimates Derived from Genotypic Data at Enzyme and Random Amplified Polymorphic DNA Loci in Black Spruce.” Proceedings of the National Academy of Sciences of the United States of America 92 (14): 6369–6373.

Jaramillo-Correa, Juan P., Jean Beaulieu, and Jean Bousquet. 2004. “Variation in Mitochondrial DNA Reveals Multiple Distant Glacial Refugia in Black Spruce (*Picea Mariana*), a Transcontinental North American Conifer.” Molecular Ecology 13 (9): 2735–2747.

Jombart, Thibaut. 2008. “Adegenet: A R Package for the Multivariate Analysis of Genetic Markers.” Bioinformatics 24 (11): 1403–1405.

Kamvar, Zhian N., Javier F. Tabima, and Niklaus J. Grünwald. 2014. “Poppr: An R Package for Genetic Analysis of Populations with Clonal, Partially Clonal, And/or Sexual Reproduction.” PeerJ 2 (March): e281.

Kilian, Andrzej, Peter Wenzl, Eric Huttner, et al. 2012. “Diversity Arrays Technology: A Generic Genome Profiling Technology on Open Platforms.” *Methods in Molecular Biology (Clifton*, N.J.) 888: 67–89.

Lachmuth, Susanne, Thibaut Capblancq, Stephen R. Keller, and Matthew C. Fitzpatrick. 2023. “Assessing Uncertainty in Genomic Offset Forecasts from Landscape Genomic Models (and Implications for Restoration and Assisted Migration).” Frontiers in Ecology and Evolution 11 (1155783): 1155783.

Lachmuth, Susanne, Thibaut Capblancq, Anoob Prakash, Stephen R. Keller, and Matthew C. Fitzpatrick. 2024. “Novel Genomic Offset Metrics Integrate Local Adaptation into Habitat Suitability Forecasts and Inform Assisted Migration.” Ecological Monographs 94 (1): e1593.

Láruson, Áki Jarl, Matthew C. Fitzpatrick, Stephen R. Keller, Benjamin C. Haller, and Katie E. Lotterhos. 2022. “Seeing the Forest for the Trees: Assessing Genetic Offset Predictions from Gradient Forest.” Evolutionary Applications 15 (3): 403–416.

Lavoie, Juliette, Pascal Bourgault, Trevor James Smith, et al. 2024. “An Ensemble of Bias-Adjusted CMIP6 Climate Simulations Based on a High-Resolution North American Reanalysis.” Scientific Data 11 (1): 64.

Lesven, Jonathan A., Milva Druguet Dayras, Jonathan Cazabonne, et al. 2024. “Future Impacts of Climate Change on Black Spruce Growth and Mortality: Review and Challenges.” Environmental Reviews 32 (2): 214–230.

Lind, Brandon M., Rafael Candido-Ribeiro, Pooja Singh, et al. 2024. “How Useful Are Genomic Data for Predicting Maladaptation to Future Climate?” Global Change Biology 30 (4): e17227.

Lind, Brandon M., and Katie E. Lotterhos. 2025a. “A Comparison of Genomic Forecasts Based on Genotypes versus Allele Frequencies.” The American Naturalist, no. 739098 (October). 10.1086/739098.

Lind, Brandon M., and Katie E. Lotterhos. 2025b. “The Accuracy of Predicting Maladaptation to New Environments with Genomic Data.” Molecular Ecology Resources 25 (4): e14008.

Liu, Qiuyu, Changhui Peng, Robert Schneider, Dominic Cyr, Nate G. McDowell, and Daniel Kneeshaw. 2023. “Drought-Induced Increase in Tree Mortality and Corresponding Decrease in the Carbon Sink Capacity of Canada’s Boreal Forests from 1970 to 2020.” Global Change Biology 29 (8): 2274–2285.

Lloret, Francisco, Eric G. Keeling, and Anna Sala. 2011. “Components of Tree Resilience: Effects of Successive Low-Growth Episodes in Old Ponderosa Pine Forests.” Oikos (Copenhagen, Denmark) 120 (12): 1909–1920.

Lortie, Christopher J., and José L. Hierro. 2022. “A Synthesis of Local Adaptation to Climate through Reciprocal Common Gardens.” The Journal of Ecology 110 (5): 1015–1021.

Marchand, William, Claire Depardieu, Elizabeth M. Campbell, Jean Bousquet, and Martin P. Girardin. 2025. “Long-Term Temporal Divergence in Post-Drought Resilience Decline between Deciduous and Evergreen Tree Species.” Global Change Biology 31 (7): e70330.

Marchand, William, Martin P. Girardin, Henrik Hartmann, Mathieu Lévesque, Sylvie Gauthier, and Yves Bergeron. 2021. “Contrasting Life-History Traits of Black Spruce and Jack Pine Influence Their Physiological Response to Drought and Growth Recovery in Northeastern Boreal Canada.” The Science of the Total Environment 794 (148514): 148514.

Marquis, Benjamin, Yves Bergeron, Daniel Houle, Martin Leduc, and Sergio Rossi. 2022. “Variability in Frost Occurrence under Climate Change and Consequent Risk of Damage to Trees of Western Quebec, Canada.” Scientific Reports 12 (1): 7220.

Mijangos, Jose Luis, Bernd Gruber, Oliver Berry, Carlo Pacioni, and Arthur Georges. 2022. “*dartR* v2: An Accessible Genetic Analysis Platform for Conservation, Ecology and Agriculture.” *Methods in Ecology and Evolution / British Ecological Society*, ahead of print, July 11. 10.1111/2041-210x.13918.

Morgenstern, E. K. 1978. “Range-Wide Genetic Variation of Black Spruce.” Canadian Journal of Forest Research 8 (4): 463–473.

Mura, Claudio, Valentina Buttò, Roberto Silvestro, et al. 2022. “The Early Bud Gets the Cold: Diverging Spring Phenology Drives Exposure to Late Frost in a Picea Mariana [(Mill.) BSP] Common Garden.” Physiologia Plantarum 174 (6): e13798.

Oksanen, J., G. Simpson, F. Blanchet, et al. 2025. Vegan: Community Ecology Package. Version R package version 2.8-0. https://vegandevs.github.io/vegan/.

Oleksyn, J., J. Modrzýnski, M. G. Tjoelker, R. Zytkowiak, P. B. Reich, and P. Karolewski. 1998. “Growth and Physiology of *Picea Abies* Populations from Elevational Transects: Common Garden Evidence for Altitudinal Ecotypes and Cold Adaptation.” Functional Ecology 12 (4): 573–590.

Patsiou, Theofania S., Tatiana A. Shestakova, Tamir Klein, et al. 2020. “Intraspecific Responses to Climate Reveal Nonintuitive Warming Impacts on a Widespread Thermophilic Conifer.” The New Phytologist 228 (2): 525–540.

Pebesma, E. 2018. “Simple Features for R: Standardized Support for Spatial Vector Data.” The R Journal 10 (1): 439.

Perry, Daniel J., and Jean Bousquet. 2001. “Genetic Diversity and Mating System of Post-Fire and Post-Harvest Black Spruce: An Investigation Using Codominant Sequence-Tagged-Site (STS) Markers.” Canadian Journal of Forest Research 31 (1): 32–40.

Prasad, Anantha, John Pedlar, Matt Peters, et al. 2020. “Combining US and Canadian Forest Inventories to Assess Habitat Suitability and Migration Potential of 25 Tree Species under Climate Change.” Diversity & Distributions 26 (9): 1142–1159.

Price, David T., R. I. Alfaro, K. J. Brown, et al. 2013. “Anticipating the Consequences of Climate Change for Canada’s Boreal Forest Ecosystems.” Environmental Reviews 21 (4): 322–365.

Prunier, Julien, Sébastien Gérardi, Jérôme Laroche, Jean Beaulieu, and Jean Bousquet. 2012. “Parallel and Lineage-Specific Molecular Adaptation to Climate in Boreal Black Spruce: LINEAGE-SPECIFIC ADAPTIVE SNP IN BLACK SPRUCE.” Molecular Ecology 21 (17): 4270–4286.

R Core Team. 2024. “R: A Language and Environment for Statistical Computing.” Preprint, R Foundation for Statistical Computing. https://www.R-project.org/.

R Core Team. 2026. “R: A Language and Environment for Statistical Computing.” Preprint, R Foundation for Statistical Computing. https://www.R-project.org/.

Rellstab, Christian, Benjamin Dauphin, and Moises Exposito-Alonso. 2021. “Prospects and Limitations of Genomic Offset in Conservation Management.” Evolutionary Applications 14 (5): 1202–1212.

Rellstab, Christian, Stefan Zoller, Lorenz Walthert, et al. 2016. “Signatures of Local Adaptation in Candidate Genes of Oaks (Quercus Spp.) with Respect to Present and Future Climatic Conditions.” Molecular Ecology 25 (23): 5907–5924.

Rhoné, Bénédicte, Dimitri Defrance, Cécile Berthouly-Salazar, et al. 2020. “Pearl Millet Genomic Vulnerability to Climate Change in West Africa Highlights the Need for Regional Collaboration.” Nature Communications 11 (1): 5274.

Robert, Etienne, Patrick Lenz, Yves Bergeron, et al. 2024. “Future Carbon Sequestration Potential in a Widespread Transcontinental Boreal Tree Species: Standing Genetic Variation Matters!” Global Change Biology 30 (6): e17347.

Robert, Etienne, Patrick Lenz, Yves Bergeron, Nathalie Isabel, and Martin P. Girardin. 2025. “Black Spruce Growth under Climate Extremes: Genetic Insights for Managing a Key Resource Production Species.” Forest Ecology and Management 597 (123129): 123129.

Rossi, Sergio. 2015. “Local Adaptations and Climate Change: Converging Sensitivity of Bud Break in Black Spruce Provenances.” International Journal of Biometeorology 59 (7): 827– 835.

Rossi, Sergio, and Jean Bousquet. 2014. “The Bud Break Process and Its Variation among Local Populations of Boreal Black Spruce.” Frontiers in Plant Science 5 (October): 574.

Schmidt, Thomas L., Moshe-Elijah Jasper, Andrew R. Weeks, and Ary A. Hoffmann. 2021. “Unbiased Population Heterozygosity Estimates from Genome wide Sequence Data.” Methods in Ecology and Evolution 12 (10): 1888–1898.

Shafer, Aaron B. A., Catherine I. Cullingham, Steeve D. Côté, and David W. Coltman. 2010. “Of Glaciers and Refugia: A Decade of Study Sheds New Light on the Phylogeography of Northwestern North America.” Molecular Ecology 19 (21): 4589–4621.

Sniderhan, Anastasia E., Steven D. Mamet, and Jennifer L. Baltzer. 2021. “Non-Uniform Growth Dynamics of a Dominant Boreal Tree Species (Picea Mariana) in the Face of Rapid Climate Change.” Canadian Journal of Forest Research 51 (4): 565–572.

Urban, M. C., G. Bocedi, A. P. Hendry, et al. 2016. “Improving the Forecast for Biodiversity under Climate Change.” Science (New York, N.Y.) 353 (6304): aad8466–aad8466.

Vallejos, Ronny, Felipe Osorio, and Moreno Bevilacqua. 2020. Spatial Relationships between Two Georeferenced Variables: With Applications in R. 2020th ed. Springer Nature.

Verrico, Brittany M., Thibaut Capblancq, Matthew C. Fitzpatrick, and Stephen R. Keller. 2025. “Reciprocal Evaluation of Genomic Offset Predictions of Climate Maladaptation with Independent Empirical Datasets.” The American Naturalist, no. 739111 (October). 10.1086/739111.

Walker, Xanthe J., Michelle C. Mack, and Jill F. Johnstone. 2015. “Stable Carbon Isotope Analysis Reveals Widespread Drought Stress in Boreal Black Spruce Forests.” Global Change Biology 21 (8): 3102–3113.

Zheng, Xiuwen, David Levine, Jess Shen, Stephanie M. Gogarten, Cathy Laurie, and Bruce S. Weir. 2012. “A High-Performance Computing Toolset for Relatedness and Principal Component Analysis of SNP Data.” Bioinformatics (Oxford, England) 28 (24): 3326–3328.

