## Supplementary Figures, Supplementary Tables for "Testing genomic offset with common gardens in genetically structured black spruce (*Picea mariana)*"

Supplementary Material

Supplementary Figures 1-27

Supplementary Tables 1-8

Supplementary Datasets (xlsx tables):

- Supplementary Dataset 1. Tree sample information.
- Supplementary Dataset 2. Sampling coordinates of red spruce populations.
- Supplementary Dataset 3. Past climate normals for 43 climate variables.
- Supplementary Dataset 4. Population means and standard errors of measured phenotypic traits.

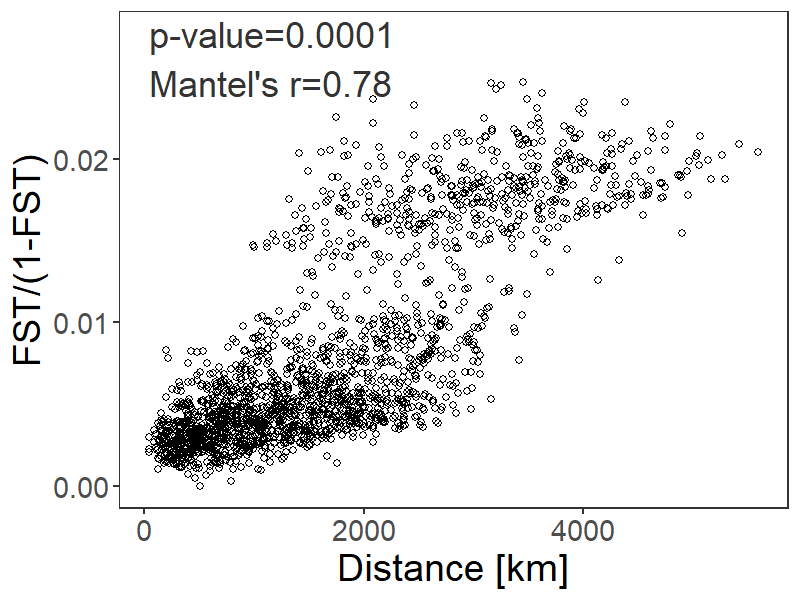

Supplementary Figure 1. Patterns of isolation by distance across species distribution. Each point represents a pair of populations.

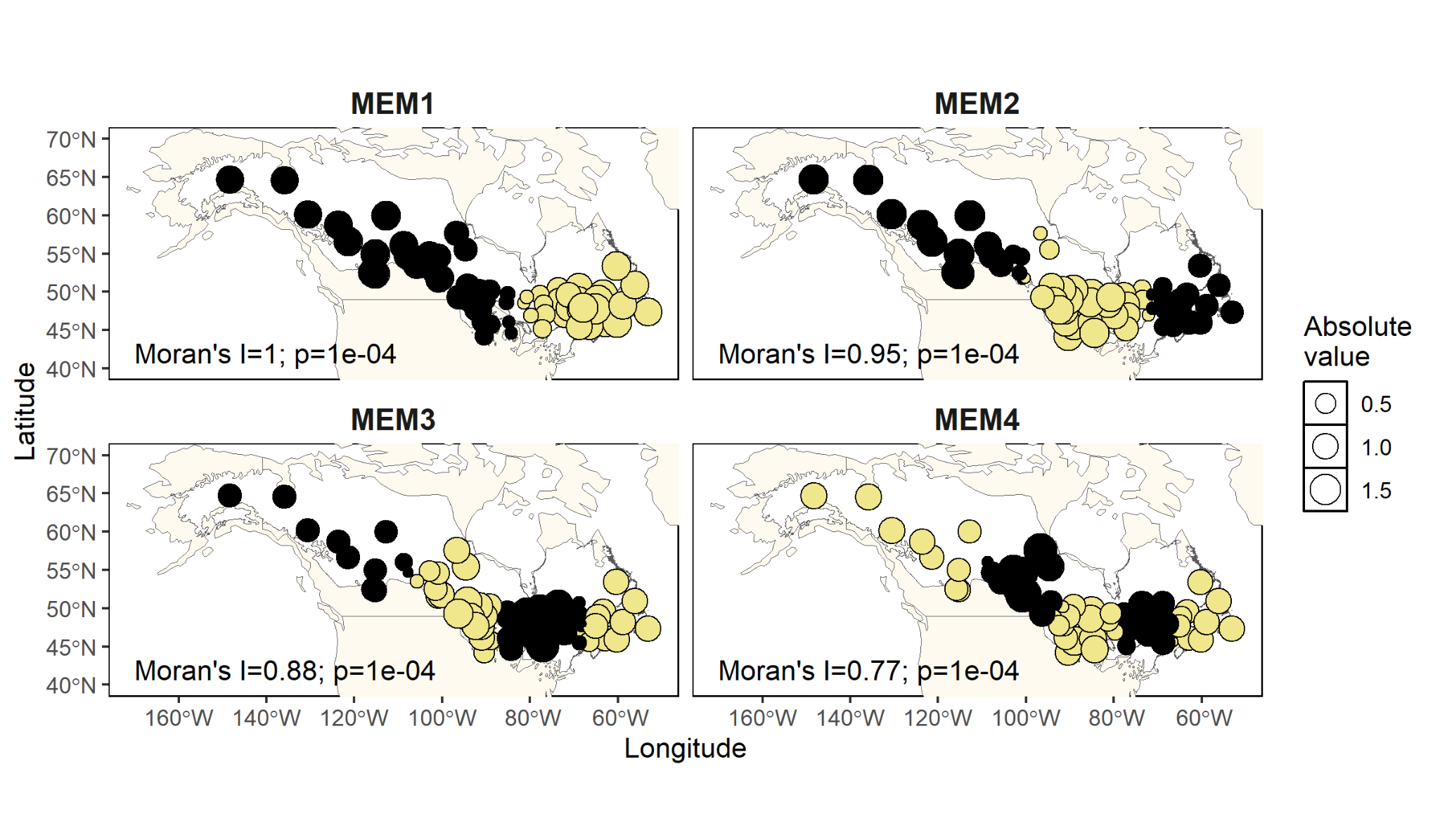

Supplementary Figure 2. Four major axes of spatial autocorrelation. Each circle represents black spruce population. The two colors indicate either positive or negative values of the corresponding axes with size scaled by its absolute value.

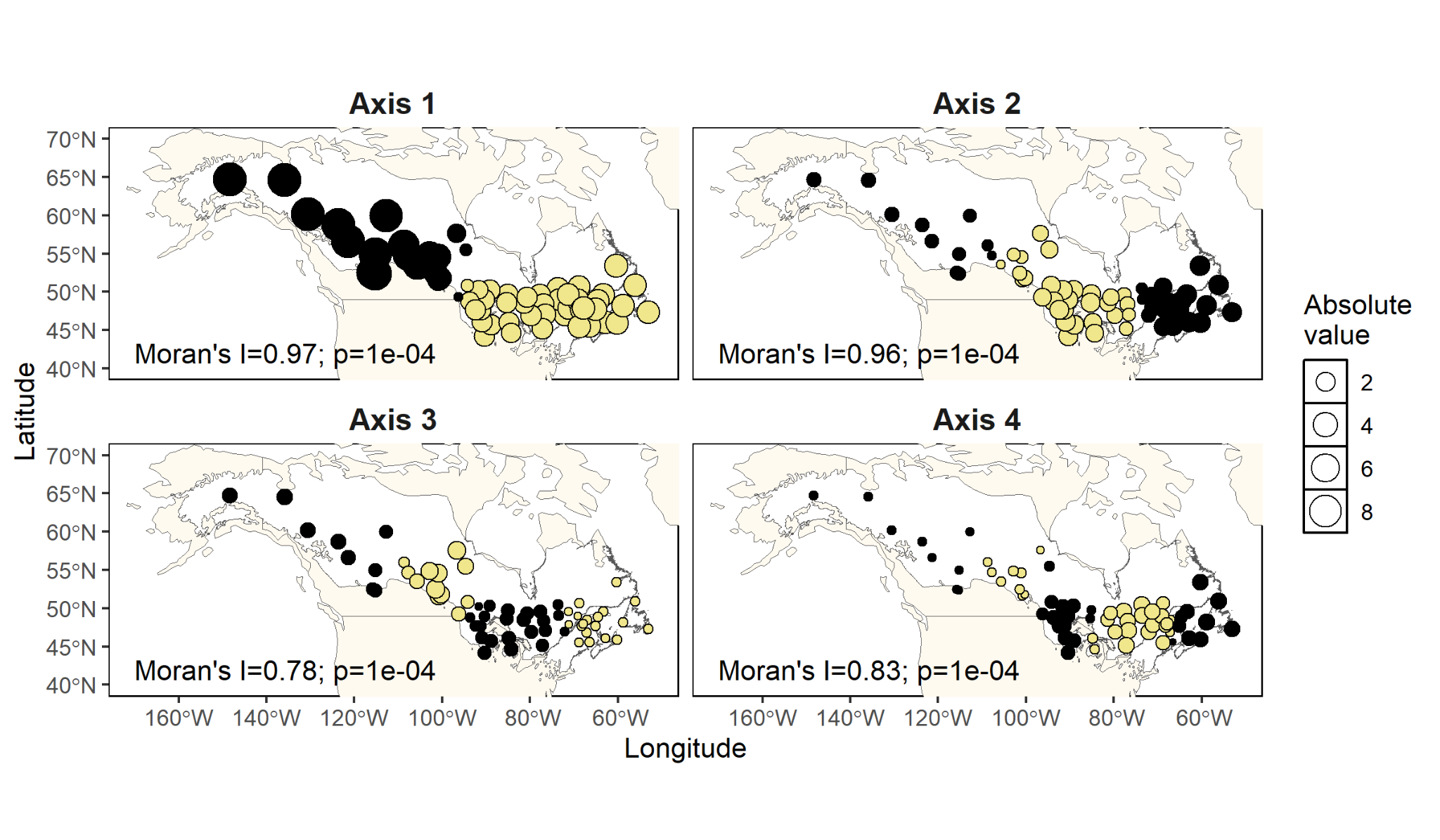

Supplementary Figure 3. Spatial visualization of sPCA four main axes. Each circle represents black spruce population. The two colors indicate either positive or negative values of the corresponding axes with size scaled by its absolute value.

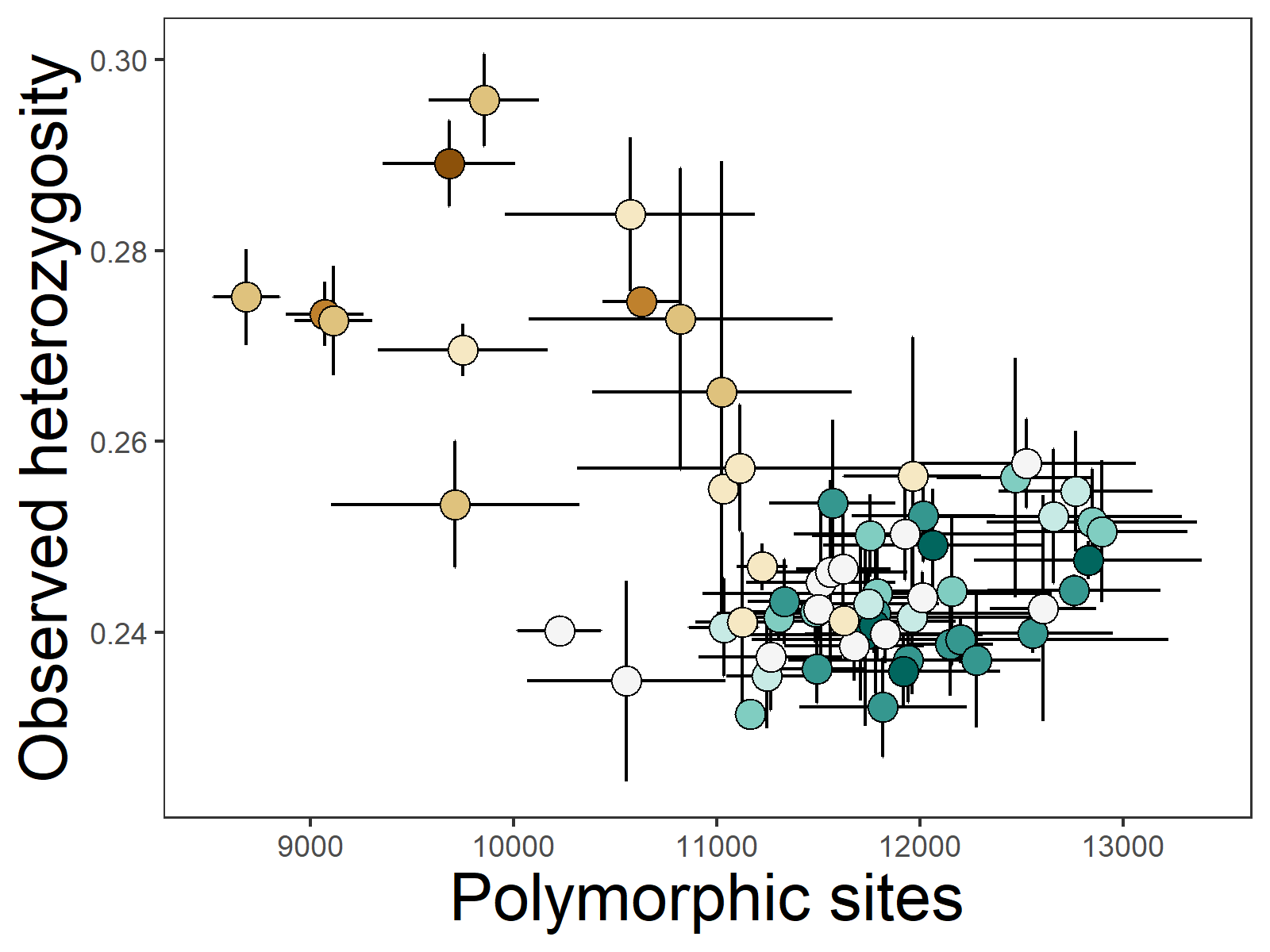

Supplementary Figure 4. Observed heterozygosity versus number of polymorphic sites across populations of black spruce. Observed heterozygosity was calculated from five random subsamples of seven individuals per population, after excluding monomorphic sites within each subsample. Colors correspond to geographical regions depicted in Figure 1A.

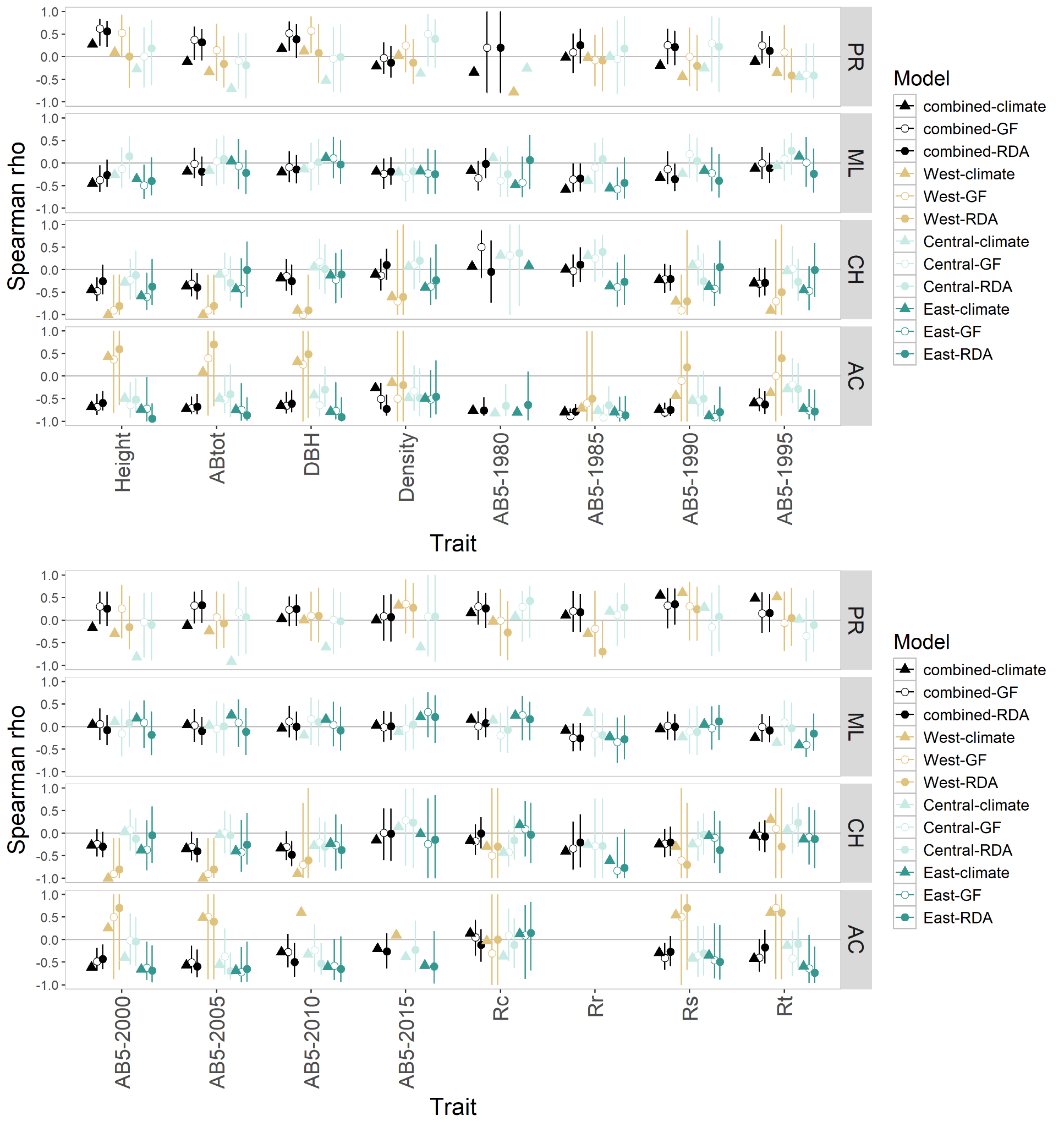

Supplementary Figure 5. Performance of Gradient Forest (GF) and redundancy analysis (RDA) models validated against all fitness traits and estimated for all populations (“combined”) or individual genetic clusters. Triangles indicate Spearman rho calculated between trait value and climate transfer distance.

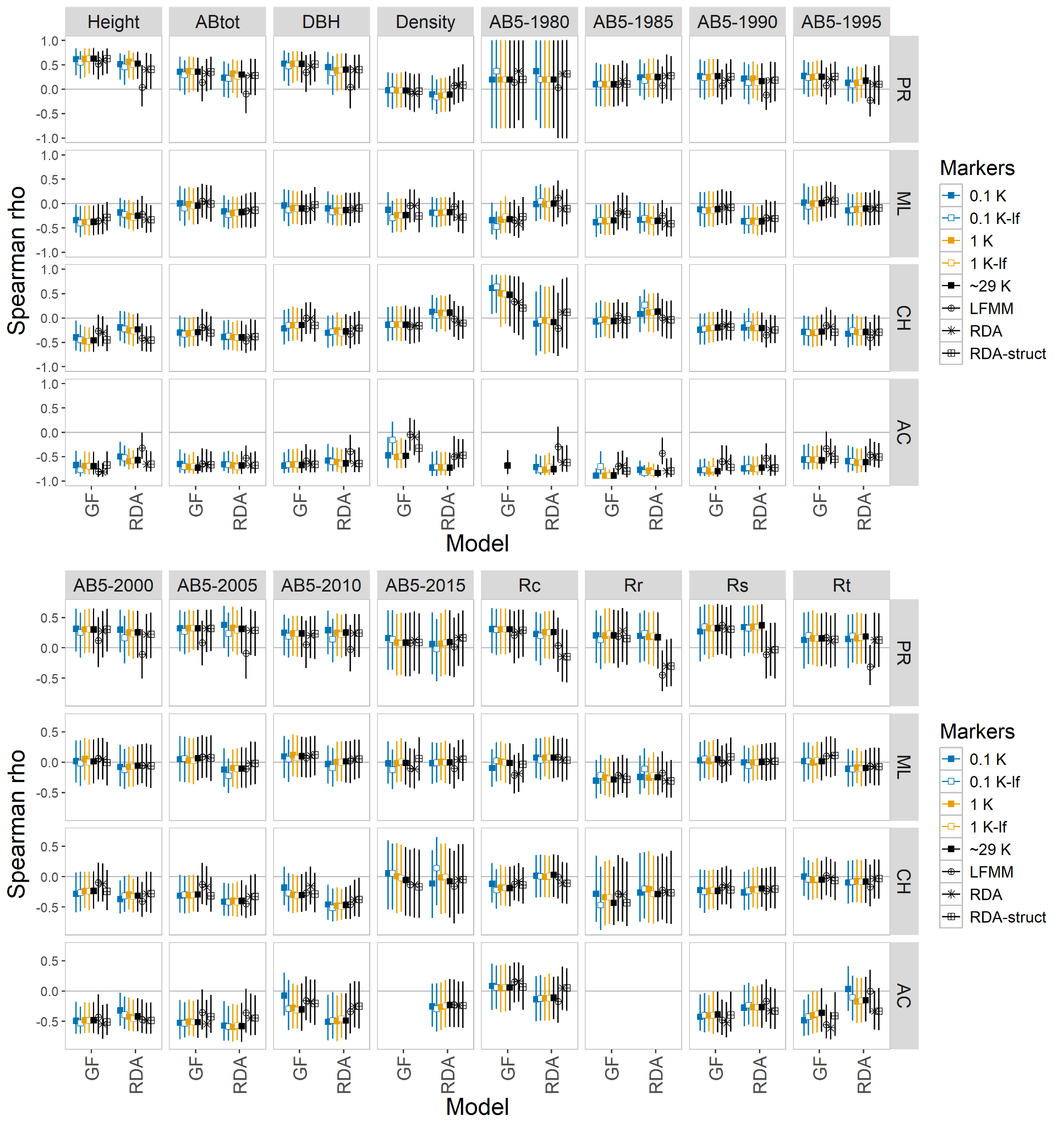

Supplementary Figure 6. Comparison of predictive performance of Gradient Forest (GF) and redundancy analysis (RDA) models trained on eight different marker types and estimated for all populations. The marker types included subsets of 100 (0.1K), 1000 (1K) or all available markers (~29K), subsets including low-frequency variants (0.1K-lf, 1K-lf), and markers found to be in association with climate (LFMM, RDA, RDA-struct).

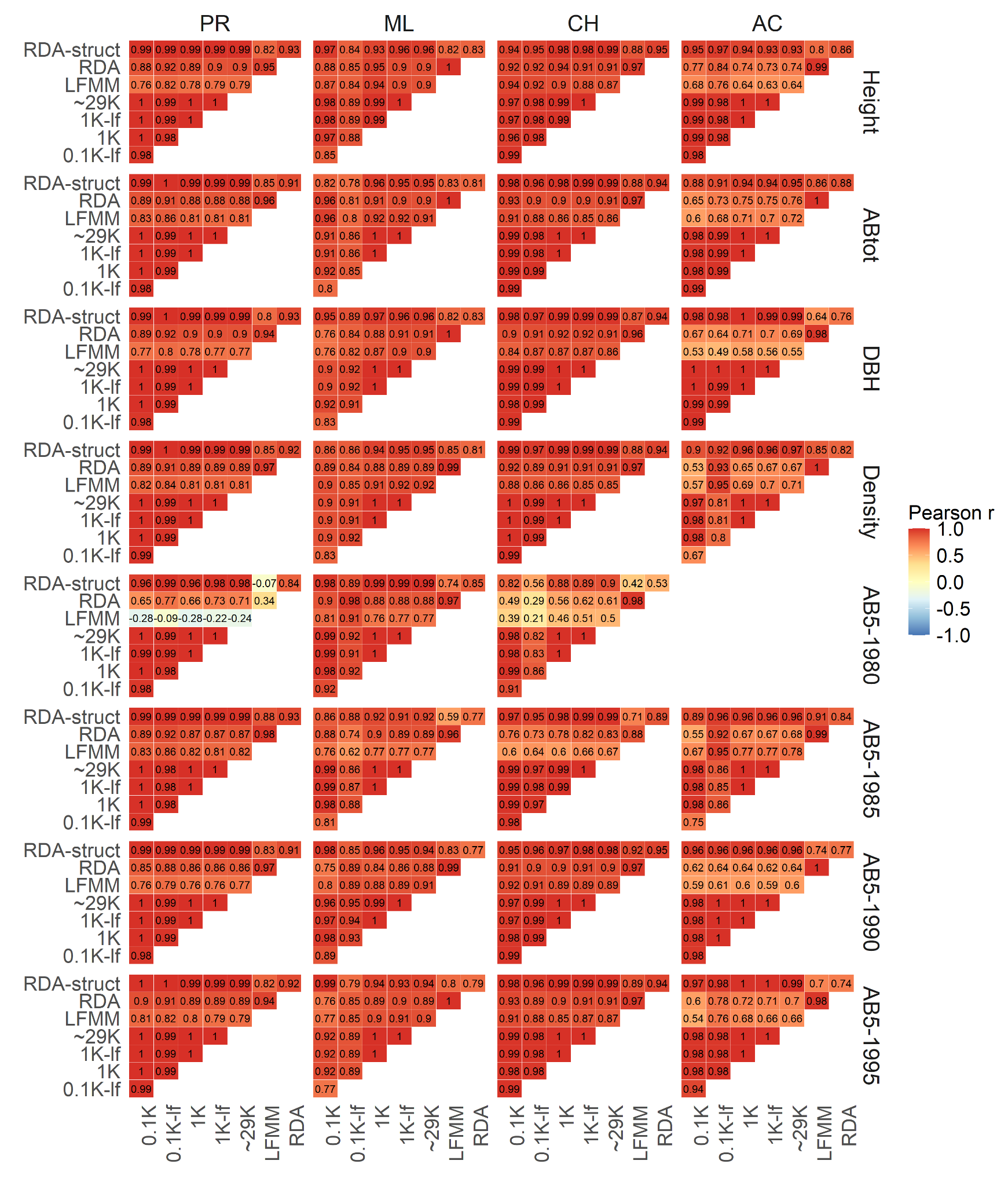

Supplementary Figure 7. Matrices of pairwise correlations between genomic offsets, estimated via Gradient Forest models trained on different sets of markers validated for the first eight fitness traits. The marker types included subsets of 100 (0.1K), 1000 (1K) or all available markers (~29K), subsets including low-frequency variants (0.1K-lf, 1K-lf), and markers found to be in association with climate (LFMM, RDA, RDA-struct).

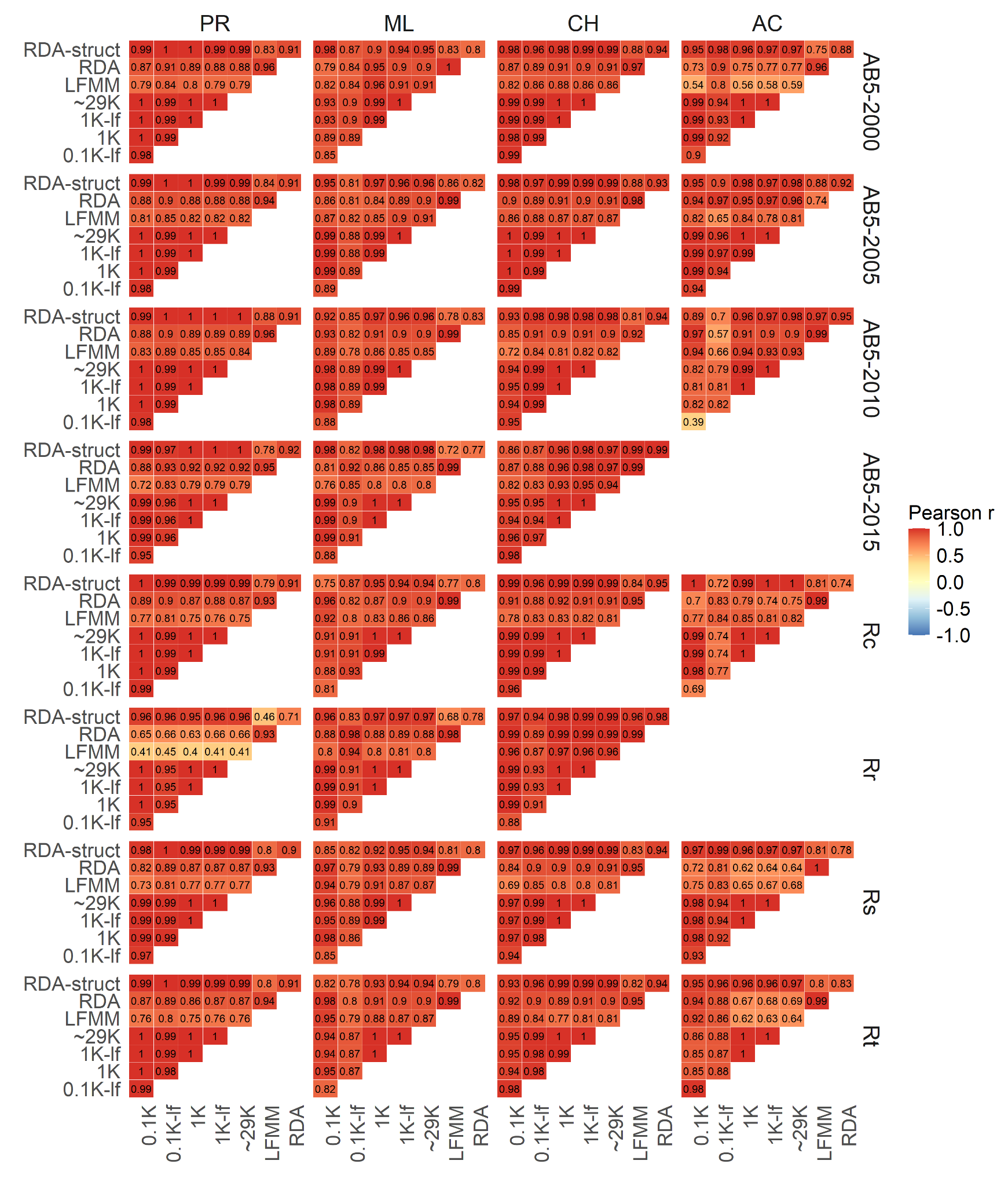

Supplementary Figure 8. Matrices of pairwise correlations between genomic offsets, estimated via Gradient Forest models trained on different sets of markers validated for the last eight fitness traits. The marker types included subsets of 100 (0.1K), 1000 (1K) or all available markers (~29K), subsets including low-frequency variants (0.1K-lf, 1K-lf), and markers found to be in association with climate (LFMM, RDA, RDA-struct).

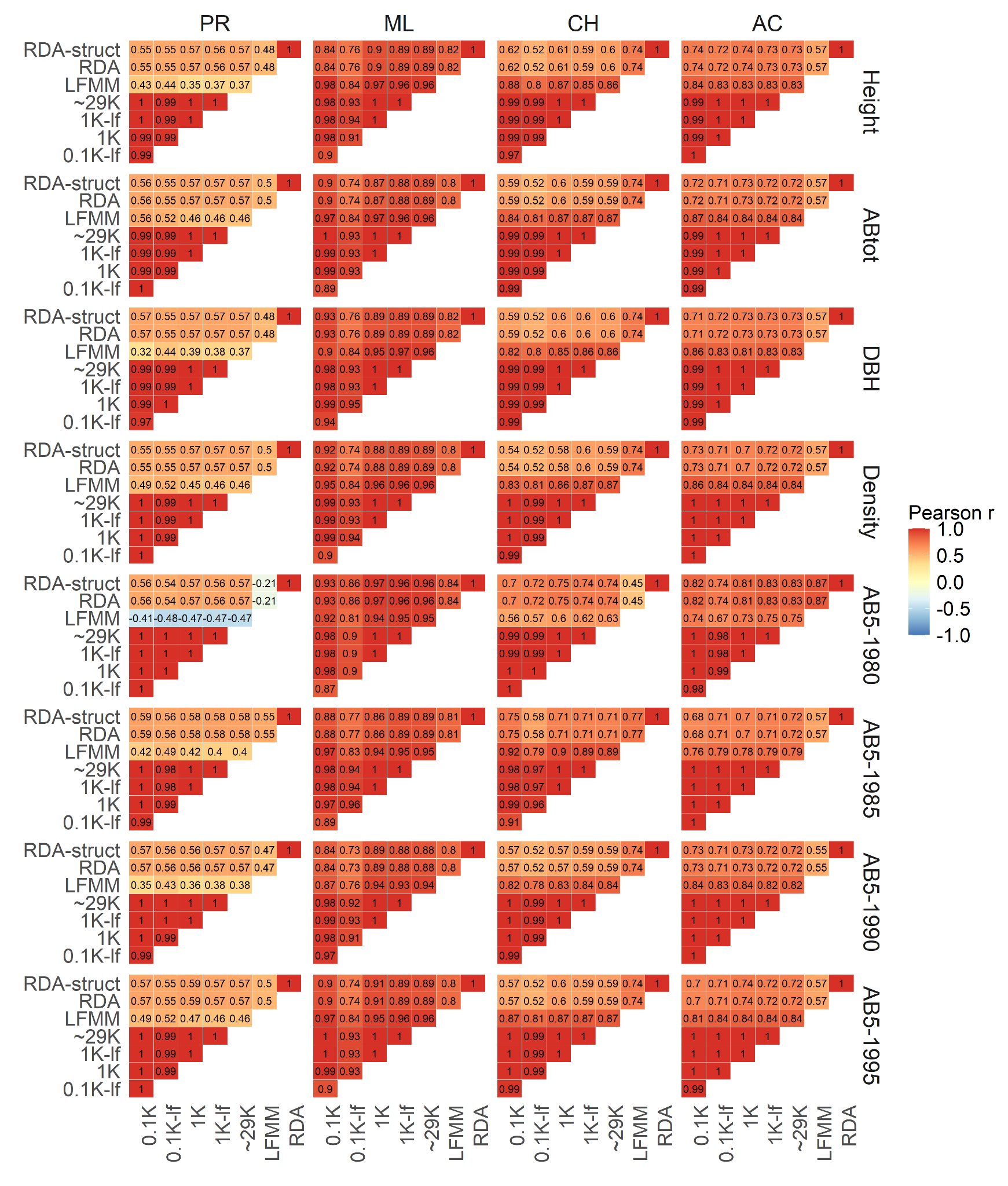

Supplementary Figure 9. Matrices of pairwise correlations between genomic offsets, estimated via RDA models trained on different sets of markers validated for the first eight fitness traits. The marker types included subsets of 100 (0.1K), 1000 (1K) or all available markers (~29K), subsets including low-frequency variants (0.1K-lf, 1K-lf), and markers found to be in association with climate (LFMM, RDA, RDA-struct).

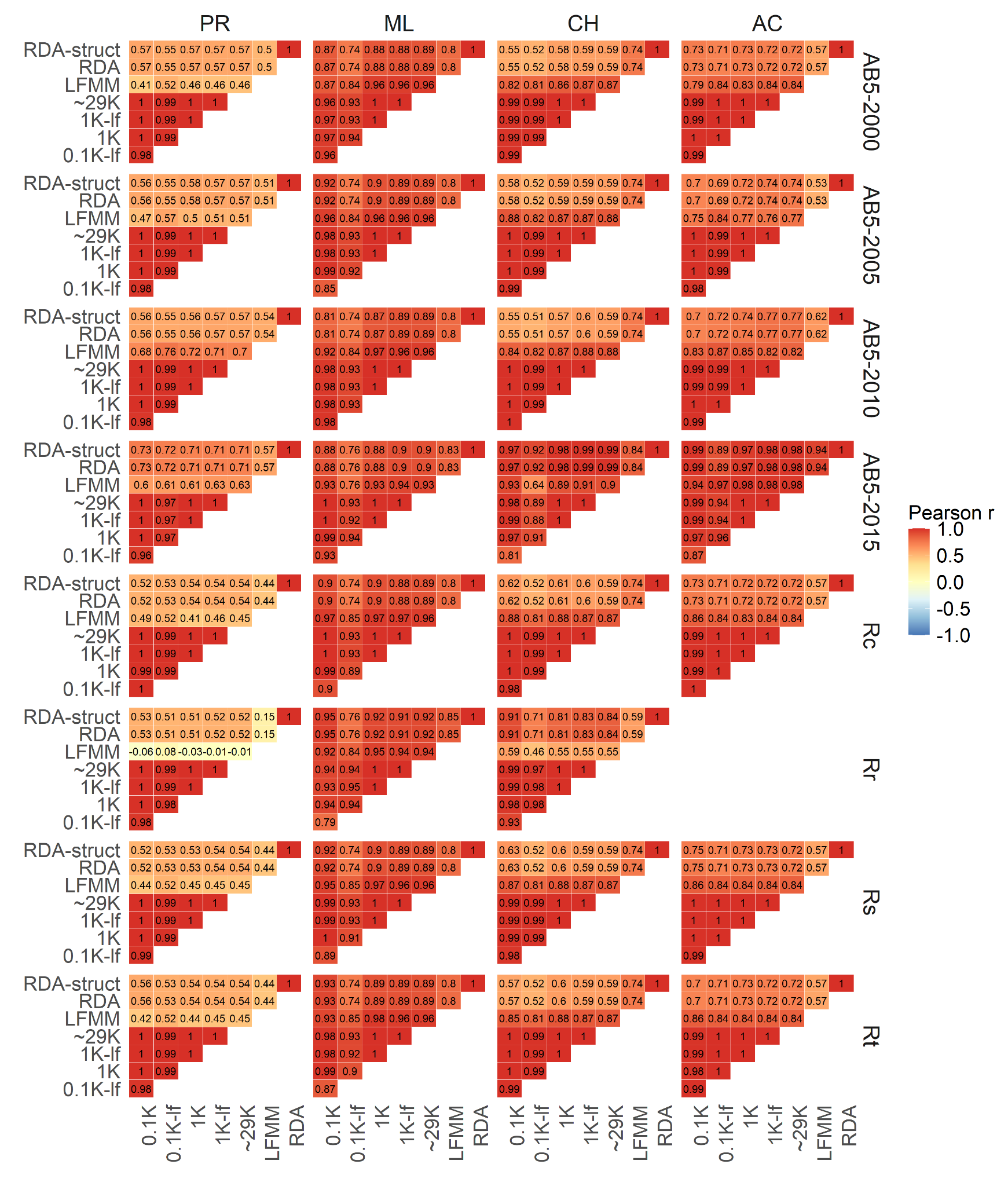

Supplementary Figure 10. Matrices of pairwise correlations between genomic offsets, estimated via RDA models trained on different sets of markers validated for the last eight fitness traits. The marker types included subsets of 100 (0.1K), 1000 (1K) or all available markers (~29K), subsets including low-frequency variants (0.1K-lf, 1K-lf), and markers found to be in association with climate (LFMM, RDA, RDA-struct).

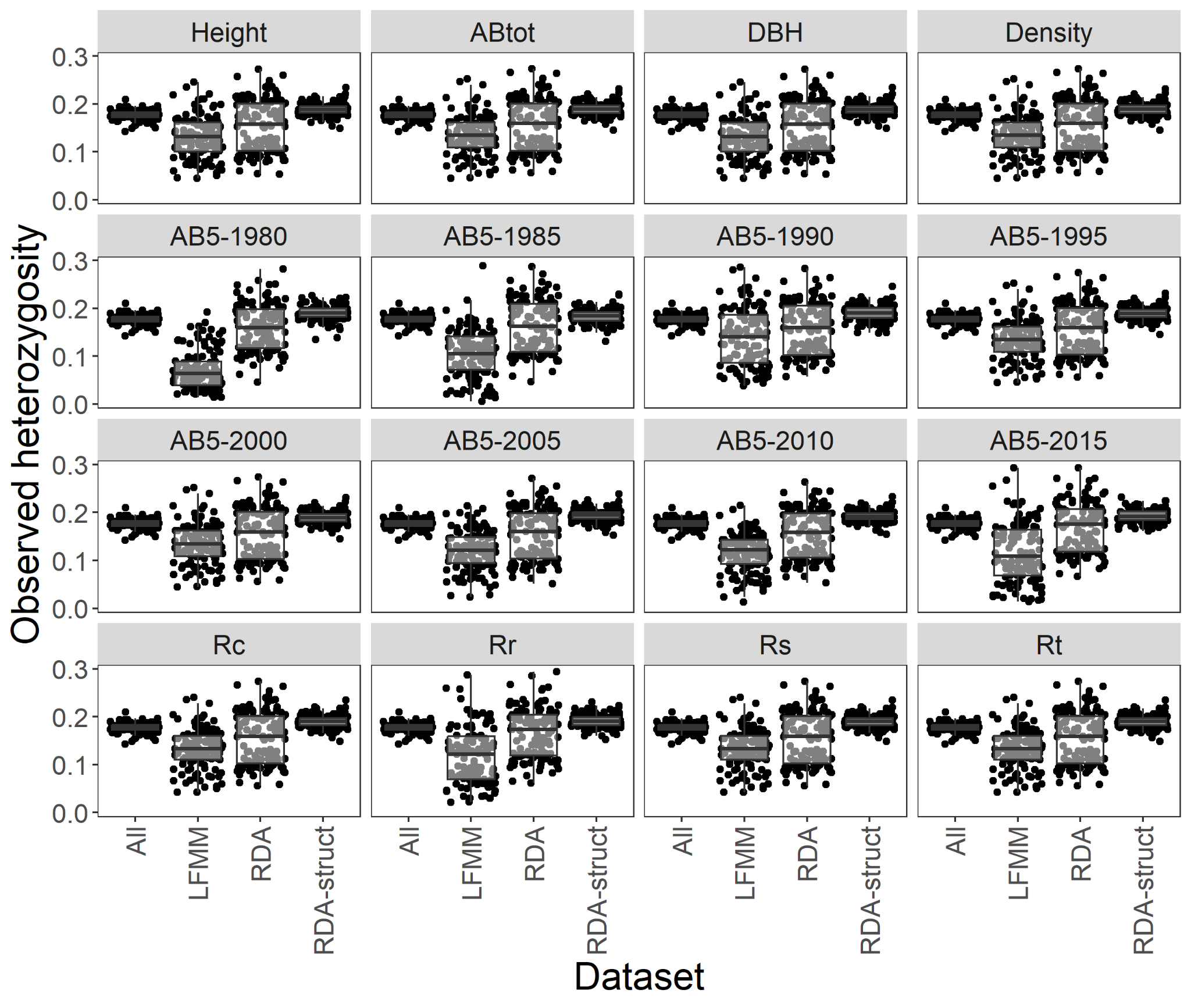

Supplementary Figure 11. Observed heterozygosity calculated per population using selected marker sets: all variants (All), LFMM outliers (LFMM), RDA outliers (RDA) and RDA outliers corrected for genetic structure (RDA-struct). Minor allele frequency cutoff of 0.05 was applied to all datasets.

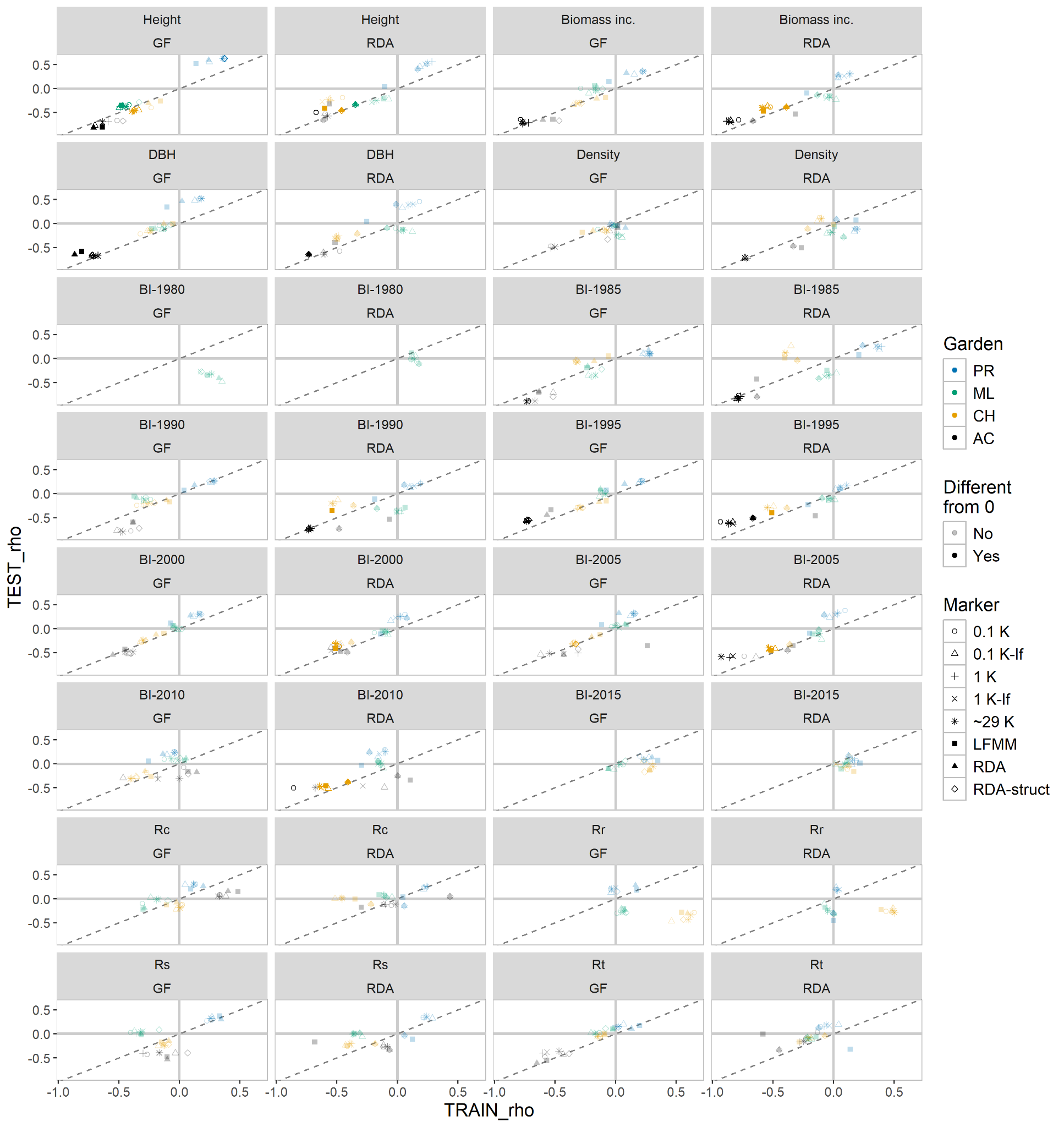

Supplementary Figure 12. Comparison of predictive performance between models trained on training (axis x) and test datasets (axis y). Models were trained using Gradient Forest (GF) or redundancy analysis (RDA). Solid colors indicate model performances significantly different from zero. The marker types included subsets of 100 (0.1K), 1000 (1K) or all available markers (~29K), subsets including low-frequency variants (0.1K-lf, 1K-lf), and markers found to be in association with climate (LFMM, RDA, RDA-struct).

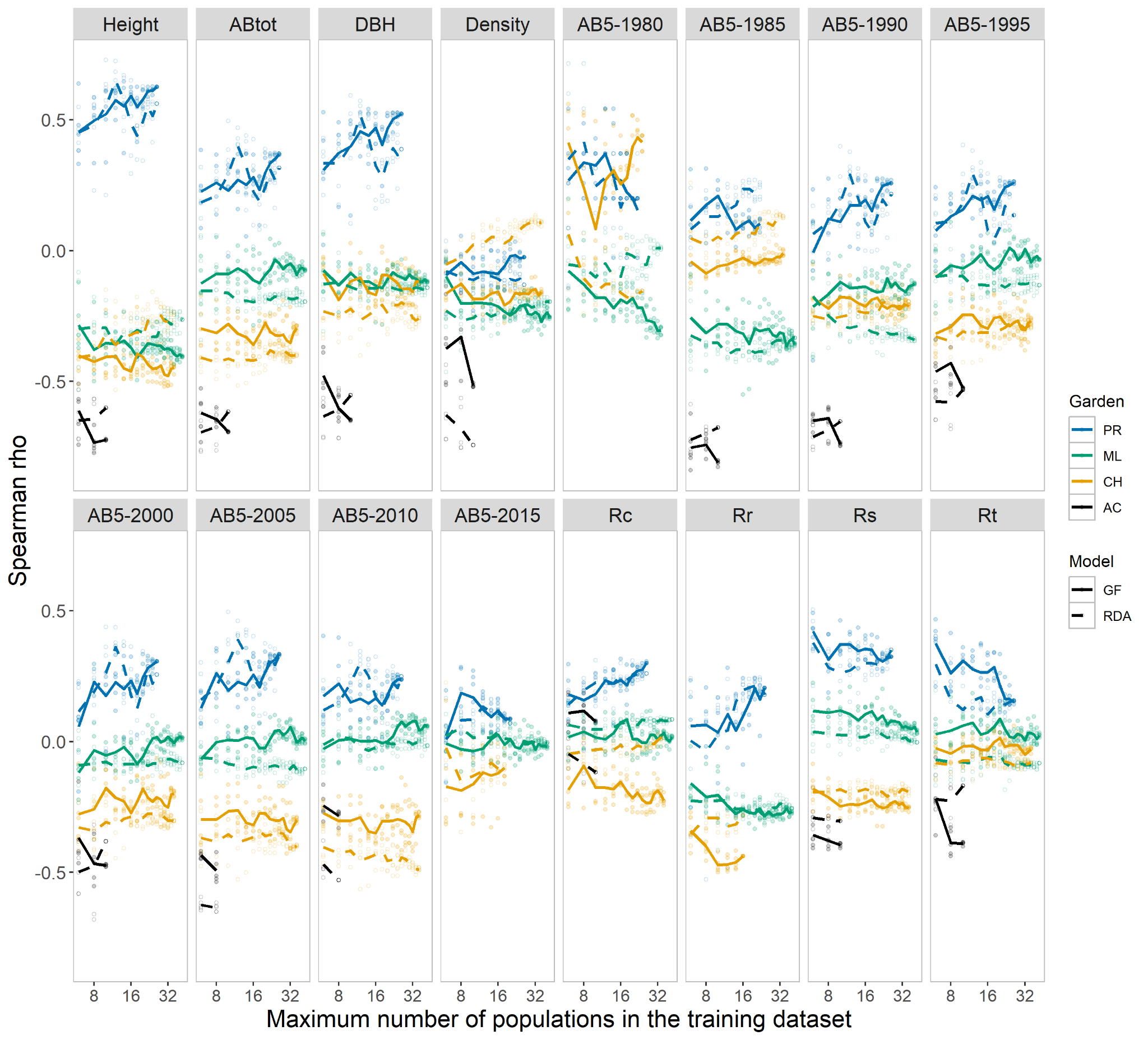

Supplementary Figure 13. Testing the impact of the number of populations in the training dataset on model performance. Points indicate five random draws of n populations (axis x). Lines connect the mean values for each population number. Gradient Forest (GF) and redundancy analysis (RDA) models were trained on a random set of 1000 markers.

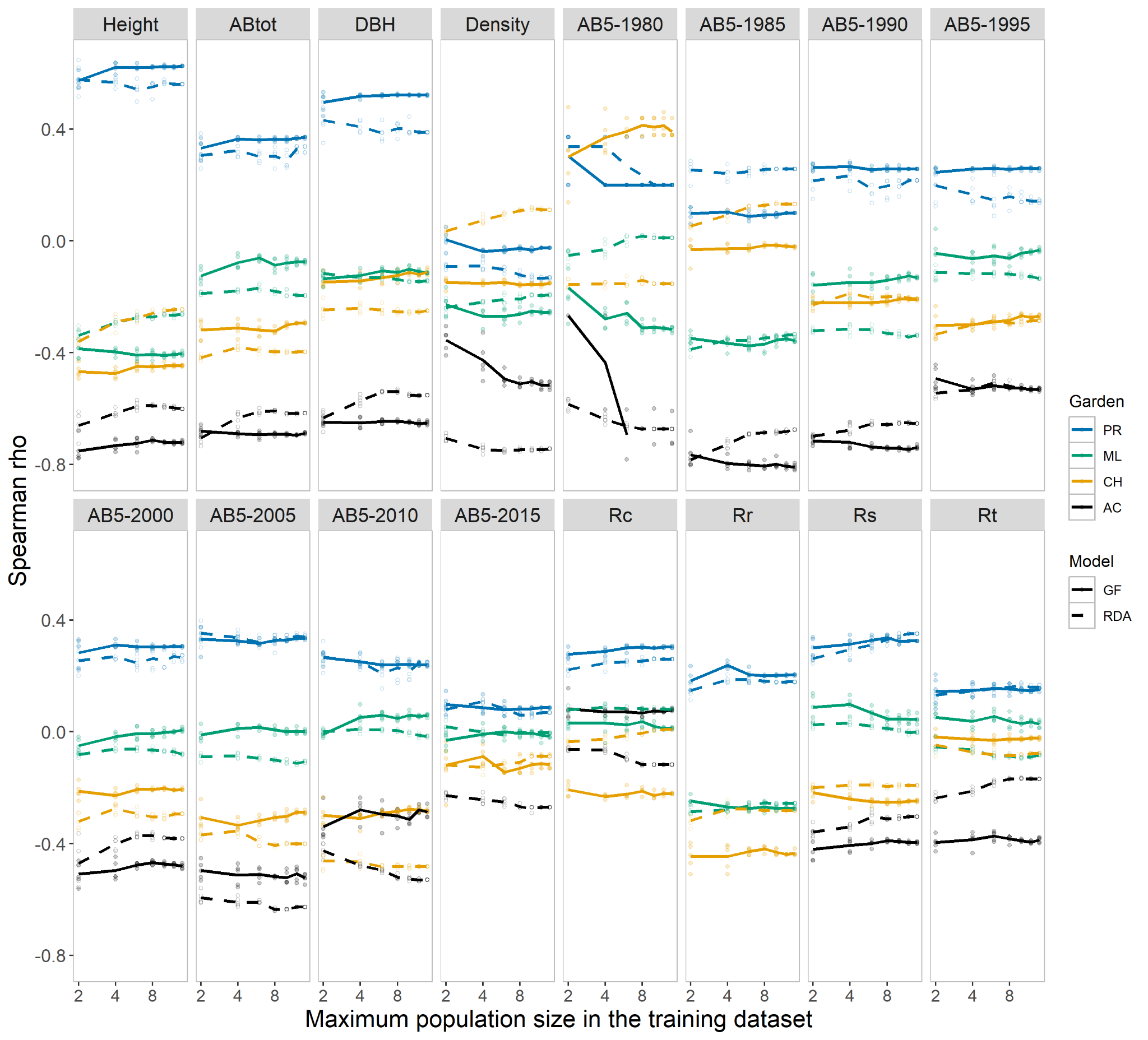

Supplementary Figure 14. Testing the impact of the population size in the training dataset on model performance. Points indicate ten random draws of n genotypes (axis x). Lines connect the mean values for each population size. Gradient Forest (GF) and redundancy analysis (RDA) models were trained on a random set of 1000 markers.

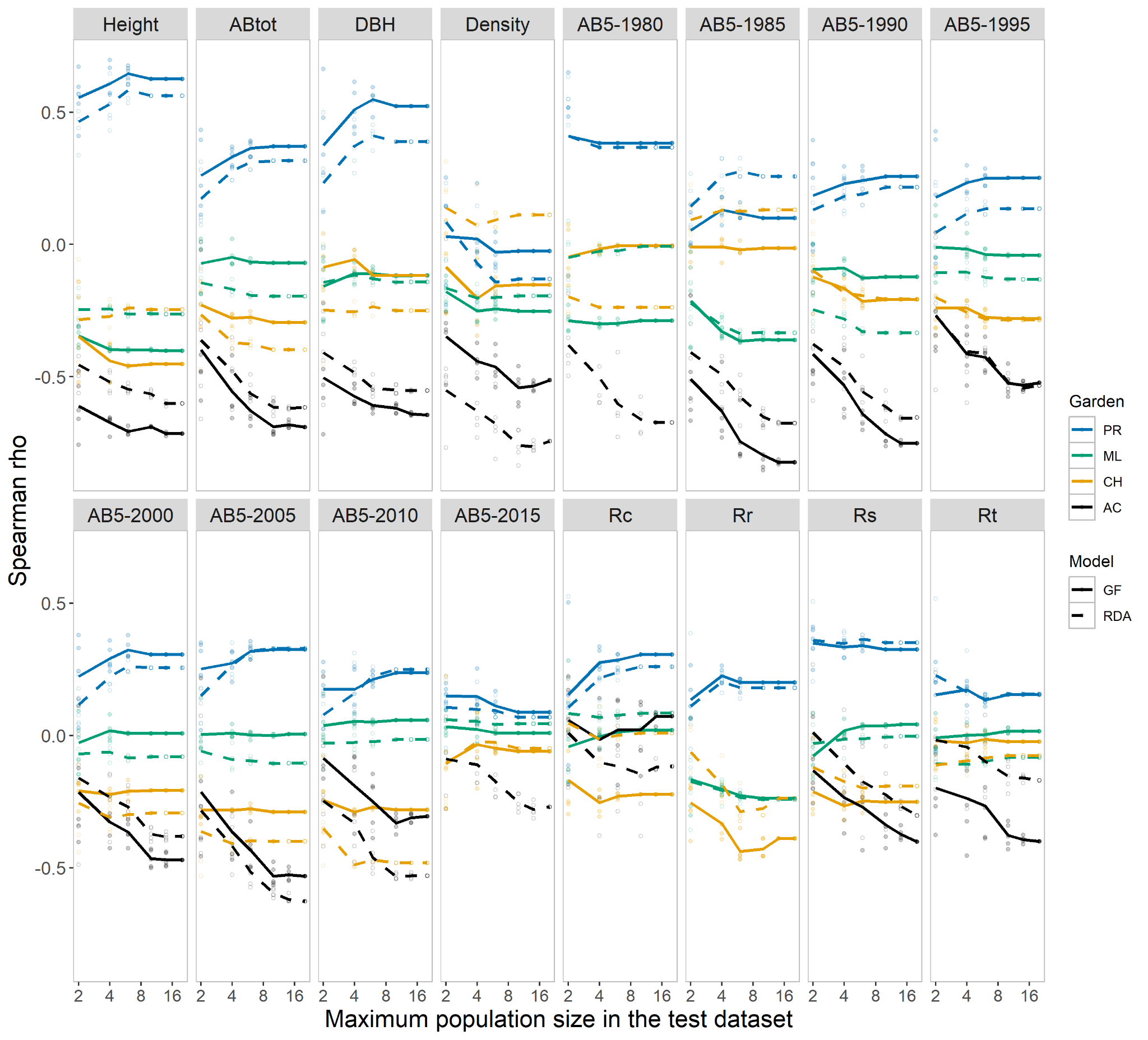

Supplementary Figure 15. Testing the impact of the population size in the test dataset on model performance. Points indicate ten random draws of individual trees of a given number per population (axis x). Lines connect the mean values for each population size. Gradient Forest (GF) and redundancy analysis (RDA) models were trained on a random set of 1000 markers.

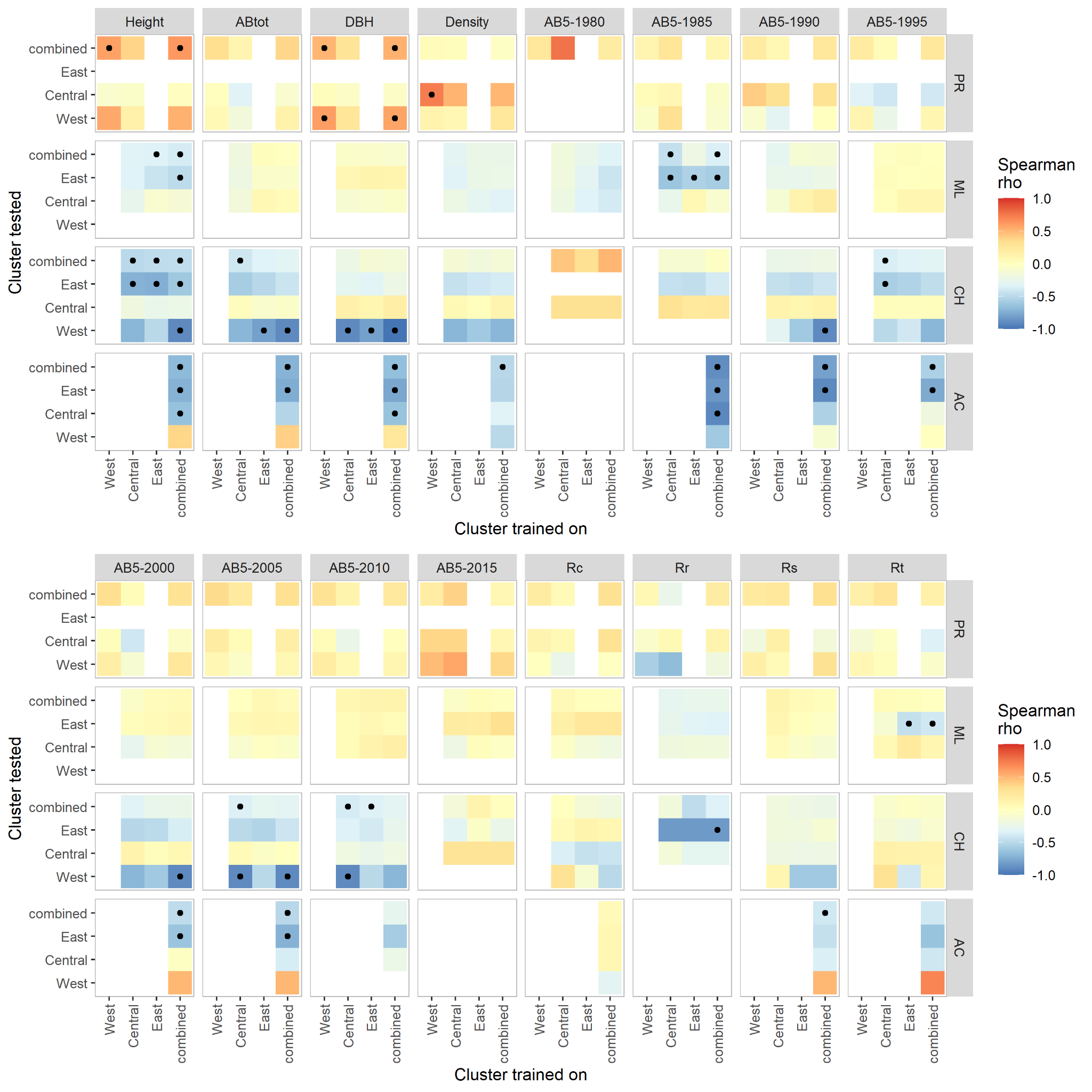

Supplementary Figure 16. Gradient Forest model performances when trained on individual genetic clusters or combined datasets and three climate PCs in predicting fitness traits of the same or other genetic clusters. Dots indicate correlations with confidence intervals not overlapping zero. Models were trained on a random set of 1000 markers.

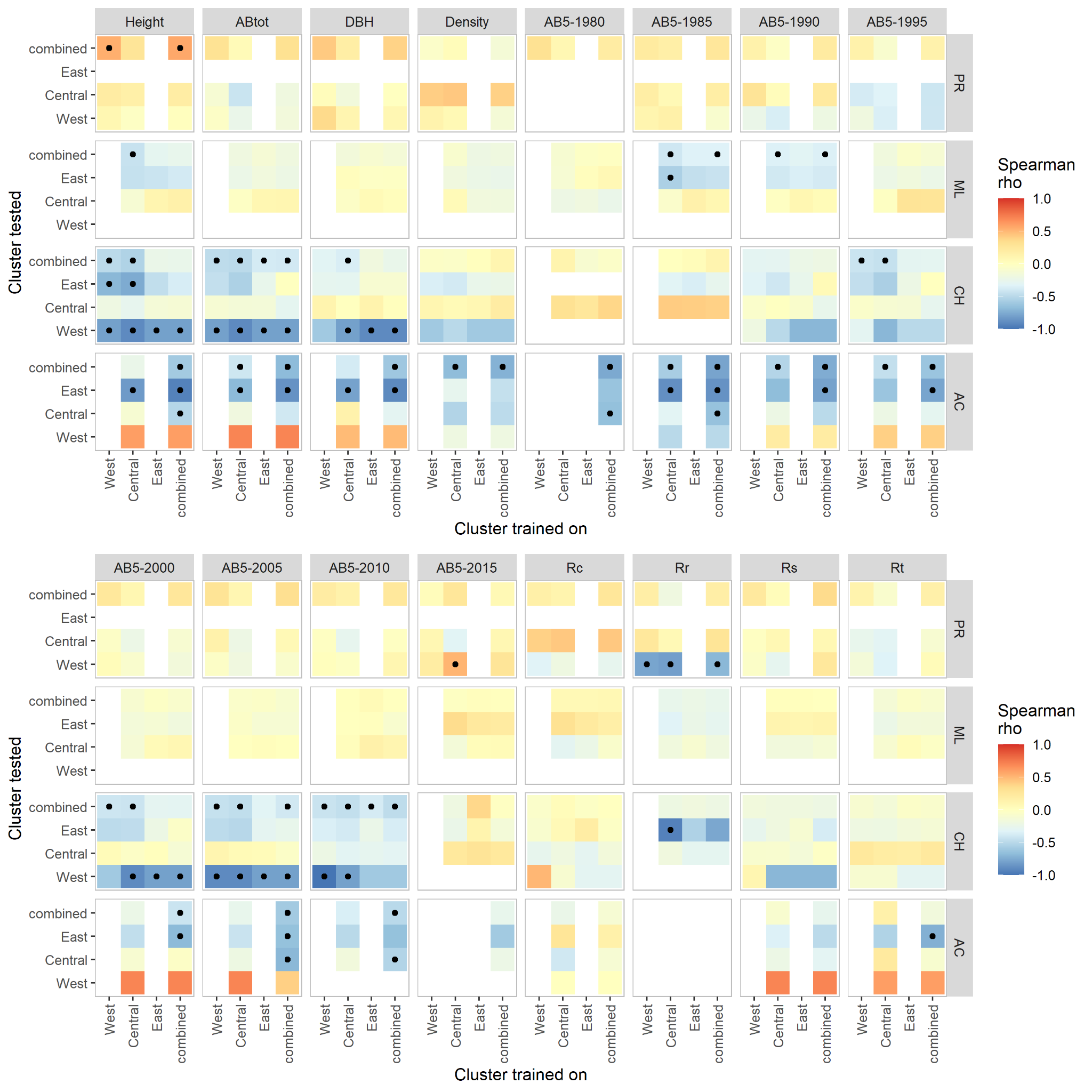

Supplementary Figure 17. RDA model performances when trained on individual genetic clusters or combined datasets and three climate PCs in predicting fitness traits of the same or other genetic clusters. Dots indicate correlations with confidence intervals not overlapping zero. Models were trained on a random set of 1000 markers and three climate PCs.

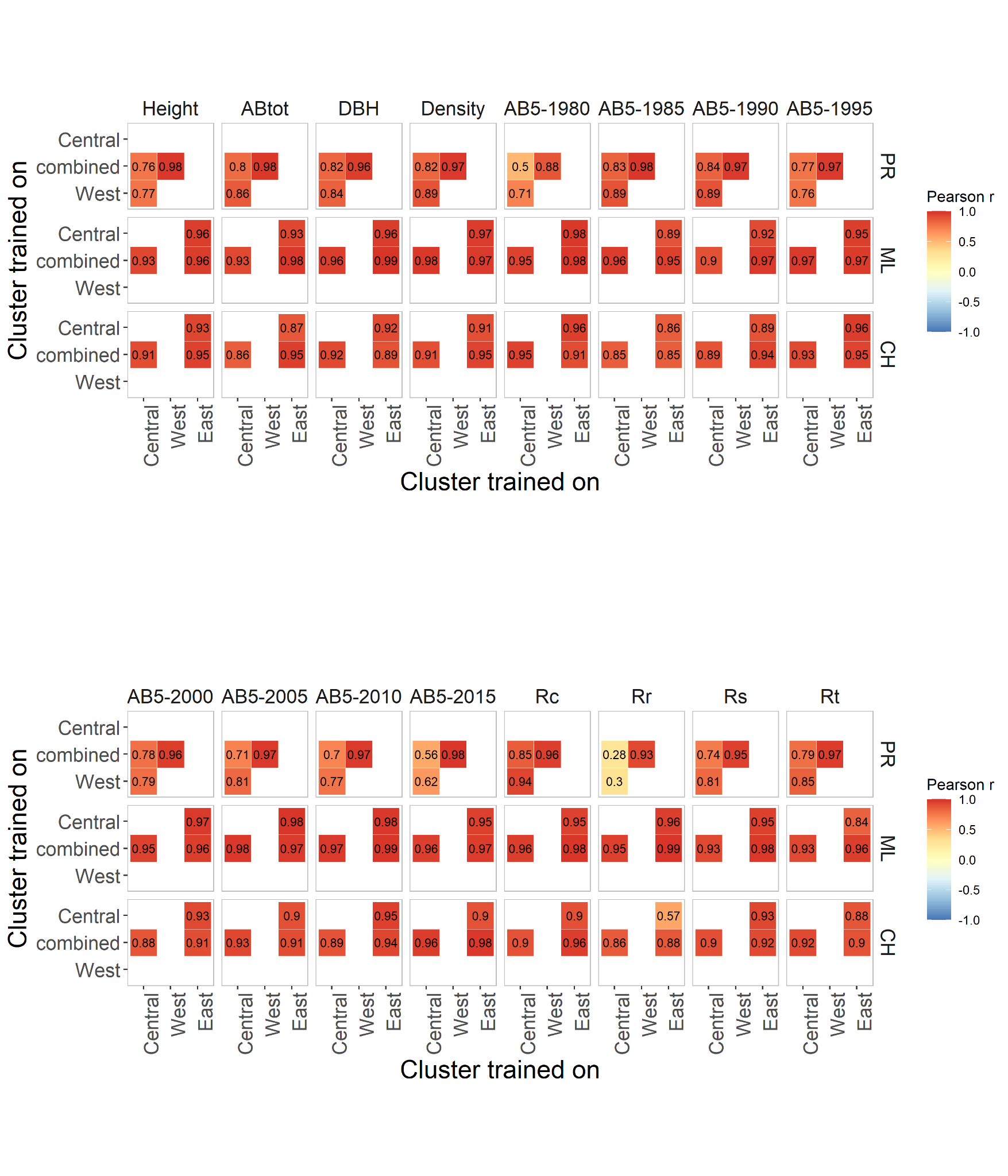

Supplementary Figure 18. Matrices of pairwise correlations between genomic offsets estimated using Gradient Forest models trained on single genetic clusters or combined datasets and three climate PCs. Models were trained on a random set of 1000 markers.

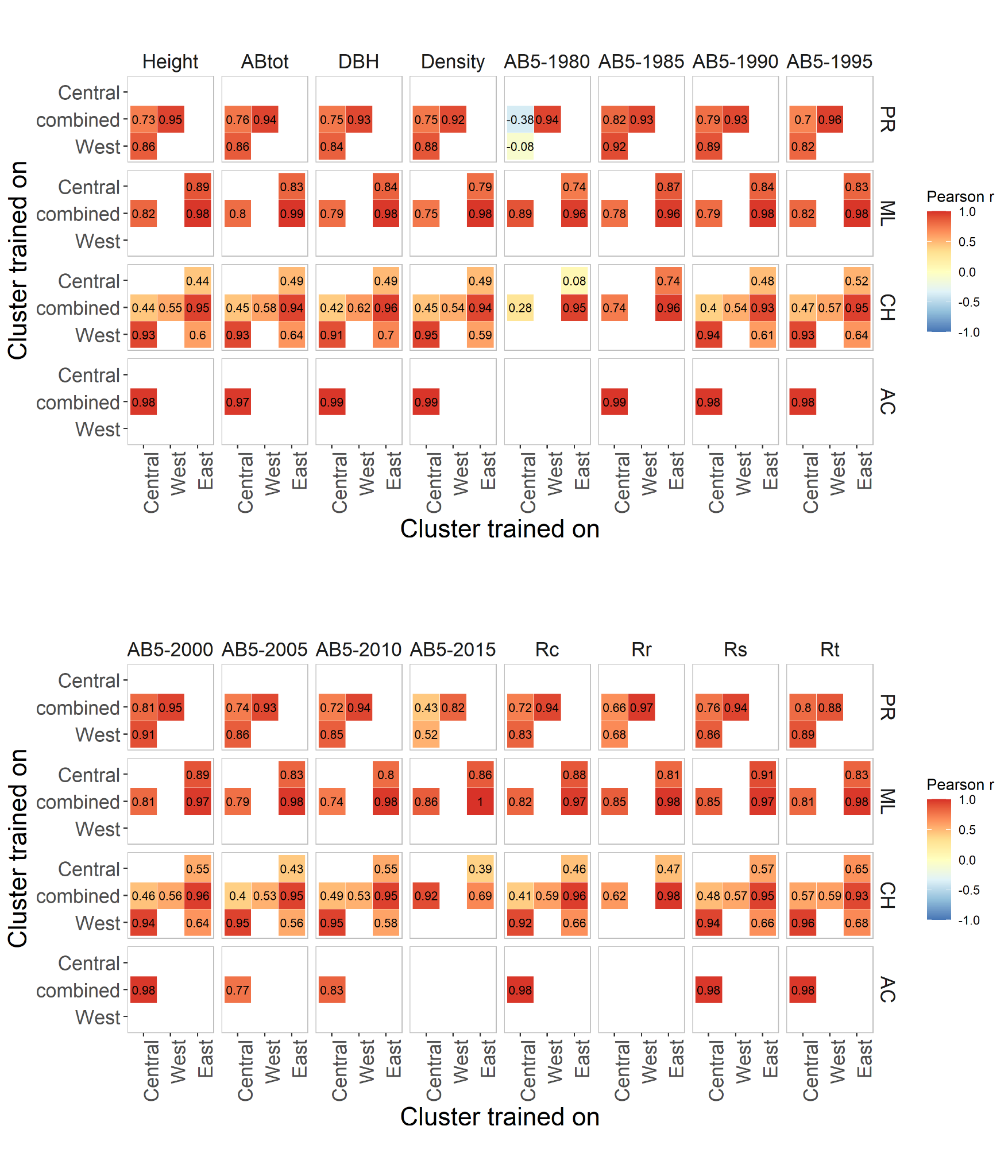

Supplementary Figure 19. Matrices of pairwise correlations between genomic offsets estimated using RDA models trained on single genetic clusters or combined datasets and three climate PCs. Models were trained on a random set of 1000 markers.

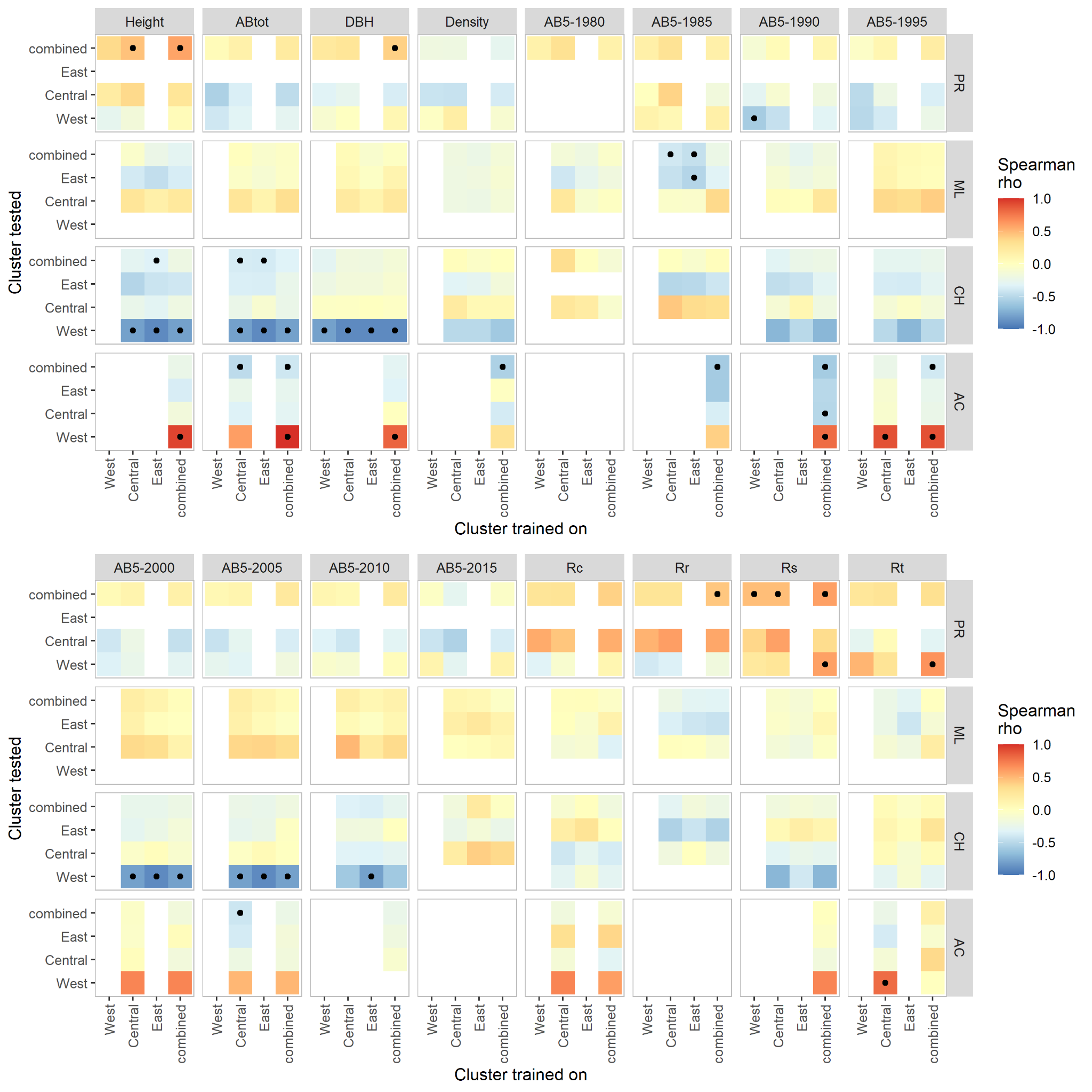

Supplementary Figure 20. Gradient Forest model performances when trained on individual genetic clusters or combined datasets and three selected climate variables in predicting fitness traits of the same or other genetic clusters. Dots indicate correlations with confidence intervals not overlapping zero. Models were trained on a random set of 1000 markers.

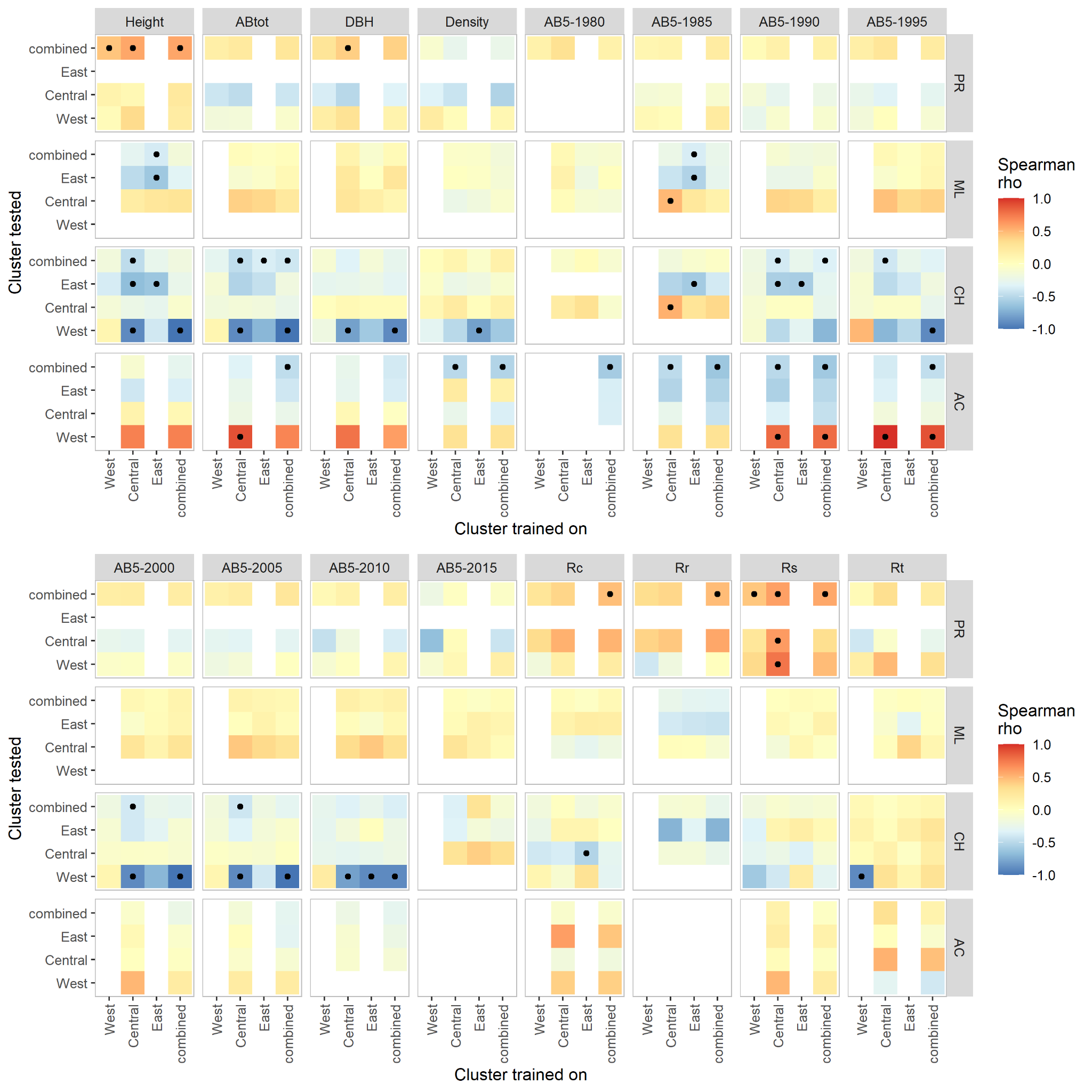

Supplementary Figure 21. RDA model performances when trained on individual genetic clusters or combined datasets and three selected climate variables in predicting fitness traits of the same or other genetic clusters. Dots indicate correlations with confidence intervals not overlapping zero. Models were trained on a random set of 1000 markers.

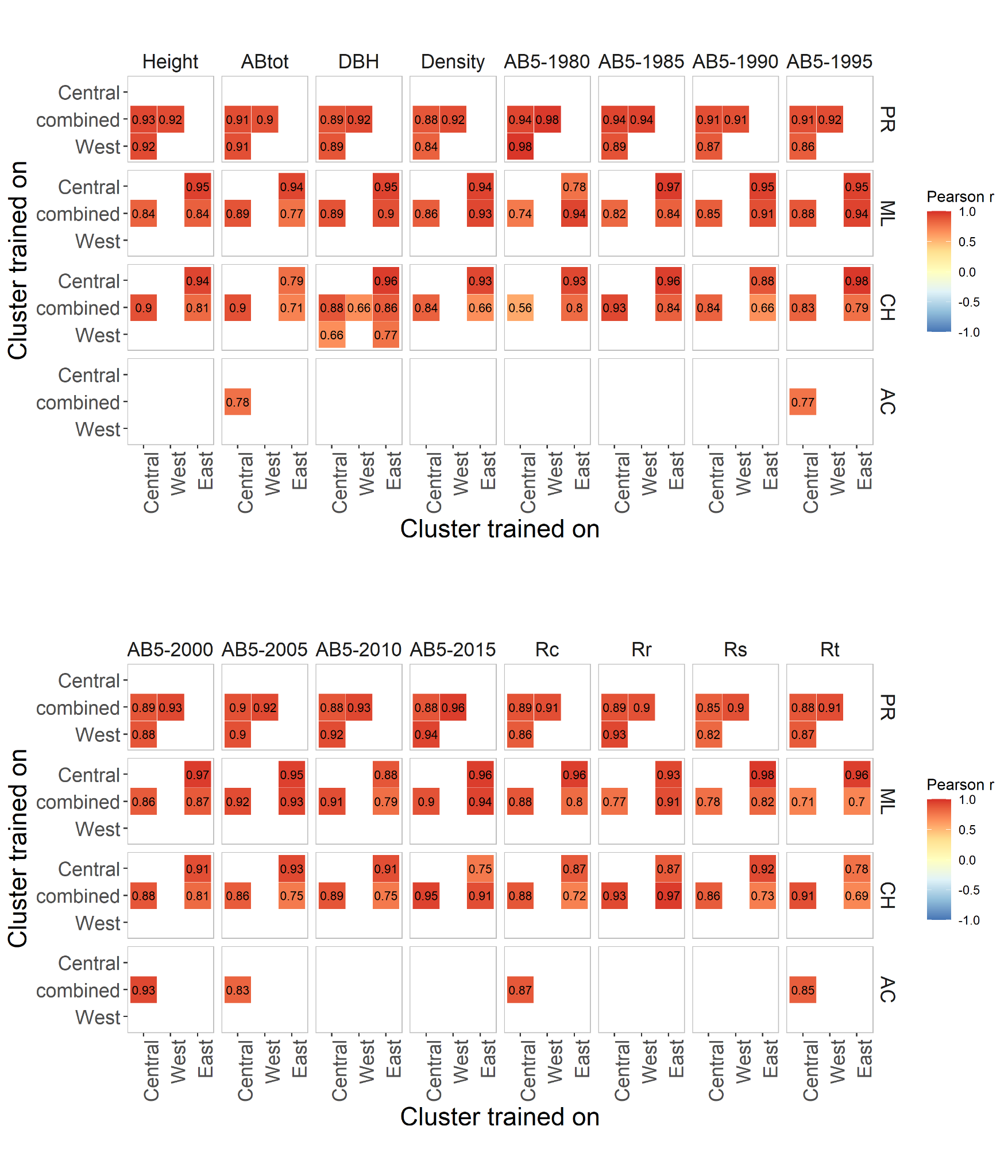
Supplementary Figure 22. Matrices of pairwise correlations between genomic offsets estimated using Gradient Forest models trained on single genetic clusters or combined datasets and three selected climate variables. Models were trained on a random set of 1000 markers.

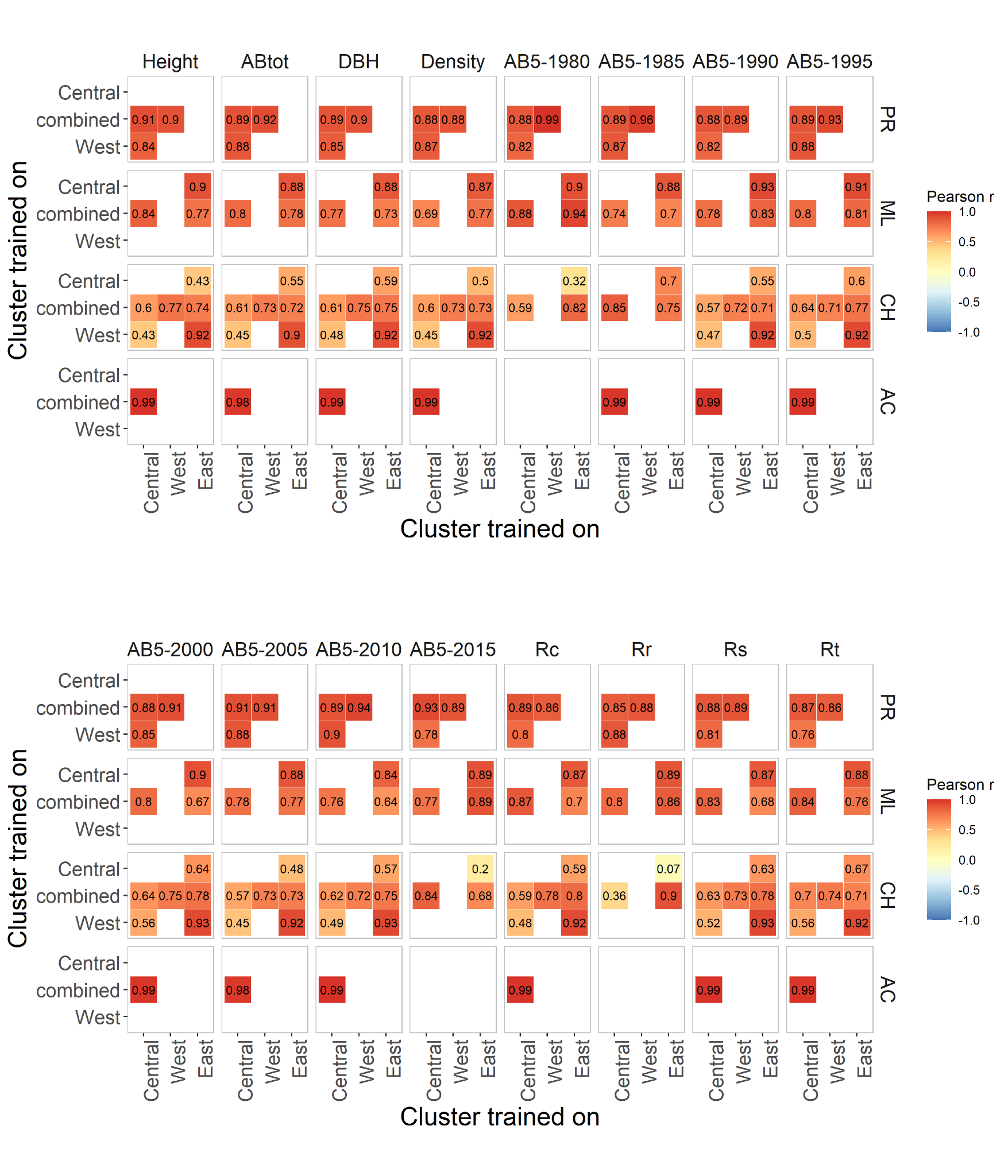

Supplementary Figure 23. Matrices of pairwise correlations between genomic offsets estimated using RDA models trained on single genetic clusters or combined datasets and three selected climate variables. Models were trained on a random set of 1000 markers.

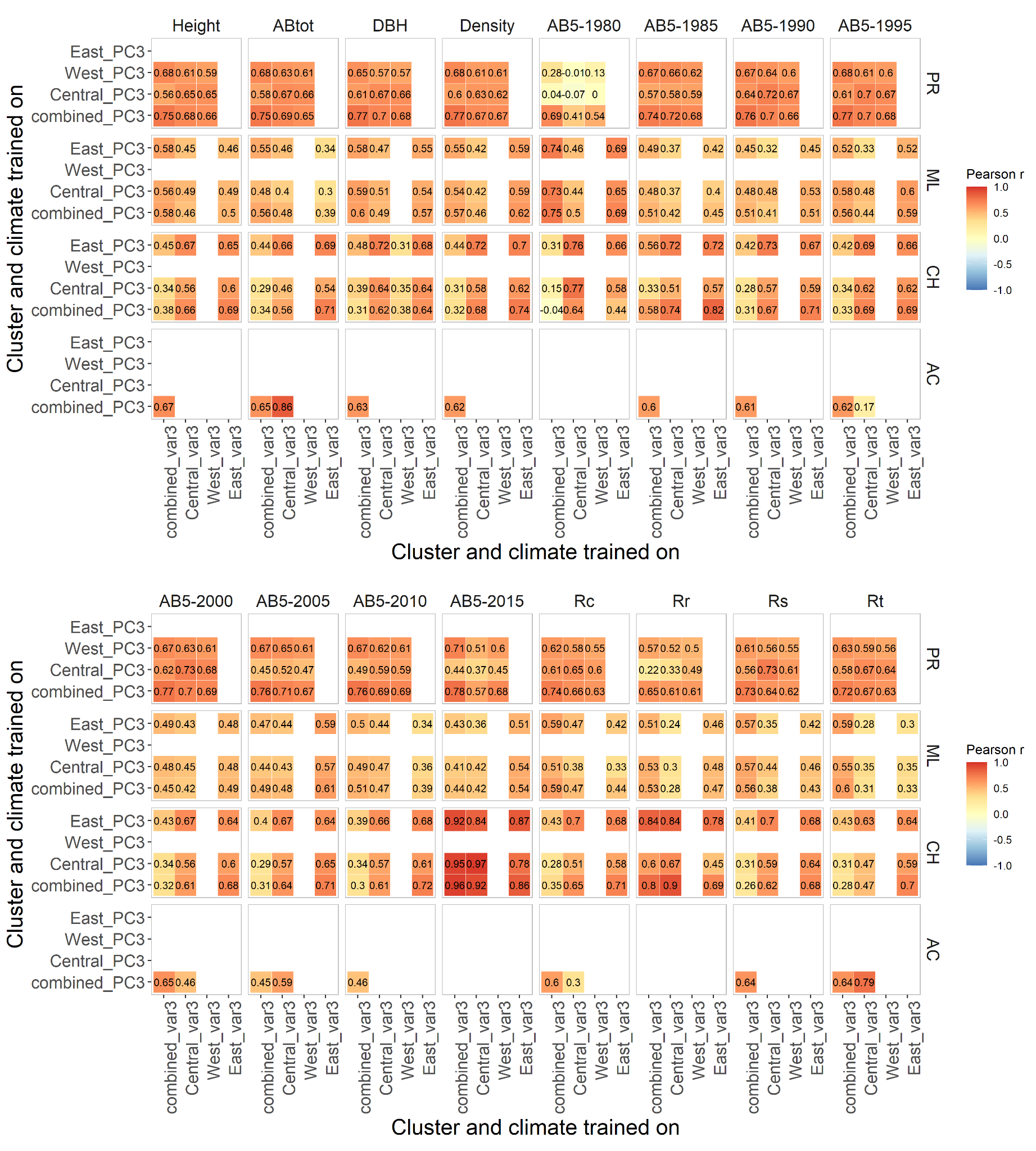

Supplementary Figure 24. Matrices of pairwise correlations between genomic offsets estimated using Gradient Forest models trained on two different climate datasets: a selected subset of three climate variables (“var3”, axis x) and three climate PCs (“PC3”, axis y). In this case models were trained on either complete datasets (“combined”) or individual genetic clusters. Models were trained on a random set of 1000 markers.

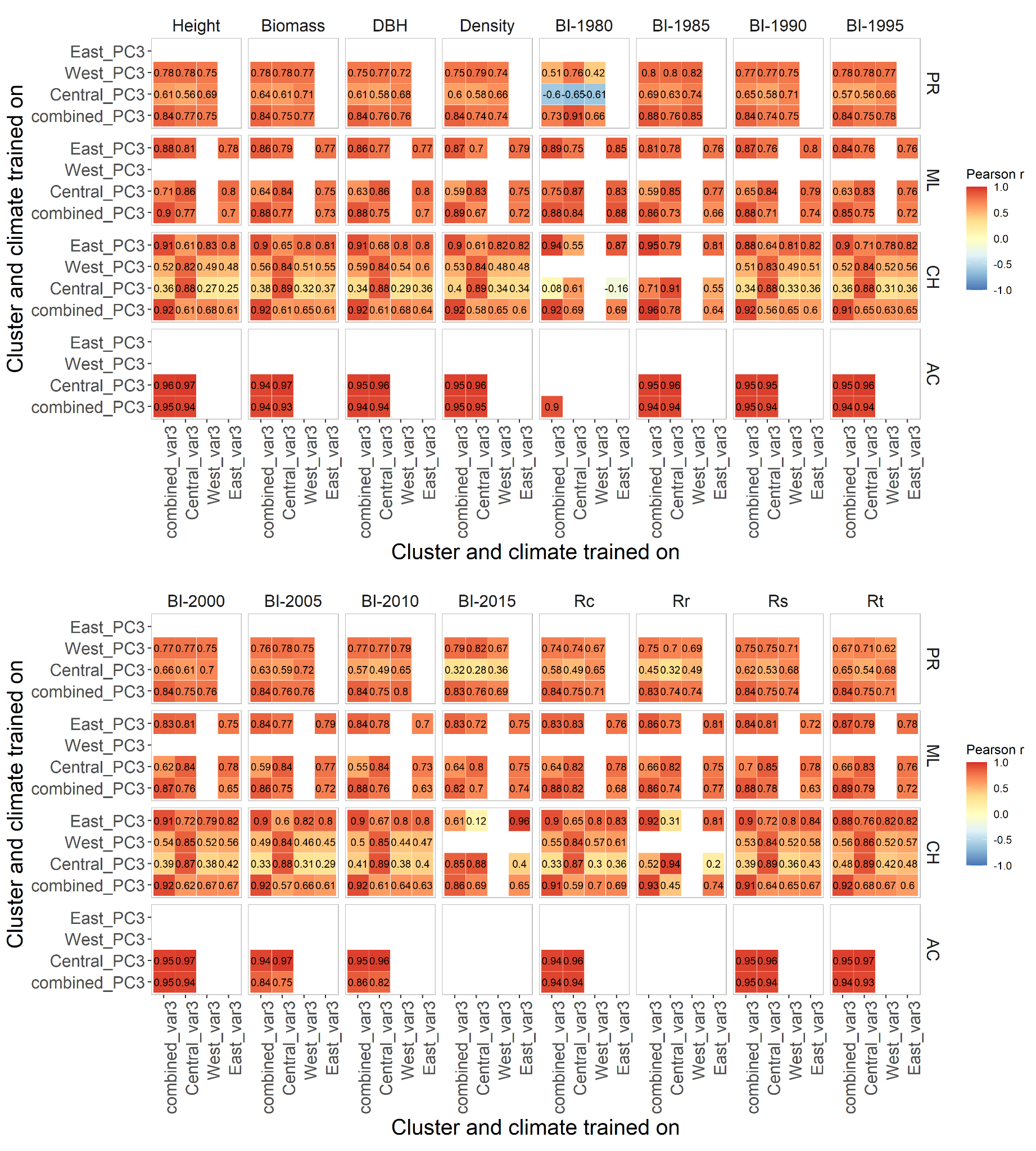

Supplementary Figure 25. Matrices of pairwise correlations between genomic offsets estimated using RDA models trained on two different climate datasets: a selected subset of three climate variables (“var3”, axis x) and three climate PCs (“PC3”, axis y). In this case models were trained on either complete datasets (“combined”) or individual genetic clusters. Models were trained on a random set of 1000 markers.

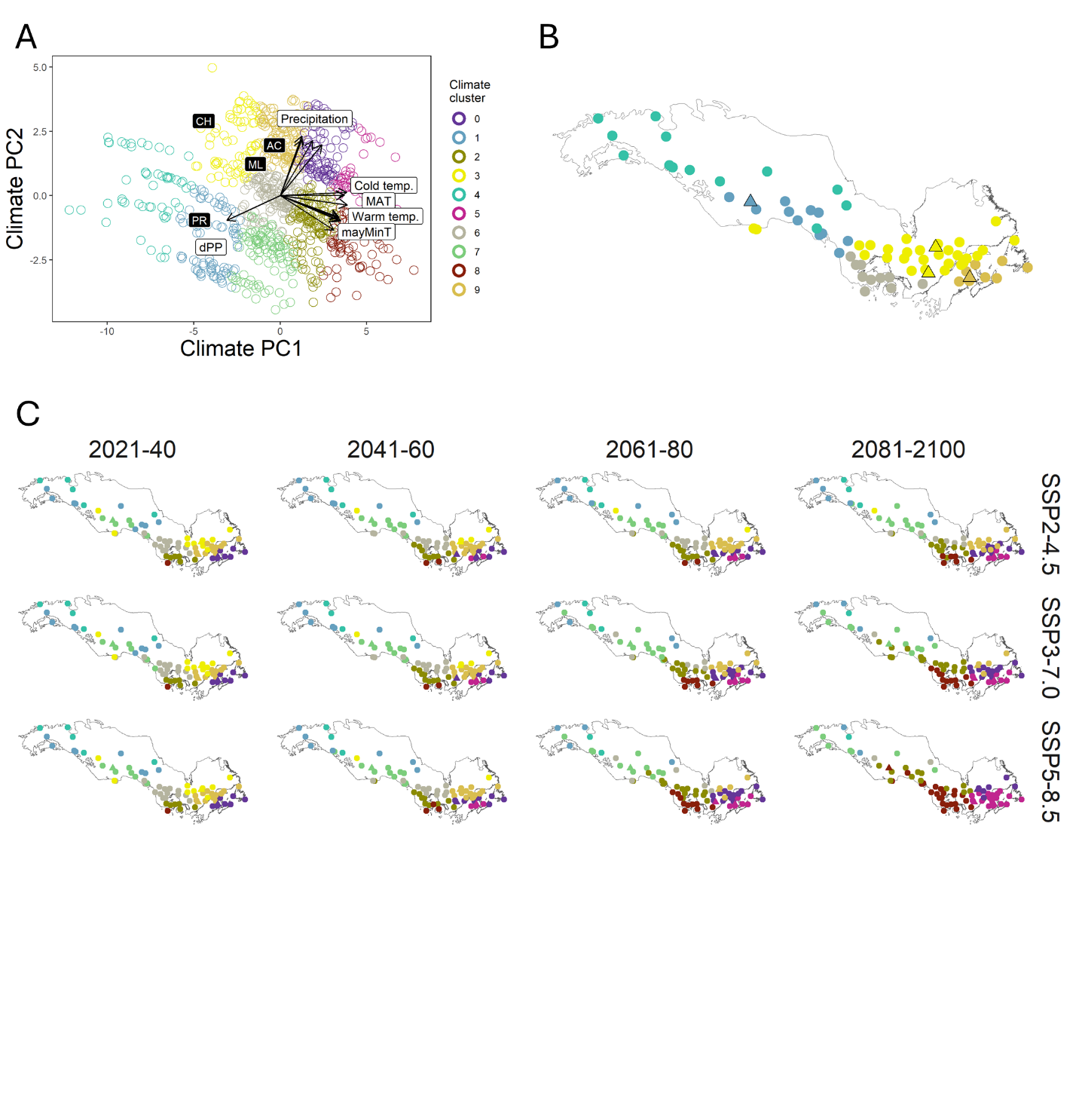

Supplementary Figure 26. Change of climate across the species range in the current century. A. Ordination of all population climate states (current and future projections according to three emission scenarios) along the first two principal components derived from 15 climate variables (arrows). On the plot climate variables are grouped by the type of variable, including variables linked to precipitation, variables describing low (Cold temp.) or high (Warm temp.) temperature regimes, mean annual temperature (MAT), minimum temperature in May (mayMinT) and photoperiod (dPP). B. Identified climate clusters present in the recent past (1961-1990) mapped onto species distribution. C. Predicted distribution of climate clusters in the four 20-year periods (columns) between 2021 and 2100 according to three emission scenarios (rows).

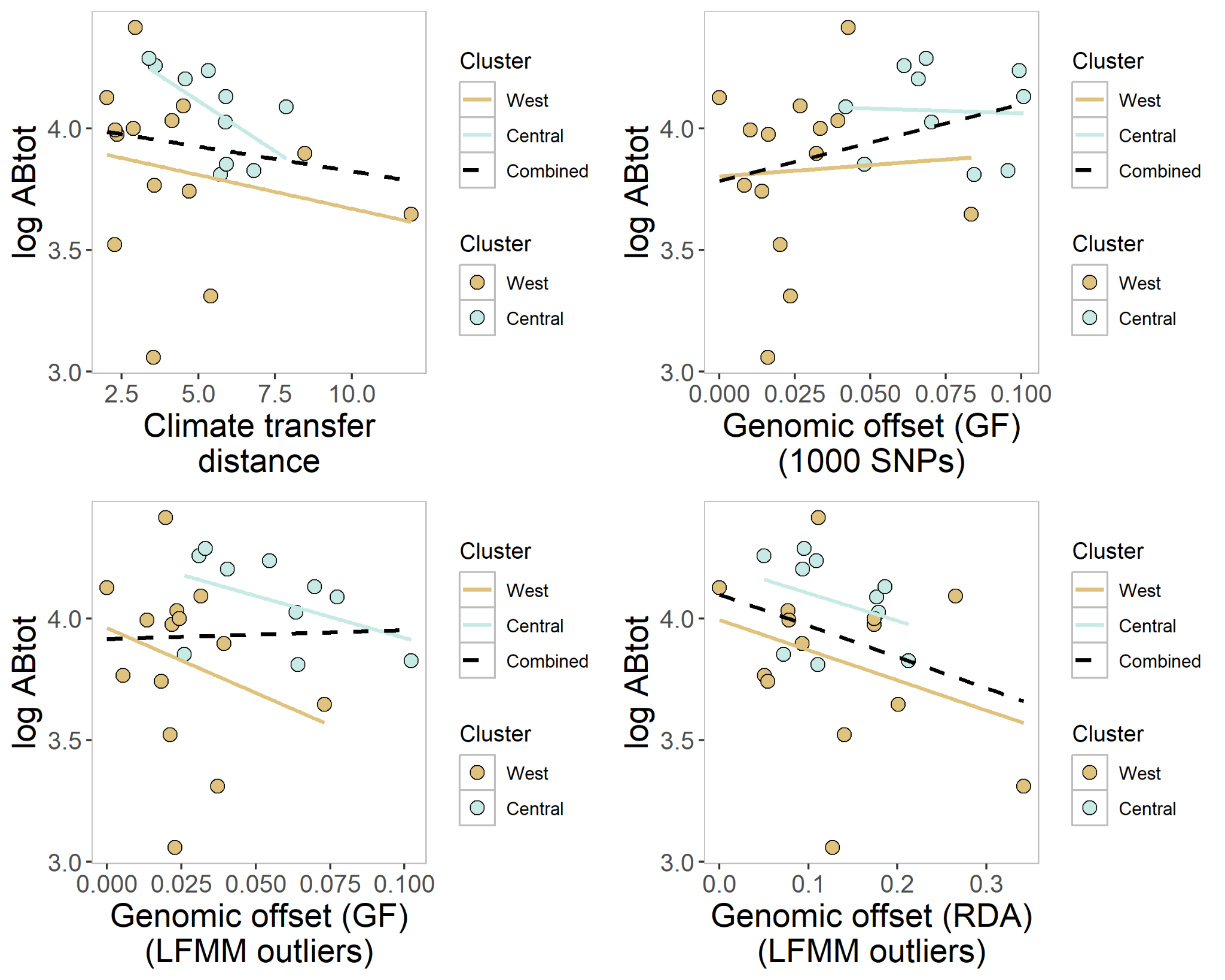

Supplementary Figure 27. Example of a confounding effect of differences in mean biomass increment between clusters on genomic offset estimates. Colored lines show linear models fitted separately to each cluster; dashed lines show linear models fitted to all samples. GF stands for Gradient Forest model and RDA for redundancy analysis prediction model that were used for calculating genomic offsets.

Supplementary Table 1. Geographical location of all initial black spruce populations (provenances) and the number of retained tree genotypes in each common garden after filtering.

| **Population** | **Region** | **Latitude** | **Longitude** | **AC** | **CH** | **ML** | **PR** | **VL** | **Total** |
| --- | --- | --- | --- | --- | --- | --- | --- | --- | --- |
| 4729 | A | 67.000 | -151.520 | 0 | 0 | 0 | 0 | 0 | 0 |
| 4420 | A | 64.700 | -148.300 | 0 | 0 | 0 | 7 | 1 | 8 |
| 7003 | A | 62.100 | -145.700 | 0 | 0 | 0 | 0 | 3 | 3 |
| 6998 | A | 67.250 | -138.330 | 0 | 0 | 0 | 0 | 0 | 0 |
| 7000 | B | 64.567 | -135.917 | 2 | 14 | 0 | 0 | 0 | 16 |
| 995 | B | 60.800 | -135.180 | 0 | 0 | 0 | 0 | 1 | 1 |
| 6994 | B | 60.533 | -134.450 | 3 | 0 | 0 | 0 | 0 | 3 |
| 2209 | B | 60.170 | -130.520 | 0 | 0 | 0 | 8 | 0 | 8 |
| 6988 | C | 58.700 | -123.600 | 6 | 0 | 0 | 19 | 0 | 25 |
| 6986 | C | 56.617 | -121.467 | 0 | 10 | 0 | 21 | 0 | 31 |
| 6980 | C | 52.500 | -115.880 | 0 | 0 | 0 | 13 | 0 | 13 |
| 6983 | C | 55.000 | -115.300 | 11 | 0 | 0 | 12 | 0 | 23 |
| 6979 | C | 52.367 | -115.250 | 0 | 6 | 18 | 13 | 0 | 37 |
| 6999 | C | 60.000 | -112.800 | 3 | 0 | 0 | 11 | 0 | 14 |
| 6970 | D | 56.050 | -108.700 | 0 | 11 | 0 | 13 | 0 | 24 |
| 6971 | D | 54.700 | -107.800 | 0 | 0 | 0 | 12 | 0 | 12 |
| 6972 | D | 53.530 | -105.770 | 0 | 0 | 0 | 15 | 4 | 19 |
| 6969 | D | 54.867 | -102.800 | 7 | 0 | 0 | 0 | 0 | 7 |
| 6965 | D | 52.483 | -101.433 | 0 | 12 | 6 | 12 | 0 | 30 |
| 6968 | D | 54.583 | -101.000 | 0 | 0 | 14 | 11 | 0 | 25 |
| 6963 | D | 51.630 | -100.780 | 0 | 0 | 0 | 15 | 0 | 15 |
| 6964 | D | 51.800 | -100.200 | 0 | 0 | 0 | 8 | 0 | 8 |
| 6977 | E | 57.600 | -96.700 | 0 | 0 | 0 | 12 | 0 | 12 |
| 6973 | E | 49.283 | -96.300 | 0 | 13 | 15 | 0 | 0 | 28 |
| 6967 | E | 55.500 | -94.667 | 0 | 15 | 0 | 11 | 0 | 26 |
| 6927 | E | 50.833 | -94.283 | 0 | 12 | 15 | 8 | 0 | 35 |
| 6930 | E | 48.800 | -93.667 | 0 | 15 | 10 | 0 | 0 | 25 |
| 6956 | E | 47.700 | -92.470 | 0 | 0 | 0 | 8 | 0 | 8 |
| 6922 | E | 50.250 | -91.667 | 0 | 11 | 17 | 0 | 0 | 28 |
| 4353 | E | 47.700 | -91.300 | 0 | 9 | 12 | 0 | 0 | 21 |
| 4344 | E | 46.117 | -90.933 | 0 | 12 | 18 | 0 | 0 | 30 |
| 6917 | E | 49.000 | -90.450 | 0 | 0 | 15 | 0 | 0 | 15 |
| 3268 | E | 44.217 | -90.367 | 0 | 8 | 13 | 0 | 0 | 21 |
| 6916 | E | 50.300 | -89.180 | 0 | 0 | 0 | 14 | 0 | 14 |
| 4277 | E | 45.733 | -88.983 | 7 | 14 | 14 | 0 | 0 | 35 |

Supplementary Table 1. Continued

| **Population** | **Region** | **Latitude** | **Longitude** | **AC** | **CH** | **ML** | **PR** | **VL** | **Total** |
| --- | --- | --- | --- | --- | --- | --- | --- | --- | --- |
| 6944 | F | 45.983 | -86.850 | 0 | 0 | 0 | 0 | 0 | 0 |
| 6914 | F | 48.633 | -85.333 | 0 | 9 | 17 | 0 | 0 | 26 |
| 6909 | F | 49.750 | -85.083 | 0 | 0 | 16 | 0 | 0 | 16 |
| 4351 | F | 46.050 | -84.783 | 10 | 9 | 18 | 12 | 0 | 49 |
| 6941 | F | 44.633 | -84.333 | 13 | 0 | 0 | 0 | 0 | 13 |
| 6907 | F | 48.533 | -81.417 | 0 | 15 | 18 | 11 | 0 | 44 |
| 6937 | F | 51.100 | -80.867 | 0 | 0 | 0 | 0 | 0 | 0 |
| 6938 | F | 49.350 | -80.750 | 0 | 0 | 14 | 0 | 0 | 14 |
| 6905 | G | 46.920 | -79.700 | 0 | 0 | 0 | 7 | 0 | 7 |
| 333 | G | 49.617 | -77.750 | 10 | 15 | 14 | 0 | 0 | 39 |
| 6901 | G | 45.167 | -77.167 | 0 | 10 | 17 | 0 | 0 | 27 |
| 336 | G | 48.367 | -76.950 | 0 | 6 | 18 | 0 | 0 | 24 |
| 338 | G | 47.083 | -76.550 | 7 | 14 | 17 | 0 | 0 | 38 |
| 325 | G | 50.450 | -73.633 | 0 | 8 | 16 | 7 | 0 | 31 |
| 326 | G | 49.033 | -73.450 | 3 | 7 | 13 | 0 | 0 | 23 |
| 321 | G | 46.933 | -72.100 | 0 | 15 | 13 | 0 | 0 | 28 |
| 352 | G | 49.600 | -71.300 | 2 | 11 | 16 | 0 | 0 | 29 |
| 329 | G | 47.867 | -71.200 | 0 | 0 | 15 | 0 | 0 | 15 |
| 345 | H | 48.933 | -69.133 | 0 | 0 | 17 | 0 | 0 | 17 |
| 4360 | H | 45.500 | -68.800 | 4 | 14 | 17 | 0 | 0 | 35 |
| 342 | H | 50.667 | -68.767 | 0 | 12 | 14 | 0 | 0 | 26 |
| 1534 | H | 47.700 | -68.317 | 2 | 13 | 16 | 0 | 0 | 31 |
| 6808 | H | 47.920 | -67.820 | 0 | 0 | 0 | 13 | 0 | 13 |
| 1528 | H | 46.833 | -67.167 | 0 | 0 | 10 | 0 | 0 | 10 |
| 332 | H | 48.500 | -67.117 | 7 | 13 | 17 | 0 | 0 | 37 |
| 1531 | H | 45.583 | -66.483 | 0 | 13 | 17 | 0 | 0 | 30 |
| 1538 | H | 47.750 | -65.117 | 0 | 12 | 14 | 0 | 0 | 26 |
| 369 | H | 48.883 | -64.650 | 0 | 0 | 14 | 0 | 0 | 14 |
| 355 | H | 49.633 | -63.367 | 6 | 11 | 18 | 0 | 0 | 35 |
| 1329 | H | 46.050 | -62.850 | 2 | 7 | 17 | 0 | 0 | 26 |
| 6805 | I | 53.417 | -60.383 | 0 | 7 | 17 | 11 | 0 | 35 |
| 1530 | I | 45.933 | -60.167 | 0 | 6 | 14 | 0 | 0 | 20 |
| 6802 | I | 48.217 | -58.917 | 0 | 13 | 8 | 0 | 0 | 21 |
| 6804 | I | 50.900 | -56.100 | 0 | 11 | 11 | 0 | 0 | 22 |
| 6801 | I | 47.333 | -53.117 | 0 | 13 | 13 | 0 | 0 | 26 |

Supplementary Table 2. Geographical location and mean climate conditions of common gardens.

| **Common garden** | **Code** | **Lat** | **Lon** | **Alt** | **MAT** | **TP** | **CMI** | **GDD** |
| --- | --- | --- | --- | --- | --- | --- | --- | --- |
| Acadia (AC) | E60-A | 46.01 | -66.39 | 108 | 4.34 | 1201 | 69.84 | 2633 |
| Peace River (PR) | G348 | 56.0 | -116.66 | 735 | 0.78 | 479 | 8.23 | 2102 |
| Mont-Laurier (ML) | E353-B1 | 46.60 | -75.84 | 236 | 3.31 | 956 | 47.25 | 2630 |
| Chibougamau (CH) | E353-B3 | 50.02 | -74.21 | 408 | -0.87 | 969 | 62.37 | 1967 |
| Valcartier (VL) | E353-B4 | 46.86 | -71.52 | 172 | 3.34 | 1278 | 83.12 | 2575 |

Lon – longitude

Lat – latitude

Alt – altitude (m)

MAT – mean annual temperature in ºC

TP – mean total annual precipitation in mm

CMI – mean annual climate moisture index

GDD – degree-days at 0ºC at the last day of the year

Supplementary Table 3. List of climate variables.

| **Abbreviation** | **Description** |
| --- | --- |
| MAT | mean annual temperature |
| MMinT | mean of monthly minimum temperatures |
| MMaxT | mean of monthly maximum temperatures |
| MWMT | mean temperature of the warmest month |
| MCMT | mean temperature of the coldest month |
| WMMAX | maximum temperature of the warmest month |
| CMMIN | minimum temperature of the coldest month |
| TAR | temperature annual range |
| sumMT | mean summer (June to August) temperature |
| winMT | mean winter (December to February) temperature |
| fallMT | mean fall (September to November) temperature |
| sprMT | mean spring (March to May) temperature |
| mayMinT | minimum May temperature |
| dGDD0 | day of the year when degree-days at 0ºC reaches 300 |
| GDD0 | degree-days at 0ºC at the last day of the year |
| GDD5 | degree-days at 5ºC at the last day of the year |
| dPP | day of the year when photoperiod reaches 10h |
| sprFD | number of frost days in spring |
| fallFD | number of frost days in fall |
| sprFP0 | day when spring frost probability at 0ºC falls below 10% |
| fallFP0 | day when fall frost probability at 0ºC exceeds 10% |
| sprFP2 | day when spring frost probability at -2ºC falls below 10% |
| fallFP2 | day when fall frost probability at -2ºC exceeds 10% |
| sprFP4 | day when spring frost probability at -4ºC falls below 10% |
| fallFP4 | day when fall frost probability at -4ºC exceeds 10% |
| TP | total precipitation sum |
| WSTP | total precipitation sum from May to September |
| sumTP | total precipitation sum from June to August |
| winSP | snowpack accumulation from December to February |
| maySP | snowpack accumulation in May |
| CMI | climate moisture index |
| sumCMI | summer climate moisture index |
| PET | potential evapotranspiration |
| sumPET | potential evapotranspiration from June to August |
| SPEI | standardized precipitation-evapotranspiration index |
| sumSPEI | summer standardized precipitation-evapotranspiration index |

Supplementary Table 3. Continued.

| **Abbreviation** | **Description** |
| --- | --- |
| RH | mean annual relative humidity |
| sumRH | mean relative humidity from June to August |
| SR | mean annual solar radiation |
| sumSR | mean summer (June-August) solar radiation |
| sumSMI | mean summer (June to August) soil moisture index |
| absMin | absolute temperature minimum |
| absMax | absolute temperature maximum |

Supplementary Table 4. Number of populations assigned to TEST and training (TRAIN) datasets. Datasets with less than 5 trees per populations were excluded from downstream analyses.

|  | **AC** | | **CH** | | **ML** | | **PR** | | **Combined** | |
| --- | --- | --- | --- | --- | --- | --- | --- | --- | --- | --- |
| **Trait name** | **TEST** | **TRAIN** | **TEST** | **TRAIN** | **TEST** | **TRAIN** | **TEST** | **TRAIN** | **TEST** | **TRAIN** |
| **Wood density** | 32 | 10 | 37 | 37 | 40 | 42 | 26 | 26 | 135 | 115 |
| **ABtot** | 32 | 10 | 37 | 37 | 40 | 42 | 26 | 26 | 135 | 115 |
| **AB5 1980** | 27 | 5 | 21 | 24 | 32 | 35 | 16 | 23 | 96 | 87 |
| **AB5 1985** | 32 | 10 | 35 | 35 | 40 | 42 | 23 | 23 | 130 | 110 |
| **AB5 1990** | 32 | 10 | 37 | 37 | 39 | 41 | 26 | 26 | 134 | 114 |
| **AB5 1995** | 32 | 10 | 37 | 37 | 40 | 42 | 26 | 26 | 135 | 115 |
| **AB5 2000** | 32 | 10 | 37 | 37 | 40 | 42 | 26 | 26 | 135 | 115 |
| **AB5 2005** | 31 | 9 | 37 | 37 | 40 | 42 | 26 | 26 | 134 | 114 |
| **AB5 2010** | 30 | 8 | 37 | 37 | 40 | 42 | 26 | 26 | 133 | 113 |
| **AB5 2015** | 27 | 4 | 20 | 19 | 38 | 40 | 20 | 20 | 105 | 83 |
| **DBH** | 33 | 10 | 37 | 37 | 42 | 42 | 26 | 26 | 138 | 115 |
| **Height** | 33 | 10 | 37 | 37 | 42 | 42 | 26 | 26 | 138 | 115 |
| **Rc** | 32 | 10 | 37 | 37 | 40 | 42 | 23 | 26 | 132 | 115 |
| **Rs** | 32 | 10 | 37 | 37 | 40 | 42 | 23 | 26 | 132 | 115 |
| **Rr** | 29 | 2 | 17 | 16 | 37 | 40 | 21 | 24 | 104 | 82 |
| **Rt** | 32 | 10 | 37 | 37 | 40 | 42 | 23 | 26 | 132 | 115 |

Supplementary Table 5. Number of genotypes assigned to TEST and training (TRAIN) datasets.

|  | **AC** | | **CH** | | **ML** | | **PR** | | **Combined** | |
| --- | --- | --- | --- | --- | --- | --- | --- | --- | --- | --- |
| **Trait name** | **TEST** | **TRAIN** | **TEST** | **TRAIN** | **TEST** | **TRAIN** | **TEST** | **TRAIN** | **TEST** | **TRAIN** |
| **Wood density** | 348 | 84 | 151 | 375 | 149 | 519 | 111 | 288 | 759 | 1266 |
| **ABtot** | 348 | 84 | 151 | 375 | 149 | 519 | 111 | 288 | 759 | 1266 |
| **AB5 1980** | 276 | 34 | 58 | 159 | 100 | 297 | 32 | 193 | 466 | 683 |
| **AB5 1985** | 348 | 84 | 140 | 313 | 148 | 510 | 90 | 207 | 726 | 1114 |
| **AB5 1990** | 348 | 84 | 151 | 375 | 138 | 514 | 111 | 277 | 748 | 1250 |
| **AB5 1995** | 348 | 84 | 151 | 375 | 149 | 519 | 111 | 288 | 759 | 1266 |
| **AB5 2000** | 348 | 84 | 151 | 375 | 149 | 519 | 111 | 288 | 759 | 1266 |
| **AB5 2005** | 341 | 78 | 151 | 375 | 149 | 519 | 111 | 287 | 752 | 1259 |
| **AB5 2010** | 332 | 66 | 151 | 368 | 149 | 518 | 111 | 286 | 743 | 1238 |
| **AB5 2015** | 310 | 28 | 82 | 131 | 129 | 369 | 74 | 166 | 595 | 694 |
| **DBH** | 351 | 84 | 141 | 369 | 156 | 515 | 113 | 289 | 761 | 1257 |
| **Height** | 351 | 84 | 141 | 369 | 156 | 515 | 113 | 289 | 761 | 1257 |
| **Rc** | 342 | 84 | 151 | 375 | 149 | 519 | 100 | 289 | 742 | 1267 |
| **Rs** | 342 | 84 | 151 | 375 | 149 | 519 | 101 | 289 | 743 | 1267 |
| **Rr** | 333 | 14 | 66 | 107 | 128 | 323 | 90 | 221 | 617 | 665 |
| **Rt** | 342 | 84 | 151 | 375 | 149 | 519 | 101 | 289 | 743 | 1267 |

Supplementary Table 6. Results of AMOVA.

|  | **Df** | **Sum of squares** | **Expected mean squares** | **Sigma** | **Percent variance** |
| --- | --- | --- | --- | --- | --- |
| Between clusters | 4 | 333655.7 | 83413.93 | 155.11 | 4.6 |
| Between populations within clusters | 58 | 436945.3 | 7533.54 | 74.82 | 2.2 |
| Between samples within populations | 1397 | 5749935 | 4115.917 | 1010.76 | 30.3 |
| Within samples | 1460 | 3057816 | 2094.394 | 2094.39 | 62.8 |
| Total | 2919 | 9578352 | 3281.381 | 3335.09 | 100 |

Supplementary Table 7. Results of partial RDA analysis. Different classes of variables include genetic structure (gen), climate variables (clim) and spatial autocorrelation axes (spa).

| **Model** | **adj *R*^2^** | **Variance** | **df** | **F** | ***P*-value** | **adj *P*** | **% adjusted variance** |
| --- | --- | --- | --- | --- | --- | --- | --- |
| gen + clim + spa | 0.413 | 164.3 | 8 | 6.45 | 0.001 | 0.005 | 100.0 |
| clim \| gen + spa | 0.011 | 12.8 | 3 | 1.35 | 0.001 | 0.005 | 2.6 |
| spa \| clim + gen | 0.007 | 11.8 | 3 | 1.24 | 0.001 | 0.005 | 1.8 |
| gen \| clim + spa | 0.015 | 10.8 | 2 | 1.70 | 0.001 | 0.005 | 3.6 |
| clim | 0.313 | 116.5 | 3 | 10.42 | 0.001 | 0.005 | 75.8 |

Supplementary Table 8. Partition of variance explained by climate in partial RDA models fitted to species-wide or within-cluster data. Models were run using climate as the only predictor (climate-only) or while controlling for spatial (MEM) and genetic variables (pure climate). Adj stands for adjusted. Negative adjusted *R*² values were set to 0.

| **Level** | **Climate-only model; adj *R*^2^ (adj. *P*)** | **Pure climate model; adj *R*^2^ (adj. *P*)** | **Shared with genetic and spatial factors (% of climate-only)** | **Shared with genetic and spatial factors (% of full model)** |
| --- | --- | --- | --- | --- |
| Species-wide | 0.313 (0.005) | 0.007 (0.005) | 0.306 (98%) | 74% |
| West | 0.096 (0.009) | 0.014 (0.25) | 0.082 (85%) | 31% |
| Central | 0.049 (0.015) | 0 (1) | 0.049 (~100%) | 38% |
| East | 0.037 (0.005) | 0.009 (0.302) | 0.028 (76%) | 32% |
